# Spatially resolved multiomics of human cardiac niches

**DOI:** 10.1101/2023.01.30.526202

**Authors:** Kazumasa Kanemaru, James Cranley, Daniele Muraro, Antonio M.A. Miranda, Jan Patrick Pett, Monika Litvinukova, Natsuhiko Kumasaka, Siew Yen Ho, Krzysztof Polanski, Laura Richardson, Lukas Mach, Monika Dabrowska, Nathan Richoz, Sam N. Barnett, Shani Perera, Anna Wilbrey-Clark, Carlos Talavera-López, Ilaria Mulas, Krishnaa T. Mahbubani, Liam Bolt, Lira Mamanova, Liz Tuck, Lu Wang, Margaret M. Huang, Martin Prete, Sophie Pritchard, John Dark, Kourosh Saeb-Parsy, Minal Patel, Menna R. Clatworthy, Norbert Hübner, Rasheda A. Chowdhury, Michela Noseda, Sarah A. Teichmann

## Abstract

A cell’s function is defined by its intrinsic characteristics and its niche: the tissue microenvironment in which it dwells. Here, we combine single-cell and spatial transcriptomic data to discover cellular niches within eight regions of the human heart. We map cells to micro-anatomic locations and integrate knowledge-based and unsupervised structural annotations. For the first time, we profile the cells of the human cardiac conduction system, revealing their distinctive repertoire of ion channels, G-protein coupled receptors, and cell interactions using a custom CellPhoneDB.org module. We show that the sinoatrial node is compartmentalised, with a core of pacemaker cells, fibroblasts and glial cells supporting paracrine glutamatergic signalling. We introduce a druggable target prediction tool, drug2cell, which leverages single-cell profiles and drug-target interactions, providing unexpected mechanistic insights into the chronotropic effects of drugs, including GLP-1 analogues. In the epicardium, we show enrichment of both IgG+ and IgA+ plasma cells forming immune niches which may contribute to infection defence. We define a ventricular myocardial-stress niche enriched for activated fibroblasts and stressed cardiomyocytes, cell states that are expanded in cardiomyopathies. Overall, we provide new clarity to cardiac electro-anatomy and immunology, and our suite of computational approaches can be deployed to other tissues and organs.

## Introduction

The heart is composed of distinct tissues which contain niches of specialised cell types conferring site-specific functionality. Single-cell and single-nuclei RNA sequencing (sc/snRNAseq) offers a powerful, unbiased framework to characterise these cells^1,2^. Our previous atlas illustrated the heart’s unique diversity, describing over 60 cell states^1^. Expanding these profiles with spatially-resolved transcriptomics across multiple tissues allows us to restore structural information lost in single-cell techniques and gain insight into collective function^3,4^ In particular, how diverse cell states are co-located, spatially organised, and interact.

The cardiac conduction system (CCS), responsible for the regular and coordinated electrical activation of the heart^5^, contains structures including the sinoatrial and atrioventricular nodes (SAN, AVN) and atrioventricular bundle (AVB) and His-Purkinje network which are each home to cells with unique electrophysiological properties^6,7^. We combined targeted dissection and histology to generate full-breadth transcriptomic profiles of human CCS cells for the first time. Furthermore, we used spatial transcriptomics to map these cells into their microanatomic locations and discovered their niche-partner cells. Inspired by the broad receptor expression profile of pacemaker (P) cells, we extended CellPhoneDB’s^8^ cell-cell interaction analysis with a new neural-GPCR module, which highlighted synaptic connections, including exciting new insights with neighbouring glial cells and glutamatergic signalling capability.

Off-target activity of non-cardiac therapies on the heart and its conduction system is a major reason for drug development failure and withdrawal^9^. Additionally, there is a major need to identify new druggable targets in cardiology^10^. Motivated by this we developed a pipeline, drug2cell, which integrates drug-target interactions from the ChEMBL database (https://www.ebi.ac.uk/chembl/) with user-provided single-cell data to comprehensively evaluate drug target expression in single cells. Applying this approach to P cells provided mechanistic insight into the chronotropic effects of non-cardiac medications.

Finally, we showed that our integrated multiomic approach is capable of niche (i.e. cellular tissue microenvironment) discovery. We discovered an epicardial immune defence system and a ventricular myocardial-stress niche, and inferred specific intercellular signalling active within each cellular microenvironment. These novel cardiac cellular niches allow us to refine the cellular components underlying the structural microanatomy of the human heart.

## Results

### Profiling cardiac tissues with multiple modalities

To identify cellular microenvironments in the adult human heart, we combined spatially resolved single-cell multiomic technologies with machine learning. We integrated previously published sc/snRNA-seq data^1^ with newly generated multiome data (paired snRNAseq and snATACseq), and spatial transcriptomics data (10X Genomics). We studied eight anatomical regions including left and right ventricular free walls (LV and RV), left and right atria (LA and RA), left ventricular apex (AX), interventricular septum (SP), sino-atrial node (SAN) and atrioventricular node (AVN)(**Fig. 1A**). In total, our data include 22 donors ranging from 40 to 75 years old (**Fig. 1A**). All tissue was from transplant donors without a history of cardiac disease or arrhythmia (**Supp. Table 1**) and hearts contributing to the SAN and AVN regions were from donors with normal conduction parameters confirmed on 12 lead electrocardiogram prior to donation (**Supp. Table 1**).

**Fig. 1:**
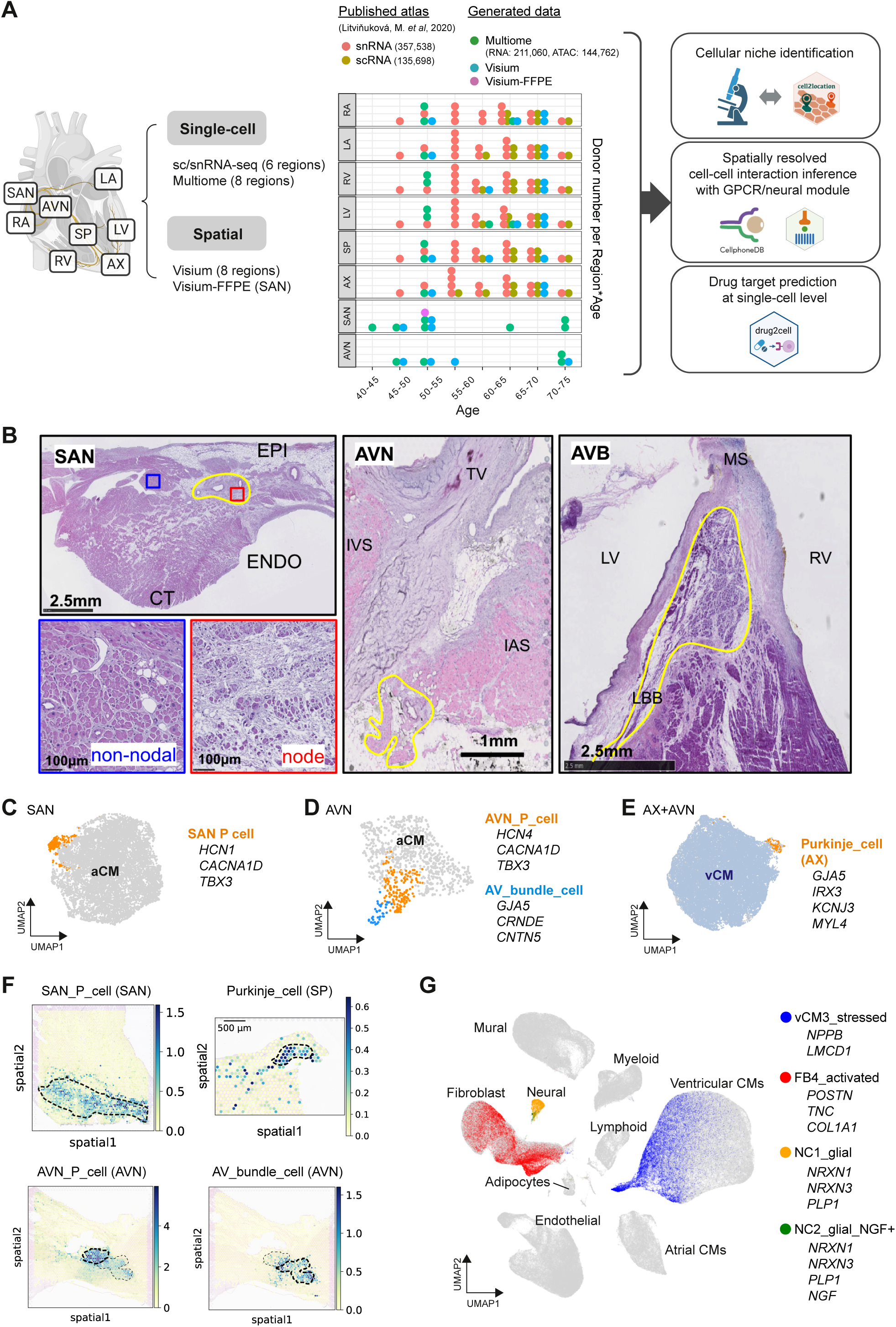
Multi-modal profiling of adult human cardiac tissues. A: Overview of study design and analysis pipelines. Multiome and Visium spatial transcriptomic data were generated from eight regions (RA, LA, RV, LV, SP, AX, SAN, AVN) of the adult human heart. sc/snRNA-seq data were integrated with a published human heart dataset. The dotplot shows the number of donors analysed per age group (x-axis) and region (y-axis). Dot colour represents data modality and the analysed cell or nuclei numbers of each modality are shown in brackets. The data were used for 1) cellular niche identification, 2) spatially-resolved cell-cell interaction analysis with CellPhone-DB with neural-GPCR module, and 3) drug target discovery analysis (drug2cell). B: H&E morphology of the CCS showing the SAN, AVN and AV bundle (AVB) (yellow bordered). Pacemaker cells (P cells) in the nodal tissue (red box) are smaller than neighbouring cardiomyocytes in non-nodal tissue (blue box) and embedded in dense ECM. The AVB is pictured with its transition to the left bundle branch. C,D,E: UMAP embedding of gene expression data of SAN aCMs (C), AVN aCMs (D), and AX and AVN CMs (E). Marker genes are displayed for CCS cells. F: Abundance of CCS cells obtained by mapping the indicated cell states to spatial transcriptomic data using cell2location. Spatial transcriptomics data were obtained from SAN, AVN and SP. Dashed lines highlight the histologically identified CCS regions including the nodes (SAN, AVN), the AV-bundle (AVN), and Purkinje cells (SP)(Supp. Fig. 3G). G: UMAP embedding of gene expression data depicts nine major cell types from eight cardiac anatomical regions. Four redefined non-CCS cell states are highlighted and marker genes indicated. **Abbreviations**: CCS (Cardiac conduction system); SAN (sinoatrial node); AVN (atrioventricular node); AVB (AV bundle); EPI (epicardium); ENDO (endocardium); CT (crista terminalis); IVS (interventricular septum); IAS (interatrial septum); TV (tricuspid valve); MS (membranous septum); LV (left ventricle); RV (right ventricle); LBB (left bundle branch); aCM (atrial cardiomyocyte); vCM (ventricular cardiomyocyte); AX (apex); SP (interventricular septum); LA (left atrium), RA (right atrium)

To capture cardiac conduction system (CCS) tissues from the SAN and AVN regions, we used a two-stage dissection protocol. In the SAN this involved an initial dissection of the posterolateral RA, which was then divided into 5mm-thick strips cut perpendicular to the *crista terminalis*, subsequently embedded and frozen in optimal cutting temperature (OCT) medium (**Supp. Fig. 1**). The presence of CCS structures was confirmed by hematoxylin and eosin (H&E) histology in sections from the same blocks (**Fig. 1B**). The SAN P cells were smaller and rounder than neighbouring cardiomyocytes, were surrounded by fibroblasts and dense extracellular matrix (ECM) and formed a discrete sub-epicardial structure at the junction of the *crista terminalis* and the RA posterior wall (**Fig. 1B**). For the AVN region a tissue sample including the Triangle of Koch (bordered by Todaro’s tendon, the coronary sinus ostium and tricuspid valve annulus) as well as the basal septum (spanning from interatrial to interventricular septum, and including the membranous septum) was dissected (**Fig.1B, Supp. Fig. 1**). As before, the sample was then divided into strips, cut perpendicular to the tricuspid annular plane. Each strip was then embedded in OCT medium and frozen, retaining information of its position in the septum-lateral wall axis. As confirmed by H&E, the lateral portions captured AV nodal tissue, whereas the septal portions included the AV bundle and its branches. After sectioning for histology, intact sections from these blocks were taken for spatial transcriptomic analysis and smFISH validation, with the remaining tissue being sectioned and homogenised to generate nuclei suspensions for multiome analysis (**Methods**).

After sample processing and quality control, 704,296 cells/nuclei (211,060 nuclei for newly generated Multiome data) were retained for gene expression and 144,762 nuclei for ATAC data downstream analyses (**Fig. 1A, Methods**). Integration of gene expression data was performed using scVI^11^ and scArches^12^ to account for batch variations such as donor, cell-or-nuclei, and the 10X Genomics chemistry version used (**Supp. Fig. 2A**) while retaining the regional differences of the atrial (RA, and LA), ventricular (RV, LV, SP, and AX), and conduction system (SAN, and AVN) samples (**Supp. Fig. 2B**)(Methods). Leiden clustering was performed on the neighbourhood graph created using batch-corrected scVI latent variables, and identified 12 coarse-grained cell types annotated using the expression patterns of curated lineage-specific gene markers (**Supp. Fig. 2C**).

To identify fine-grained cell states of the cardiac conduction system (CCS), SAN region atrial cardiomyocytes (CMs) were sub-clustered, revealing a clearly separated cluster of P cells (SAN_P_cell) which expressed canonical pacemaker channel genes (*HCN1*, *HCN4, CACNA1D*) and the transcription factor *TBX3*^7,13^ (**Fig. 1C, Supp. Fig. 3A**). Sub-clustering of AVN atrial and ventricular CMs (aCMs and vCMs) showed two CCS cell state clusters, P cells (AVN_P_cell) and AV bundle cells (AV_bundle_cell) (**Fig. 1D, Supp. Fig. 3B**). AV bundle cells formed a distinct cluster defined by enrichment of *GJA5* (encoding the gap junction protein Cx40), *CRNDE*, *CNTN5*, which were previously identified as markers of AV bundle cells in the mouse heart ^14^ (**Fig. 1D, Supp. Fig. 3C**). Purkinje cells (originally termed ‘fibers’^15^) are specialised CM-like cells which constitute the distal ramifications of the ventricular conduction system. They play an important role in propagating impulses from the AV bundle branches and their fascicles to the ventricular muscle, and are most abundant at the cardiac apex^16^. To identify this rare population at single-cell resolution, CM populations from AVN samples were integrated and clustered with those from the AX. This showed one cluster which contained, in addition to the CCS cells from the AVN, a population derived from the AX expressing Purkinje cell marker genes (*GJA5, IRX3, KCNJ3, MYL4*)(**Fig. 1E, Supp. Fig. 3D,E**)^17^. All the identified CCS cells were observed in multiple donors (**Supp. Fig. 3F**). By mapping identified CCS cell states to spatial transcriptomic data using cell2location^18^, we confirmed each cell state localised to the relevant histologically-identified CCS structures in SAN, AVN, the AV bundle in AVN, and the left bundle branch in intraventricular basal septum (**Fig. 1F, Supp. Fig. 3G**), supporting these annotations.

Other cell states were defined using label transfer^19^ with the published dataset^1^ as a reference. Minor revisions of annotation were included. One of these relates to the annotation of neural cells and is based on recent studies of the heart and the gut defining cells expressing *PLP1*, *NRXN1* and *NRXN3* as glial cells^20–23^. Here, cells expressing pan-glial gene markers and lacking core neuronal genes (**Fig. 1G, Supp. Fig. 3H**) are described as glia and are labelled below with the _glial suffix. ‘FB4’ was renamed as ‘FB4_activated’ based on the expression of FB activation signature genes (*POSTN*, *TNC*) which are responsive to TGFβ signalling and genes encoding ECM proteins, such as collagens (*COL1A1*, *COL1A2*, *COL3A1*) and fibronectin (*FN1*), at levels higher than other FB populations (**Fig. 1G, Supp. Fig. 3I**). ‘vCM3’ was renamed ‘vCM3_stressed’ since they express markers of cardiomyocyte stress (*NPPB*, *LMCD1*, *XIRP2*, *ACTA1*, *PFKP*) (**Fig. 1G, Supp. Fig. 3J**)^24–26^. ‘EC7_endocardial’ and lymphoid cell states were annotated based on a recently published study^27^. Myeloid cell state annotations were revised based on leiden clustering and marker expression (**Supp. Fig. 3K,L**). Thus, after extending the number of anatomical regions to eight by including the nodes, and refining past cell annotations, we defined a total of 75 cell states (**Supp. Fig. 2D**).

ATAC data was analysed by using ArchR^28^ to understand the chromatin landscape and the genomic regulation of cardiac cell identity. Peak calling was performed per fine-grained cell state annotated using the paired gene expression data. We performed dimensional reduction of the peak data using peakVI^29^ and computed a neighbourhood graph based on the peakVI latent space. The nuclei were embedded in a two-dimensional UMAP space based on the neighbourhood graph (**Supp. Fig. 4A**). Coarse grain cell types were clearly separated, indicating that each cardiac cell type has a distinct pattern of chromatin accessibility (**Supp. Fig. 4A,B**). In the vCM compartment, vCM3_stressed showed clear separation in the UMAP and the markers of cardiomyocyte stress were differentially accessible compared with other CMs (**Supp. Fig. 4A,C**). Differential chromatin accessibility analysis comparing CCS cells and other aCMs identified major markers for pacemaker cells such as ion channels (*CACNA1D* and *CACNA2D2*) and transcription factors (*ISL1*, *TBX3*, and *SHOX2*)(**Supp. Fig. 4D**), the latter representing a conserved genetic programme in the development of mouse and human SAN cells^30,31^.

### Unbiased discovery of cellular niches

To understand the cellular composition of microanatomical structures within each region, we mapped the fine-grained cell states defined by sc/snRNAseq analysis to spatial transcriptomic data using cell2location^18^ (**Fig. 2A**). In parallel, to examine the localization and enrichment of cell states within defined tissue structures, histological structural annotation based on H&E images was performed manually by an expert (YH). The mapping of expected cell types to histologically defined structures such as EC7_endocardial in the ‘endocardium’, glial cells (NC1_glial) in the ‘nerve’, arterial smooth muscle cells (SMC2_art) in the ‘vessel’, and fibroblasts and macrophages in ‘fibrosis’ regions (**Supp. Fig. 5A-D**), provides further validation of our approach. In the histologically annotated ‘node’ of SAN samples, we found that not only P cells but also other cell states, such as tissue-resident macrophages (MPs)(LYVE1+IGF1+MP)^32^ and fibroblasts (FBs) were enriched (**Fig. 2B,C** and **Supp. Fig. 6A,B**). Similarly, the annotated ‘node’ and ‘AV bundle’ regions of AVN samples also include FBs and MPs together with P cells or AV bundle cells, respectively (**Supp. Fig. 6A,C,D**). In contrast to results reported for mouse AVN, single-cell/nuclei data did not highlight connexin 43 (*GJA1*) and other connexin expressions in MPs and monocytes of SAN and AVN (**Supp. Fig. 6E**)^33^. In the epicardium of all four cardiac chambers, we detected enrichment not only of expected cell states (mesothelial cells, FBs, lymphatic endothelial cells (EC8_ln) and adipocytes), but also plasma B cells (B_plasma) and LYVE1+IGF1+MP (**Supp. Fig. 6F**).

**Fig. 2:**
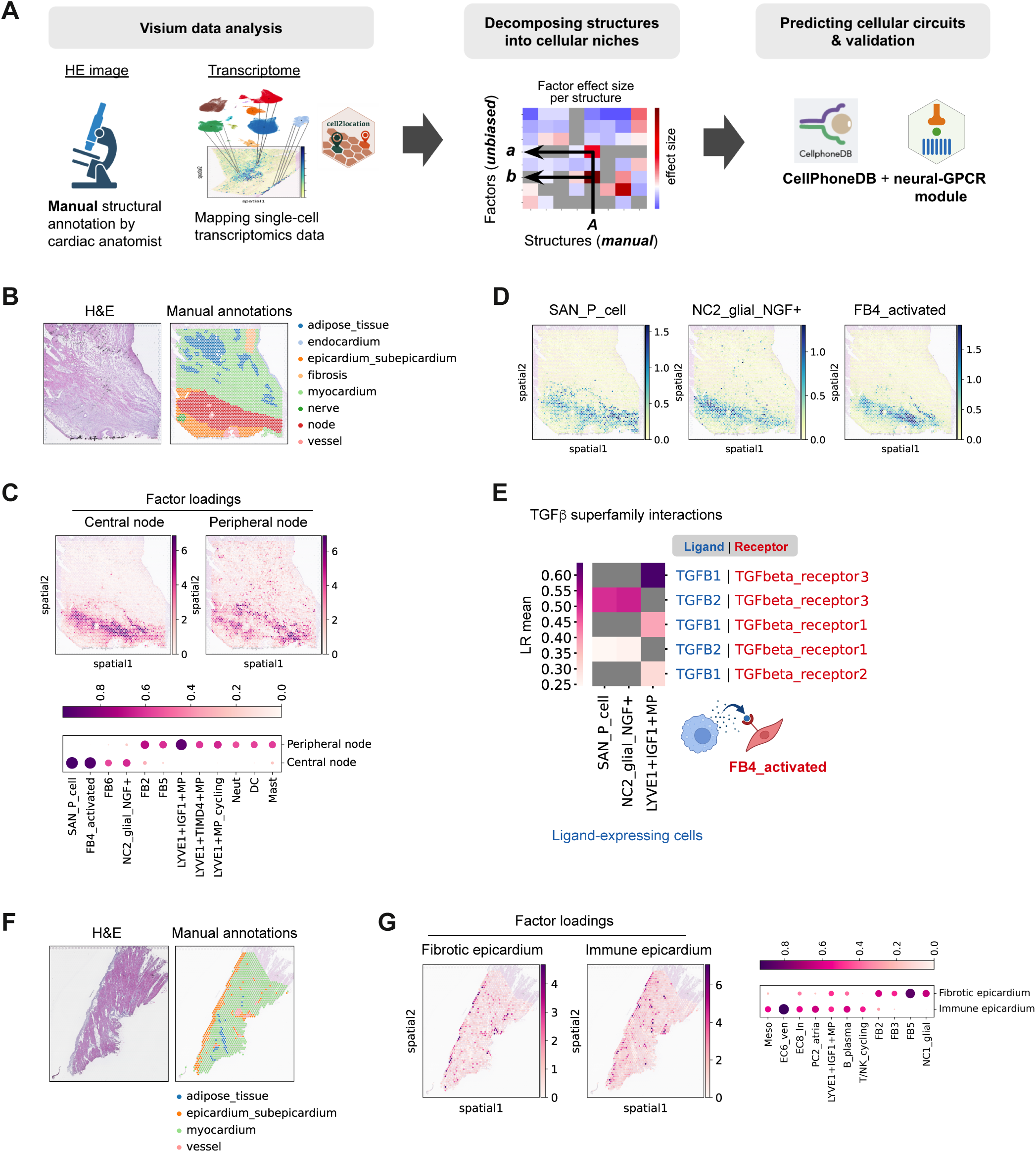
Identification of cellular niches in the adult human heart. A: Overview of the spatial data analysis workflow. Visium spots were annotated based on the accompanying H&E image. Cell states defined in sn/scRNAseq data were mapped to Visium spots using cell2location. Non-negative matrix factorisation (NMF) was used to decompose manually annotated regions into factors. Spatially-resolved analysis of cell-cell interactions was performed using NMF, histologically defined microenvironments, and a custom neural-GPCR CellPhoneDB extension module. B-G: Cellular microenvironment identification in the SAN (B-E) and the RV (F, G). Histological structures were manually annotated based on the H&E images (B, F). Factors from cell2location NMF analysis which showed high similarity (Cohen’s d) with individual histological structures were selected (as described in Methods). Factor loadings across locations (abundance of cell state group) are shown in spatial coordinates for selected factors (C, G). The accompanying dot plot illustrates factor loadings normalised per cell type (dot size and colour). Cell states with more than 0.4 in the selected factors are shown (C, G). Estimated abundance of representative cell states in the central node of SAN slide (D). Inferred spatial cell-cell interactions involving TGFβ superfamily member ligands (HGNC GID:1664) in the central nodal niche, signalling to cognate receptors in the FB4_activated cells (LR mean > 0.5) (E). **Abbreviations**: GPCR (G protein-coupled receptors); P_cell (pacemaker cell); H&E (hematoxylin and eosin); NMF (non-negative matrix factorization); LR mean (mean expressions of the interacting ligand-receptor partners); EC8_ln (lymphatic endothelial cell)

In order to identify cellular niches in an unbiased manner and decompose the H&E-discernible structures, we performed non-negative matrix factorisation (NMF) on the cell2location spot-by-cell output to identify ‘factors’ of co-occurring cell states^18^. We assessed the similarity between the NMF-identified factors and manually annotated structures by calculating an effect size of the spot factor loadings in a given structure compared to other areas (**Fig. 2A, Supp. Fig. 7, and Methods**). This revealed that NMF analysis can identify multiple distinct cellular niches within histologically defined structures. For example, the ‘node’ structure in SAN slides from two donors separated into central and peripheral regions (**Fig. 2C, Supp. Fig. 8A,B_(i)_,C_(i)_**). The central region contained P cells, FB4_activated, and NC2_glial cells expressing neuronal growth factor encoding gene *NGF* (NC2_glial_NGF+). The peripheral region was enriched for tissue-resident LYVE1+ MPs and other FBs including the basal FB2 (**Fig. 2C,D; Supp. Fig. 8B_(iii)(iv)_,C_(iii)(iv)_; Supp. Fig. 9A-D)**. These data indicate unambiguously for the first time that the human SAN is a compartmentalised structure, consisting of a central region with the functionally important pacemaker cells embedded amongst activated fibroblasts and glial cells, surrounded by a peripheral region of immune and other fibroblast populations. This two-layered structure of the SAN may contribute to insulating the P cells from the surrounding atrial tissue to optimise the source-sink relationship and maintain sinus node function^34,35^. In the ‘node’ and ‘bundle’ structures of the AVN region, we observed the enrichment of P cells and NC2_glial_NGF+ together with FBs and tissue-resident LYVE1+ MP, however, unlike the SAN, there was no concentric cell arrangement and FB4_activated cells were not abundant (**Supp Fig. 6A-D**).

Spatial transcriptomics data showed enrichment of genes encoding extracellular matrix (ECM) proteins and metalloproteinases in the node of SAN (**Supp. Fig. 9E**). Spatially resolved CellPhoneDB analysis^8,36^ of the node highlighted that *TGFB1* (LYVE1+IGF+MP) and *TGFB2* (SAN_P_cell), which are major mediators of FB activation^37^, interact with the receptors (*TGFBR1* and *TGFBR3*) expressed in FB4_activated (**Fig. 2E**). This suggests that cells from both central and peripheral nodes contribute to ECM formation.

In ventricular and atrial free walls, we assessed the similarities of NMF-identified factors compared with the manually annotated ‘epicardium-subepicardium’ structure (comprising the ‘true’ epicardium monolayer and subepicardial fat)(**Supp. Fig. 10A,B**). This integrated analysis permitted decomposition of the ‘epicardium-subepicardium’ structure into a distinct ‘immune epicardium’ niche (enriched for lymphatic endothelial and immune cells), and a ‘fibrotic epicardium’ niche (consisting of multiple fibroblast cell states)(**Fig. 2F,G; Supp. Fig. 10C,D**). Plasma B cells appeared in both epicardial niches, although at a higher proportion in the immune niche, in line with the enrichment analysis (**Fig. 2G; Supp. Fig. 10D**).

### Ion channel profile of CCS cells

Differential ion channel expression profiles (relative to non-CCS aCMs) highlighted the electrophysiological specialisation of the CCS cell states that we identify in our data: SAN P cells, AVN P cells, AV bundle cells, and Purkinje cells (**Fig 3A; Supp. Fig. 11B; Supp. Table 2**). In contrast to other cardiomyocytes, the pacemaker cell action potential upstroke is mediated by calcium rather than sodium influx^6^. In keeping with this, both the SAN and AVN P cells showed downregulation of the sodium channel gene *SCN5A* and upregulation of the L-type calcium channel gene *CACNA1D* (**Fig. 3A**). The two classical hyperpolarisation-activated cyclic nucleotide-gated (HCN) ‘pacemaker channel’ genes (*HCN1* and *HCN4*) were enriched in P cells from both nodes (**Fig. 3A**, phase 4). *CACNA1G*, encoding the T-type calcium channel (TTCC) ⍺ subunit (Cav3.1) known to be critical in pacemaking in rodents^6^, was only highly expressed in SAN P cells and the expression was less robust than *CACNA1D,* highlighting the importance of the latter in human pacemaking (**Fig. 3A**, phase 0). Several ion channels (*CACNA1D*, *HCN1*, *KCNJ3*, *TRPM3*, *CLIC5*) appear to have cross-species importance as they were also upregulated in mouse SAN P cells (relative to working atrial myocytes)^38,39^(**Supp. Fig. 11A**).

**Fig. 3:**
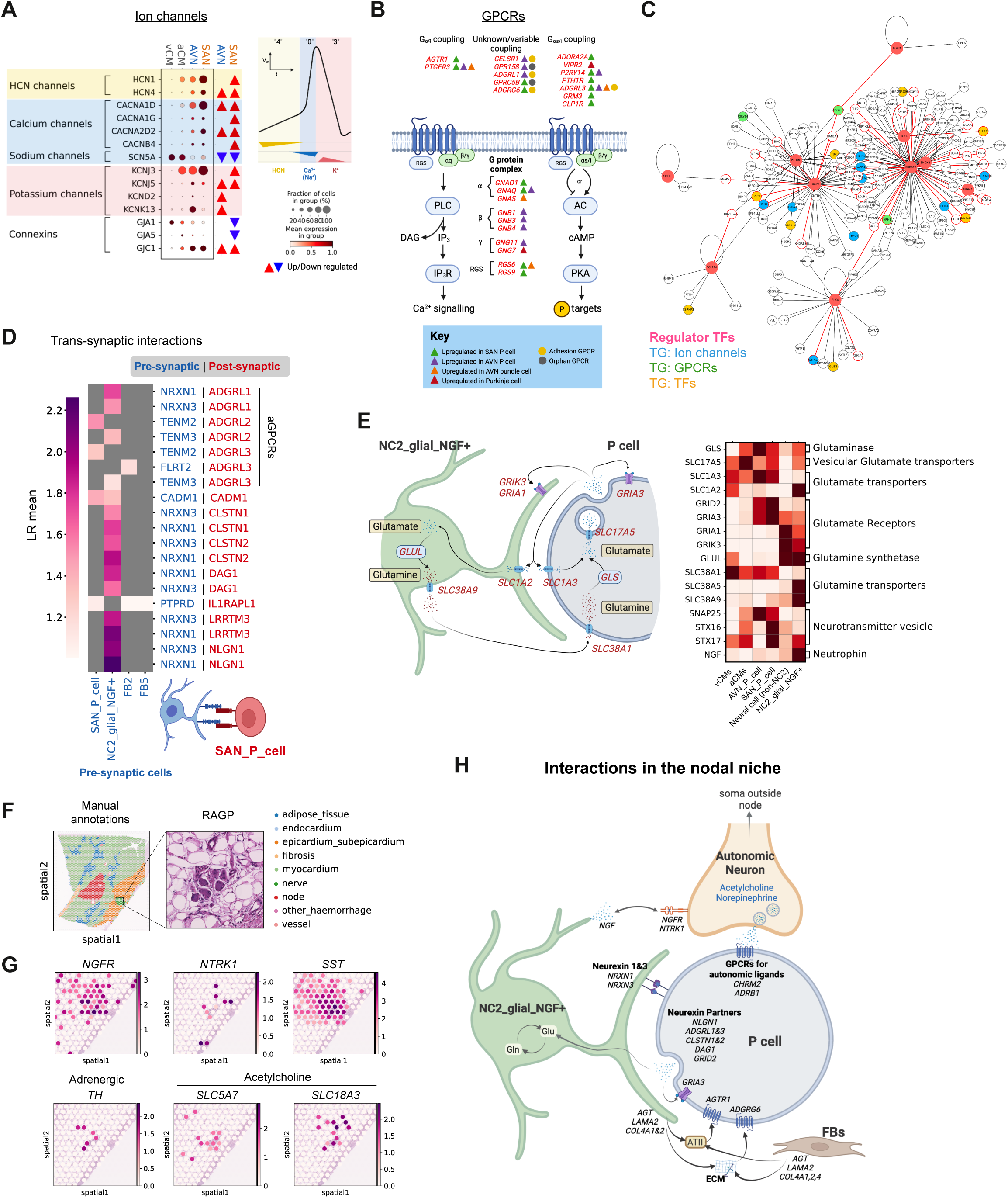
Characterisation of human CCS cells and the nodal niche. A: Dot plot shows the expression of DEGs encoding for ion channels (HGNC GID:177) in SAN and AVN P cells (relative to other aCMs), with reference to a typical pacemaker cell action potential. Classical ‘pacemaker channels’ (*HCN1*, *HCN4*) contribute to diastolic depolarisation (Phase 4, yellow). The upstroke (Phase 0, blue) is driven by calcium influx with Ca_V_1.3 (*CACNA1D*) as the dominant alpha subunit, with downregulation of the sodium channel Na_V_1.5 (*SCN5A*). Potassium channel opening mediates repolarisation (Phase 3, red). Colour-coded triangles represent the differentially expressing cell state as well as up- or down-regulation (up- or down-pointing). B: Schematic representation shows GPCR and G-protein signalling machinery genes that are upregulated (red text) in CCS cells compared to other aCMs. C: Analysis of transcription factor network governing P cells using pySCENIC. The network includes transcription factors (nodes coloured as salmon pink) and their predicted target genes differentially expressed in P cells. Transcription factor and target gene interactions inferred from the ATAC data analysis are highlighted in red (Methods). Target gene node colours are based on the class: GPCRs (green, HGNC GID:139), ion channels (blue, HGNC GID:177), or TFs (yellow). D: Inferred cell-cell trans-synaptic interactions (CellPhoneDB neural-GPCR module) between cell states of the central nodal niche (LR mean > 1). SAN_P_cells are the ‘receiver cells’ expressing genes encoding for receptors. E: Schematic of the paracrine glutamatergic signalling involving P cells and supported by NC2_glial_NGF+ (left). Matrixplot shows expression of glutamatergic signalling machinery genes in P cells and NC2_glial_NGF+ with aCM, vCM, and neural cells as cell types of reference (right). F: Manual structural annotations of a SAN visium-FFPE slide based on the associated H&E images. The RAGP ganglion is seen embedded in subepicardial adipose tissue, superficial to the SAN. G: Images show gene expressions in the RAGP (Visium, 10X Genomics). Genes encoding for NGF receptors (*NGFR*, *NTRK1*), Somatostatin (*SST*), the adrenergic marker tyrosine hydroxylase (*TH*), and the cholinergic markers including the high-affinity choline transporter (*SLC5A7*) and vesicular acetylcholine transporter (*SLC18A3*) are shown. H: Schematic of predicted cellular circuits in the nodal niches. Pacemaker cells are embedded in and interact with ECM proteins secreted by fibroblasts and NC2_glia_NGF+, forming synapse-like connections with the latter *via* neurexins which support paracrine glutamatergic signalling. NC2_glia_NGF+ releases Nerve Growth Factor which interacts with receptors on autonomic neurons, whose soma are present in the adjacent RAGP. P cells express receptors for autonomic neuron ligands (both parasympathetic and sympathetic represented by a single ‘autonomic neuron’), as well as Angiotensin II, which is produced locally in the node (NC2_glial_NGF+, fibroblasts). **Abbreviations**: DEG (differentially expressed gene); RAGP (right atrial ganglionated plexus); RGS (regulator of G protein signalling); LGIC (ligand-gated ion channel; ICNS (intracardiac nervous system); HGNC (HUGO Gene Nomenclature Committee); ATII (angiotensin II); ECM (extracellular matrix); TF (transcription factor); TG (target gene); LR mean (mean expressions of the interacting ligand-receptor partners).

Potassium channel currents contribute to the repolarisation of the membrane potential, thereby regulating P cell firing rate (**Fig. 3A**, phase 3). *KCNJ3* and *KCNJ5*, subunits of G protein-gated inwardly rectifying potassium (GIRK) channels were highly expressed in both P cell types (**Fig. 3A**). Amongst other potassium channels, *KCNN2* was highly expressed in both P cell types, whereas *KCNK13* was specific to AVN P cells (**Supp. Fig. 11B**). The former is activated by intracellular calcium whereas the latter is activated by a range of stimuli including arachidonic acid and purine receptor agonism^41^.

Various ion channel genes, not traditionally associated with cardiac pacemaking, were highly expressed in P cells. These included several members (*TRPC4*, *TRPM3*, *TRPM7*, *PKD2*, *MCOLN2*, *MCOLN3*) of the transient receptor potential (TRP) family of non-selective cation channels (**Supp. Fig. 11B**), with *TRPM3* being robustly expressed in SAN P cells. *ANO6,* encoding a calcium-activated chloride channel, was upregulated in SAN and AVN P cells as well as AV bundle cells (**Supp. Fig. 11B**). The roles of these unexpected ion channels in CCS cell function require further investigation.

Altogether these data provide a highly specific map of genes and cells of the conduction system essential for designing novel refined functional studies and defining cell-specific druggable targets.

### GPCR profile of CCS cells

Heart rate is tightly regulated by a variety of neurohumoral factors, primarily via GPCR signalling^44^. We used the HUGO Gene Nomenclature Committee (HGNC) gene group of GPCR genes (group ID: 139) to comprehensively assess their expression in P cells using an unbiased data-driven approach to discovery. We found that CCS cells exhibit a wide variety of GPCRs, G protein subunits, and second-messenger machinery compared to other atrial cardiomyocytes (**Fig. 3B; Supp. Fig. 11C,D**). Within the receptor genes were the classical acetylcholine muscarinic (M_2_) receptor (*CHRM2*) and catecholamine β receptors (*ADRB1*), responsible for mediating the major heart rate-modulating effects of the autonomic nervous system (ANS)^45^. In addition, for the first time in humans, we found receptors for several neurohumoral ligands in P cells, including Angiotensin II (*AGTR1*), Calcitonin gene-related peptide (*CALCRL*), glucagon-like peptide 1 (*GLP1R*), parathyroid hormone (*PTH1R*), and vasoactive intestinal peptide (*VIPR2*). The adhesion GPCR (aGPCRs) family, which are activated by the binding of their extracellular adhesion molecule domain to ligands on neighbouring cells or within the ECM, and are involved in neuronal cell migration and synaptogenesis^46^, were highly represented within CCS cells (*ADGRL3*, *ADGRG6*, *CELSR1*)(**Fig. 3B and Supp. Fig. 11C**).

To enable comprehensive analysis of GPCR interactions in the node, we supplemented the CellPhoneDB database^8^ with a custom module of GPCR and trans-synaptic adhesion molecule interactions, adding more than 1000 new ligand-receptor pairs (**Methods; Supp. Table 3,4**). Spatially resolved CellPhoneDB analysis within the neural-GPCR module showed multiple interactions through aGPCRs expressed in SAN P cells. Extracellular matrix proteins including laminin (*LAMA2*) expressed in NC2_glial_NGF+ and FBs interact with the receptor encoded by *ADGRG6* in SAN P cells (**Supp. Fig. 11E**). Interaction of angiotensinogen (*AGT*), the precursor peptide for the ligand angiotensin II and its receptor encoded by *AGTR1* was predicted in the nodes of both SAN and AVN (**Supp. Fig. 11E,F**). Spatial transcriptomics data confirmed *AGT* expression in the manually-annotated ‘node’ region (**Supp. Fig. 11G**). This is consistent with the identification of local renin-angiotensin circuits^47^ and a previous study showing that the angiotensinogen protein is expressed in the human heart and enriched in the CCS^48^. Taken together, these GPCR repertoires support CCS cell responsiveness to an array of neurohumoral factors, as well as interactivity with ECM components and cells in their niche.

### Gene regulatory networks in P cells

To reveal gene regulatory networks that influence P cell gene expression profiles, we combined pySCENIC^49^ analysis with gene accessibility information derived from the snATACseq component of our multiome snRNA+ATACseq data. We constructed a network of P cells based on transcription factors (TFs) and their predicted target genes differentially expressed in P cells (**Supp. Fig. 12A**). TF and target gene interactions inferred from the ATAC data analysis were largely overlapping with the network predicted by gene expression (**Fig. 3C; Supp. Fig. 12B**).

Interestingly, our results indicate that *FOXP2*, a TF associated with language centre development in the brain, is a key TF targeting major P cell ion channel genes, such as *HCN1* and *CACNA1D,* as well as *TBX3* (a repressor of working myocyte gene programmes^30,31,50^) (**Fig. 3C**)*. FOXP2* and *HCN1* interaction was also observed in the ATAC analysis (**Supp. Fig. 12C**). This may explain why mice carrying one non-functional *FOXP2* allele have lower pulse rates than wild type littermates^51^. SREBF2, a TF binding the sterol regulatory element 1 motif^52^, is predicted to target multiple genes encoding ion channels, GPCR, and TFs. *PRDM6*, associated with heart rate recovery during exercise^53^ regulates multiple ion channels and GPCR genes. *CREB5*, a cAMP response element-binding protein (downstream of G_⍺s_ signalling) associated with atrial fibrillation^54,55^, was also one of the TFs differentially expressed in P cells.

Overall, this analysis points towards new TFs governing core P cell gene programmes, contributing mechanistic insights into the regulators of ion channels and GPCRs critical to P cell function. These predicted control mechanisms may be useful in engineering better models of human P cells *in vitro*.

### NC2_glial_NGF+ support CCS cell glutamatergic signalling

In the manually annotated ‘node’ region of both SAN and AVN regions, we observed enrichment and colocation of P cells with NC2_glial_NGF+ cells (**Fig. 2C**, **Supp. Fig. 6A**, and **Supp. Fig. 8**). This population expresses neurexins (*NRXN1* and *NRXN3*) which form trans-synaptic complexes with cell adhesion molecules. Our neural-GPCR CellphoneDB module highlighted several such interactions with P cells (*NLGN1*, *ADGRL1, ADGRL3*, *DAG1, CLSTN1*)^56,57^ (**Fig. 3D; Supp. Fig. 13A**).

We also discovered that P cells and NC2_glial_NGF+ cells express genes involved in glutamatergic signalling. CCS cells differentially express the glutamate receptor *GRIA3* (an AMPA-type receptor) and *GRID2* (**Fig. 3E**), known as a postsynaptic auxiliary subunit which forms a ‘molecular bridge’ to the presynaptic membrane^58^. The glutamate transporter *SLC1A3* and synthetic enzyme *GLS* as well as synaptic vesicle genes such as *SNAP25* and *STX7* are expressed in both SAN and AVN P cells (**Fig. 3E**). Spatially resolved CellPhoneDB analysis of the central nodal niche in the SAN or the ‘node’ structure of AVN highlighted P cell-to-P cell glutamatergic signalling *via* three ionotropic glutamate receptors (*GRIA3*, *GRIK1* and *GRIK2*) in both SAN and AVN (**Fig. 3E; Supp. Fig. 13B,C**). NC2_glial_NGF+ cells express genes involved in the glutamate-glutamine cycle, the means by which astroglia replenish glutamine pools for neighbouring glutamatergic neurons^59^: the glutamate reuptake transporter EAAT2 (*SLC1A2*), glutamine synthetase (*GLUL*) which remakes glutamine from glutamate, and the glutamine transporter (*SLC38A9*) releasing extracellular glutamine (**Fig. 3E**). Conversely, P cells express glutamine transporter (*SLC38A1*) as well as glutaminase (*GLS*) which converts glutamine to the active neurotransmitter glutamate (**Fig. 3E**).

Glutamatergic signalling has recently been reported in rodent cardiomyocytes, with effects on cell excitability^60^. Our findings suggest that human CCS cells express the requisite machinery for such signalling, and furthermore that NC2_glial_NGF+ may form synapses with CCS cells, playing an astrocyte-like support role.

### Innervation of the nodes

While the majority of cardiac autonomic neurons reside in extracardiac structures (the paravertebral ganglia and brainstem), a number of neurons are native to the heart, forming the Intrinsic Cardiac Nervous System (ICNS)^61^. The ICNS neuronal bodies are concentrated within the subepicardial fat in structures known as ganglionated plexi (GP)^62–64^. Our snRNAseq data do not contain cells matching the profiles of autonomic neurons as expected based on single-cell profiling of extracardiac ganglia^65^, likely due to the rarity of ICNS neurons. However, in the H&E images accompanying spatial transcriptomics slides (in six sections across three donors), we identified the right atrial ganglionated plexus (RAGP)(**Fig. 3F**), which houses the somata of neurons innervating the SAN^64^.

A score calculated for expression of pan-neuronal cytoskeletal markers (*PRPH*, *NEFL*, *NEFM*, *NEFH*) mapped to the Visium voxels that sit in the RAGP region of the corresponding H&E images (**Supp. Fig. 14A**). Correlating this score with gene expression for the corresponding Visium voxels allows us to reveal the first transcriptome-wide profile of a human GP (**Supp. Fig. 14B; Supp. Table 5**). This allows us to delve into multiple novel insights regarding human intra-cardiac neuronal characteristics.

For instance, the neuropeptide somatostatin (*SST*), previously identified as a marker of RAGP neuronal populations in pig^66^ was highly correlated with the pan-neuronal cytoskeletal score (**Supp. Fig. 14B**). Amongst the significantly correlated genes were the cholinergic markers (*SLC18A3*, *SLC5A7*, *CHAT, ACHE*) as well as the catecholamine synthetic enzymes (*TH*, *DDC*) and the sympathetic co-transmitter Neuropeptide Y (*NPY*)(**Fig. 3G; Supp. Fig. 14B,C**). The relevant corresponding receptors (adrenergic and cholinergic) are expressed in P cells (**Supp. Fig. 14C**). Several markers typical of neuroendocrine cells such as Synaptophysin (*SYP*), Secretogranin II (*SCG2*), Chromogranin A (*CHGA*), and chromaffin granule amine transporter (*SLC18A1*) were also correlated with the pan-neuronal cytoskeletal score **(Supp. Table 5)**.

Cardiac sympathetic neuron viability is dependent on a continuous source of the neurotrophin Nerve Growth Factor (*NGF*)^62,67^, with recent work in rats suggesting cardiomyocytes as a source^68,69^. However, amongst the neural cells from our human dataset, NC2_glial_NGF+ specifically expresses this factor (**Supp. Fig. 3G**). Furthermore, expression of NGF receptors p75NTR (*NGFR*) and TrkA (*NTRK1*) were correlated with the pan-neuronal cytoskeletal markers (**Fig. 3G; Supp. Fig. 14B**). In porcine RAGP, *NTRK1* expression was found specifically in neurons with axonal projections to the SAN (and not in other neurons)^66^. Thus, nodal NGF signalling (from NC2_glial_NGF+) may promote and maintain innervation from the nearby RAGP.

Overall, we propose a nodal cellular circuit wherein NC2_glial_NGF+ play a pivotal role: synaptically interacting with P cells, facilitating glutamatergic signalling (which potentially modulates firing rate), and promoting autonomic innervation *via* NGF signalling (**Fig. 3H**).

### Drug2cell: drug target discovery at single-cell level

The cells of the CCS and auxiliary structures such as the ICNS are important targets for chronotropic drugs. Therefore, we sought to leverage our human heart data to map drugs to target-expressing cells. Several single-cell studies and methods have been published which use drug-response transcriptional signatures, obtained from cell line experiments, and data-mining^70,71^ to predict drug effects. However, these require the development of cell type-specific *in vitro* models, which (despite their increasing complexity^72^) can fail to fully recapitulate the profiles of their *in vivo* counterparts. Using our sc/snRNAseq data as a reference, we took pairs of drug and target molecules from the ChEMBL database (https://www.ebi.ac.uk/chembl/) to comprehensively evaluate drug target expression in user-provided single-cell data and annotated cell populations (**Fig. 4A, Methods**). We developed a Python package, drug2cell, to streamline the workflow and apply selected drug-target pairs to single-cell datasets (**Fig. 4A**). Drugs and target molecules can be filtered based on quantitative bioactivity metrics, drug categories and clinical trial phases, and target molecule classes. Once drug scores are calculated, it is possible to 1) find cells which are targeted by drugs of interest, 2) find drugs which target cells of interest, and 3) find target molecules expressed in the target cells and potentially mediate a drug’s effect. (**Fig. 4A**).

**Fig. 4:**
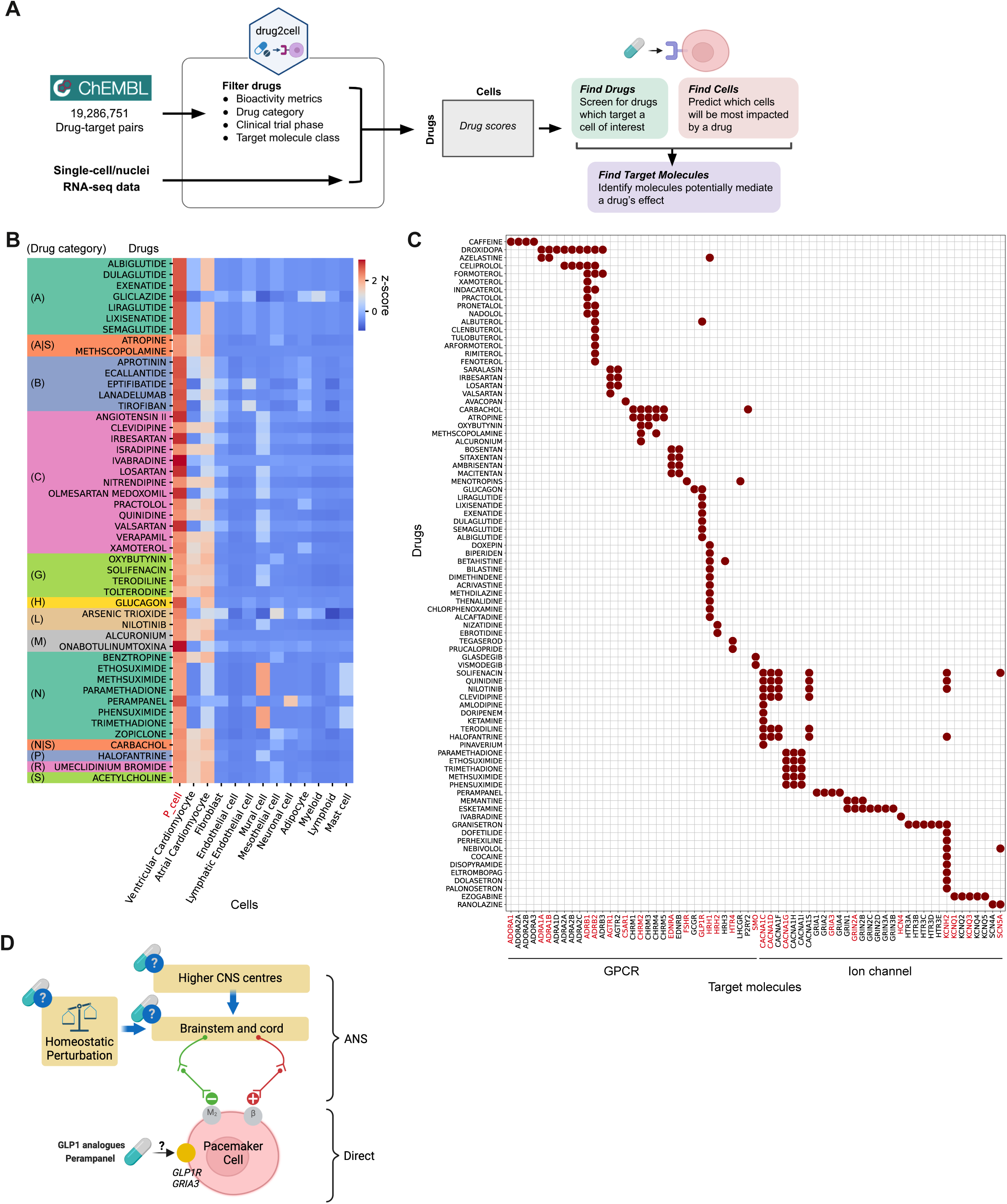
Drug target exploration at the single-cell level. A: Schematic of drug2cell analysis workflow. Drug2cell takes pairs of drugs and their target molecules from the ChEMBL database (https://www.ebi.ac.uk/chembl/). Drugs and target molecules can be queried and filtered based on quantitative bioactivity metrics, drug categories (ATC classification), clinical trial phase, and classes of molecular targets. The scores of the selected drugs in single cells or cell populations is calculated based on the target gene expressions in sn/scRNAseq data (as described in Methods). Once drug scores are calculated, one can 1) find cells which are targeted by drugs of interest, 2) find drugs which target cells of interest, and 3) find target molecules expressed in the target cells to infer a drug’s effect. B: Hetmap shows scores of drugs that are significantly higher in P cells compared with other cell states in the heart (t-test, adjusted P-value < 0.05, log fold change > 2). Heatmap colours were scaled by the z-scores of the drug scores on each cell state. The column reporting drug names is coloured by drug category (ATC classification): A (alimentary tract and metabolism), S (sensory organs), B (blood and blood forming organs), C (cardiovascular system), G (genito-urinary system and sex hormones), H (systemic hormonal preparations, excl. sex hormones and insulins), J (antiinfectives for systemic use), L (antineoplastic and immunomodulating agents), M (musculo-skeletal system), N (nervous system), P (antiparasitic products, insecticides and repellents), R (respiratory system). C: Dotplot shows clinically approved drugs which target GPCRs or Ion channels and genes encoding for their molecular targets. Maroon dots pinpoint the drug-target pairs predicted in the P cells. Target genes which are expressed in at least 10% of P cells are highlighted in red. D: Single cell profiling provides novel insights into chronotropic drug sites of action. Schematic shows ANS input to SAN P cells and potential druggable sites affecting P cells firing. Parasympathetic (green) and sympathetic (red) branches of the ANS are activated by homeostatic perturbations as well as CNS input, acting on M_2_ and β receptors to slow or accelerate firing, respectively. Drugs may exert chronotropic effects via the ANS (blue circles) or directly on P cells. Single-cell profiling reveals that P cells express an array of genes encoding targets of chronotropic drugs (yellow circle), potentially explaining direct (ANS-independent) action of the drugs on P cells, such as in the case of the GLP1 analogues and Perampanel. **Abbreviations**: CNS (central nervous system); ANS (autonomic nervous system); ACh (acetylcholine), NE (norepinephrine); ATC (Anatomical Therapeutic Chemical)

As an example, we explored clinically approved drugs of all categories (Anatomical Therapeutic Chemical classification) registered in ChEMBL to identify the drugs with the strongest predicted effect on P cells compared with other cardiac cell states (**Fig. 4B**). This highlighted the group of drugs belonging to the ‘Cardiovascular system’ category, including Ivabradine (HCN4 inhibitor), Quinidine (class I antiarrhythmic agent) and Atropine (anticholinergic agent) which are clinically used drugs with chronotropic effects and are known to act on P-cells^73,74^. To further explore other groups of drugs that potentially affect P cell function, we searched broadly for clinically approved and preclinical bioactive molecules with drug-like properties which target GPCRs or ion channels. As a result, we found drugs, such as beta-adrenoceptor blockers, which target GPCRs (*ADRB1*,*2*, *CHRM2*, and *GLP1R)* and ion channels (*CACNA1C,D,G*, *HCN4*, and *TRPV1*) expressed in P cells (**Fig. 4C; Supp. Fig. 15**). In addition, we found several drugs which block angiotensin II receptors (*AGTR1*), suggesting those drugs potentially have a direct effect on P cells.

Importantly, our analysis identified P cell-expressed targets for non-cardiac medications with documented chronotropic effects (**Fig. 4B,C**). These included the anti-diabetic medication Liraglutide (GLP1 analogue), belonging to ‘Alimentary tract and metabolism’, and the anti-epileptic medication Perampanel (glutamate AMPA receptor inhibitor), belonging to ‘Nervous system’ (**Fig. 4B,C**). Both drugs are known to alter heart rate, but since their targets were not known to be present in P cells prior to this study, alternative sites of action (the ANS and CNS, respectively) had been previously proposed as mediators of the side effects^75,76^ (**Fig. 4D**).

In summary, drug2cell can identify drug-impacted cells in user-provided single-cell datasets, potentially revealing hidden mechanisms of action and predicting drug impact on cell types of interest.

### Immune niche in the epicardium

To further understand the immune niche incorporating epicardial plasma B cells (**Fig. 2G; Supp. Fig. 16A**), we first localised plasma B cells and the associated immunoglobulin heavy chain encoding genes in the spatial transcriptomics data. This showed distinct localisation of the cluster of spots expressing *IGHG1* and *IGHA1* in the ‘epicardium-subepicardium’ histological structure (**Fig. 5A**). The immunoglobulin heavy chain genes (*IGHG1*, *IGHG3*, *IGHG4*, and *IGHA1*) were significantly enriched in this region (**Supp. Fig. 16B**). Interestingly, the expressions of IgG genes were anti-correlated with *IGHA1* (**Fig. 5B**), suggesting different mechanisms regulate the fate of IgG+ and IgA+ plasma B cells within the ‘epicardial-subepicardial’ niche.

**Fig. 5:**
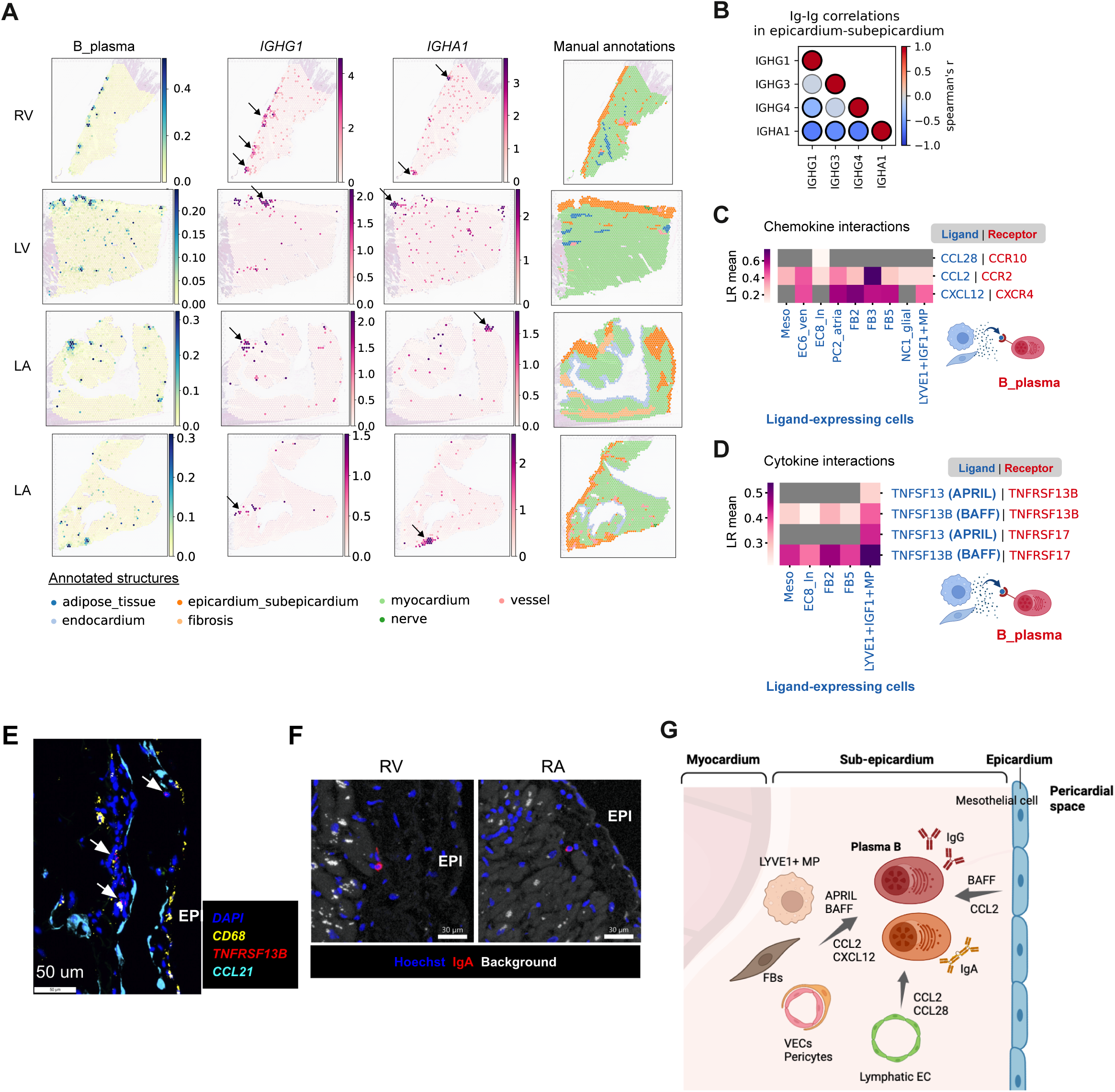
Immune niche in the epicardium. A: Images show abundance of plasma B cells (mapped by using cell2location), mapping of *IGHG1* and *IGHA1* expressions, and manual annotation of histological structures in spatial transcriptomic sections from the indicated anatomical regions. B: Spearman correlation between paired IGH gene expressions in the ‘epicardium-subepicardium’. Colour scale indicates coefficient (Spearman’s r) and significant correlations are indicated by thick edges (adjusted p-value < 0.05). C, D: Heatmaps summarise the inferred cell-cell interactions mediated by chemokines (HGNC, GID:189) (C) and cytokines (HGNC GID:599,602,1932,781,1264) (D) with their cognate receptors expressed on plasma B cells. The cells sending signals (x-axis) are those identified in the spatially refined niche. E: Multiplex single-molecule fluorescence *in situ* hybridization of the LV epicardial region stained with RNAscope probes against *TNFRSF13B* (red), *CCL21* (turquoise), and *CD68* (yellow) and DAPI (blue) for nuclei. F: Immunofluorescence of the epicardial regions (RV and RA) stained with anti-IgA antibody (red) and Hoechst (blue). In white non-specific background signal. G: Schematic of the immune niche: the epicardial enrichment of plasma B cells and neighbouring macrophages, fibroblasts, and endothelial cells suggests that plasma B cells may secrete immunoglobulins into the pericardial space, potentially providing an immune barrier against invading pathogens. **Abbreviations**: IGH (immunoglobulin heavy chain); MP (macrophage); FB (fibroblast); EC (endothelial cell); VEC (vascular endothelial cell); LR mean (mean expressions of the interacting ligand-receptor partners).

To understand the mechanisms regulating the dynamics of this cellular niche, we performed spatially resolved CellPhoneDB analysis (**Supp. Fig. 16C**). Given that endothelial cells, FBs, mesothelial cells and MPs expressed genes encoding for *CCL2* and *CXCL12,* and plasma B cells express genes encoding for their receptors, *CCR2* and *CXCR4* (**Fig. 5C**), we inferred that plasma B cells may be recruited via a chemokine-dependent mechanism regulated by cells present in the epicardial/subepicardial niche^77–79^. Our data point towards a CCL28-CCR10 interaction from lymphatic endothelial cells (EC8_ln) to IgA+ plasma B cells, suggesting a specific mechanism of recruitment^80^ (**Fig. 5C**). Our results also suggest that Tumor necrosis factor (TNF)-dependent signalling is enriched in the epicardial immune niche. Ligand-encoding genes TNFSF13B (*BAFF*) and TNFSF13 (*APRIL*) were expressed in MPs, monocytes and FBs, while their receptor counterparts TNFRSF13B (*TACI*) and TNFRSF17 (*BCMA*) were expressed in plasma B cells (**Fig. 5D**). This suggests a fine regulation is in place to ensure the homeostasis of the niche and potentially prevent autoimmune mechanisms^81^. Consistent with our CellPhoneDB predictions, RNAscope multiplex smFISH analysis confirmed the co-localisation of plasma B cells expressing *TNFRSF13B* (BAFF receptor) alongside MPs in the epicardium (**Fig. 5E**). Immunofluorescence staining also confirmed the presence of IgA in the sub-epicardial region (**Fig. 5F**). Furthermore, CellPhoneDB predicts interaction between plasma B cells and FBs (FB2, FB3, and FB5), LYVE1+IGF1+MP, and endothelial cells (EC6_vec and EC8_ln), via TGFβ1 and TGFβ receptors (**Supp. Fig. 16D**). Plasma B cells, LYVE1+IGF1+MP and endothelial cells were the major sources of *TGFB1* (**Supp. Fig. 16E**). This suggests that plasma B cells in the epicardium may contribute to ECM formation, consistent with a previous study showing the role of plasma B cells in fibrosis^82^.

The manually-annotated ‘epicardial-subepicardial’ structure highly expressed genes for secreted antimicrobial defence molecules, including *SLPI*^83^ and *RARRES2* (encoding Chemerin)^84^, as well as monocyte/neutrophil-attracting chemokines (*CXCL1*, *CXCL6*, and *CXCL8*). Upon inspection of the matched snRNAseq data, these genes were expressed by Mesothelial cells (**Supp. Fig. 16F,G**). These epicardial cells therefore contribute to the immune defensive function of this outer cellular zone of the heart.

In summary, we defined a novel epicardial niche where MP- and stromal cell-derived signals mediate the recruitment and homeostasis of plasma B cells that secrete immunoglobulins. These antibodies are likely secreted in the epicardium layer, and together with other antimicrobial molecules, provide an immune barrier protecting the heart against invading pathogens (**Fig. 5G**).

### Ventricular myocardial-stress niche

Above, we noted activated fibroblast cell states with high collagen expression profiles and stressed ventricular CMs expressing cardiomyocyte stress signatures (**Fig 1G**). To further probe the role of these cells, we investigated their abundance and localisation in publicly available datasets of dilated cardiomyopathy (DCM) and hypertrophic cardiomyopathy (HCM)^27,85^. By mapping identified cell states to the disease dataset and analysing changes in cell state proportions, we detected expansions of FB4_activated and vCM3_stressed populations in both DCM and HCM samples compared with healthy control samples (**Supp. Fig. 17A,B**). We examined ventricular myocardium samples from dilated cardiomyopathy (DCM) patients using multiplex single-molecule RNA fluorescence *in-situ* hybridization (smFISH). This revealed the co-location of FB4_activated and vCM3_stressed marker genes (*COL1A1* and *NPPB*), which was increased compared to healthy controls (**Supp. Fig. 17C**). Within our spatial transcriptomic data of the septum region of the healthy heart, we found these cell states focally co-located in a region with high expression of their marker genes (*COL1A1* and *NPPB)* (**Supp. Fig. 17D,E**, **Methods**). Other cell states, such as vascular cells and immune cells, were also enriched in this myocardial-stress niche (**Supp. Fig. 17F**). Spatially-refined CellPhoneDB analysis suggested that the TGFβ superfamily contributes to intercellular communications within this niche: endothelial cells, smooth muscle cell (SMC2_art), FBs (FB4_activated and FB5), and immune cells (MPs and NK cells) expressed genes encoding for TGFβ ligands, with TGFβ receptors being expressed in both FB4_activated and vCM3_stressed populations (**Supp. Fig. 17G**). In addition, several endothelial cell populations and SMC2_art expressed inflammatory cytokines (*IL6*, *TNFSF10*, and *TNFSF12*) which interact with their cognate receptors expressed in vCM3_stressed (**Supp. Fig. 17H**), suggesting a direct proinflammatory effect towards the cardiomyocytes. In fact, compared with other vCMs, vCM3_stressed expressed higher levels of IL-6 and TNFSF12 (TWEAK) cytokine receptors (*IL6*, *TNFRSF12A*) as well as interleukin-1 receptors (*IL1R1* and *IL1RAP*) (**Supp. Fig. 17I**) which are associated with the pathogenesis of heart failure^86–88^.

We speculate that this myocardial-stress niche may be a precursor to adverse cardiac remodelling and pathogenic fibrosis typical of cardiomyopathy (**Supp. Fig. 17J**). We hypothesise that even in the absence of clinical heart disease there may be a spectrum of tissue health, with areas of incipient micropathology, as found in our samples from deceased transplant donor tissue.

## Discussion

Here, we develop computational approaches to combine and interpret spatial and single-cell multi-omic data from donor adult human hearts. Our work leads to the assembly of a refined heart cell and gene atlas encompassing 8 anatomical regions that can be used as a reference for disease studies including those affecting the cardiac conduction system.

The analysis of P cells confirms the importance of the slowly inactivating L-type calcium channel Ca_V_1.3 (*CACNA1D*) in cardiac pacemaking: being the most robustly expressed ion channel gene in both human pacemaker cell types, as well as in mice. This is consistent with the bradycardia observed in patients affected by the sinoatrial node dysfunction and deafness syndrome caused by deleterious *CACNA1D* mutations ^89,90^ and with the recurring implication of this gene in heart rate-related GWAS studies^91–93^. Based on co-expression and open chromatin patterns, we predict *FOXP2* as a key TF regulating *CACNA1D* expression, and also that of *HCN1*, another critical pacemaker channel gene. This TF has not been previously associated with the regulation of cardiac channel gene expression, and its role requires further investigation and validation.

Unexpectedly, P cell show enrichment of ion channels not traditionally associated with cardiac pacemaking, including several members of the transient receptor potential (TRP) family of non-selective cation channels (*TRPC4*, *TRPM3*, *TRPM7*, *PKD2*, *MCOLN2*, *MCOLN3*). These are regulated by a variety of physicochemical stimuli and their opening has been shown to modulate cardiomyocyte excitability^94^, thus their depolarising current could directly alter P cell firing rate. *TRPM3*, involved in noxious heat sensation by sensory neurons^95^, is inhibited by the βγ subunits of G_i/o_-coupled GPCRs^96^. It has been implicated in SAN- and AVN-associated GWAS traits^91,97^ and could potentially modulate P cell excitability in a fashion similar to GIRK channels ^98^. We also report *ANO6* expression in the human conduction system for the first time. This encodes a calcium-activated chloride channel, which is notable since another member of the same anoctamin family (*ANO1*) is the primary pacemaker channel responsible for peristaltic ‘slow-wave’ generation in the gut’s Cajal cells^21,99,100^, and was detected in mouse SAN^39^.

Furthermore, we discovered that human CCS cells express machinery enabling paracrine glutamatergic signalling, via a glutamate-gated ion channel (*GRIA3*), similar to cardiomyocyte-cardiomyocyte glutamate signalling recently observed in rodents (also via *Gria3*)^60,101^. Opening of these ionotropic channels increases cardiomyocyte excitability and may directly accelerate P cell firing.

We also show that the neural population NC2_glial_*NGF*+ is a niche-partner of CCS cells, being found in both nodes and the AV bundle. These cells express key elements required to maintain the glutamine pool and may therefore facilitate cardiac glutamatergic signalling in an astrocyte-like role. Similarly, our analysis highlights multiple trans-synaptic adhesion interactions suggesting a synapse-like interconnection. Finally, in a fashion similar to that proposed for cardiomyocyte-neuron signalling^68,69^, NC2_glial_*NGF*+ may promote CCS innervation by expressing the neurotrophic factor *NGF*, whose receptors (*NGFR*, *NTRK1*) were expressed in the nearby RAGP. Taken together these findings suggest NC2_glial_*NGF*+ play an important multifaceted role supporting CCS cells.

Besides niches related to cardiac electroanatomy, we shed light on cardiac immune microenvironments enriching for immune cells. Here, for the first time, we detect IgA+ plasma B cells in the human heart, and demonstrate their localisation in epicardial foci, which are distinct from foci of IgG+ plasma B cells. The niche contains plasma B cells but not undifferentiated B cells or T follicular helper cells which support B cell differentiation^102^. This suggests plasma B cells are either recruited to the niche after differentiation in a draining lymph node, or develop locally in a T-cell independent manner. This is consistent with the epicardium being the innermost layer of the pericardium, a multilayered structure which encloses the heart and provides a barrier to direct invasion of pathogens infecting the neighbouring lungs^103^. Mesothelial cells may have a key role in the formation of the epicardial immune niche, mediated by the expression of chemokine (*CCL2*) and pro-survival factors for B cells (*BAFF*) similarly to the immunoregulatory function in the peritoneal cavity ^104^. Indeed, we report the expression of antimicrobial molecules in mesothelial cells, which also supports a role of the epicardium as a barrier against invading pathogens. Therefore, we propose that the epicardial immune niche represents a defence structure, poised to respond to future infections.

Besides immunity, we also discovered a niche associated with inflammation and stress, including fibroblast and cardiomyocyte populations in the healthy heart, which are expanded in cardiomyopathies^27,85^. We define this as a myocardial-stress niche given it is rich in pro-fibrogenic and inflammatory signalling signatures. Circulating inflammatory cytokines, including IL-6 and IL-1β, are elevated in heart failure and directly relate to disease severity^105^. Moreover, clinical trials showed benefit for Canakinumab in heart failure, suggesting a causal role for IL-1β in its pathogenesis^106^. Although recent studies pointed to cytokine-mediated fibroblast activation^107^ and expansion of stressed CMs in ischemia^108^, the precise cellular mechanisms responsible for fibroblast activation or CM transition to a stressed status remains unclear. Our finding of inflammatory cytokine receptor expression in the vCM3_stressed cells provides a potential mechanism mediating CM susceptibility to inflammation, remodelling, and death. vCM3_stressed expresses *NPPB* encoding the peptide hormone BNP^109^, as similarly reported in a recent study of ischemic heart^108^. This is detectable at low levels in healthy individuals but is markedly elevated in response to increased wall stress in heart failure states^26^, being used as a clinical biomarker which is reduced by successful treatment^109^. This is consistent with our observation that the vCM3_stressed population can be found in the healthy heart, but is expanded in cardiomyopathy. Tackling the pro-inflammatory signalling received by vCM3_stressed may be a promising target for future heart failure therapies.

In order to identify targets and mechanisms of action of drugs, we developed drug2cell and demonstrated its potential by identifying novel putative cardioactive effects of non-cardiac medications. Specifically, based on their expression profile, drug2cell highlighted P cells as unexpected cellular targets for non-cardiac drugs with chronotropic effects (the GLP1 analogues and Perampanel), the mechanisms of which had remained elusive^75,110,111^. Currently drug2cell does not account for effects such as physiological antagonism (i.e. agonism of an inhibitory receptor causing downstream inhibition) and therefore predicts drug effect magnitude rather than direction, additionally there is scope to expand its predictions by coupling it to databases of intracellular signalling modules^112^. In the future, applying drug2cell to whole body cell atlas datasets will improve *in silico* screening by predicting adverse effects in cell types across all organs.

Covering eight anatomical regions, our analysis represents the largest (>700,000 single cells plus ATAC and spatial data) and most finely annotated healthy human heart cell atlas to date, shedding light on novel cell states with implications in disease. Crucially, it provides a framework to combine multimodal data and to integrate knowledge-based and unsupervised structural annotations in order to spatially resolve cellular niches. This dataset, together with the new CellPhoneDB neural-GPCR module and drug2cell pipeline, will be valuable resources for the community going forwards.

## Methods

### Research ethics for donor tissues

All heart tissue was obtained from transplant donors after Research Ethic Committee approval and written informed consent from donor families as previously described^1^. Ethical approvals for additional heart tissue were: Donor A61 (REC ref 15/EE/0152, East of England Cambridge South Research Ethics Committee), AH1 (DN_A17), AH2 (DN_A18), AH5 (DN_A19), AH6 (DN_A20) (REC ref 16/LO/1568, London - London Bridge Research Ethics Committee), AV1 (HOPA03), AV3 (HOPB01), AV10 (HOPC05), AV13 (HOPA05), AV14 (HOPA06) (REC ref 16/NE/0230, North East - Newcastle & North Tyneside Research Ethics Committee). Failing hearts samples used for validation were obtained under the ethical approval given to the Royal Brompton & Harefield Hospital Cardiovascular Research Centre Tissue Bank (REC ref: 19/SC/0257).

### Tissue acquisition and processing

Cardiovascular history was unremarkable for all donors (**Supp. Table 1**). Hearts contributing to the SAN and AVN regions were from donors confirmed to be in sinus rhythm with normal conduction parameters on ECG prior to donation (**Supp. Table 1**). Hearts were acquired after circulatory death (DCD) (D61 and AH1) and after brain death (DBD) (HOPA03). For DCD donors after confirmation of death there was a mandatory five-minute stand-off prior to sternotomy. In all cases the aorta was cross-clamped and cold cardioplegia solution was administered to the aortic root prior to cardiotomy. HOPA03 was procured in the standard fashion and then immediately preserved with and transported on a hypothermic perfusion machine. It then underwent 4 hours of normothermic perfusion before samples were taken. For single nuclei sequencing, all donor samples were full-thickness myocardial tissues from the SAN, AVN, left and right atrium, left and right ventricles, interventricular septum and apex. For spatial transcriptomics, ventricle regions, whose thickness were larger than one side of the Visium frame (6 mm), were separated into epicardial and endocardial portions. Samples used for single nuclei isolation and spatial transcriptomics were flash-frozen or frozen in OCT and stored at −80 °C, or formalin-fixed and subsequently embedded in paraffin blocks. All tissues were stored and transported on ice at all times until freezing or tissue dissociation to minimise any transcriptional degradation.

### Single nuclei isolation

Single nuclei were obtained from flash-frozen tissues using sectioning and mechanical homogenization as previously described ^1,113^. 5-10 mm thickness frozen tissues were first sectioned with cryostat in a 50 um thickness section. All sections from each sample were homogenised using a 7 ml glass Dounce tissue grinder set (Merck) with 8–10 strokes of a loose pestle (A) and 8–10 strokes of a tight pestle (B) in homogenization buffer (250 mM sucrose, 25 mM KCl, 5 mM MgCl2, 10 mM Tris-HCl, 1 mM dithiothreitol (DTT), 1× protease inhibitor, 0.4 U μl−1 RNaseIn, 0.2 U μl−1 SUPERaseIn, 0.1% Triton X-100 in nuclease-free water). Homogenate was filtered through a 40-μm cell strainer (Corning). After centrifugation (500g, 5 min, 4 °C) the supernatant was removed and the pellet was resuspended in storage buffer (1× PBS, 4% bovine serum albumin (BSA), 0.2U μl−1 Protector RNaseIn). Nuclei were stained with 7-AAD Viability Staining Solution (BioLegend) and positive single nuclei were purified by fluorescent activated cell sorting (FACS) using MA900 Multi-Application Cell Sorter (Sony). Nuclei purification and integrity was verified under a microscope, and nuclei were further processed for multiome paired RNA and ATACseq using the Chromium Controller (10X Genomics) according to the manufacturer’s protocol.

### Chromium 10X library preparation

Single nuclei were manually counted by Trypan blue exclusion. Nuclei suspension was adjusted to 1000–3,000 nuclei per microlitre and loaded on the Chromium Controller (10X Genomics) with a targeted nuclei recovery of 5,000–10,000 per reaction. 3’ gene expression libraries and ATAC libraries were prepared according to the manufacturer’s instructions of the Chromium Single Cell ATAC and Multiome ATAC+Gene Expression Kits (10X Genomics). Quality control of cDNA and final libraries was done using Bioanalyzer High Sensitivity DNA Analysis (Agilent) or 4200 TapeStation System (Agilent). Libraries were sequenced using NovaSeq 6000 (Illumina) at Wellcome Sanger Institute with a minimum depth of 20,000–30,000 read pairs per nucleus,

### Visium slides and library preparation

Fresh-frozen (FF) samples: Fresh samples were frozen and embedded in optimal cutting temperature medium (OCT) using a dry ice-cooled bath of isopentane at −45°C. OCT-embedded samples were sectioned using a cryostat (Leica CX3050S) and were cut at 10 μm. Formalin-fixed paraffin embedded (FFPE) samples: Fresh samples were fixed in >5 times their volume of 4% v/v formalin at ambient temperature for 24 hours before processing to paraffin on a Tissue-Tek Vacuum Infiltration Processor 5 (Sakura Finetek). FFPE blocks were sectioned at 5um using a microtome (Leica RM2125RT).

Samples were selected on the basis of morphology with expert review (YH), orientation (based on HE) and either RNA integrity number (fresh-frozen samples) or DV200 (formalin-fixed) that was obtained using Agilent 2100 Bioanalyzer. In addition, FFPE tissues were checked for possible detachment issues using the 10x Genomics Adhesion test slides. FFPE Visium Spatial Gene Expression (10x genomics) was performed following the manufacturer’s protocol. For FF samples, the Tissue Optimization protocol from 10x Genomics was performed to obtain a permeabilisation time of 45 min, and the FF Visium Spatial Gene Expression experiment was performed as per the manufacturer’s protocol (10x Genomics). H&E stained Visium Gene Expression slides were imaged at 40× on Hamamatsu NanoZoomer S60. After transcript capture, Visium Library Preparation Protocol from 10x Genomics was performed. Eight cDNA libraries were diluted and pooled to a final concentration of 2.25 nM (200 μl volume) and sequenced on 2× SP flow cells of Illumina NovaSeq 6000.

### Read mapping

After sequencing, samples were demultiplexed and stored as CRAM files. Each sample of single-nuclei (sn) RNA-seq was mapped to the human reference genome (GRCh38) provided by 10X Genomics, and using the CellRanger software or STARsolo with default parameters. For single-nuclei samples, the reference for pre-mRNA, was created using the 10X Genomics (https://support.10xgenomics.com/single-cell-gene-expression/software/pipelines/latest/advanced/references). Each sample of Multiome, or Visium were mapped to the human reference genome (GRCh38) provided by 10X Genomics, and using the CellRanger ARC (v.2.0.0) or SpaceRanger (v.1.1.0) with default parameters. For Visium samples, SpaceRanger was also used to align paired histology images with mRNA capture spot positions in the Visium slides. Part of the SAN samples were mixed with different donors after the nuclei isolation for cost-efficient experimental design (**Supp. Table 6**) and computationally demultiplexed (Soupercell, v2.0) ^114^ based on genetic variation between donors.

### Quality control and processing of suspension data

For scRNA-seq, snRNA-seq, and Multiome gene expression data, the CellBender algorithm ^115^ (remove-background) was applied to remove ambient and background RNA from each count matrix produced by CellRanger pipeline. Scanpy (v.1.7.1) was used for the quality control and processing. Cells or nuclei for each sample were filtered for more than 200 genes and less than 20% or 5% mitochondrial reads, respectively, and genes were filtered for expression in more than three cells. A Scrublet ^116^ (v.X) score cut-off 0.3 of was applied to remove doublets. The Scanpy toolkit was used to perform downstream processing, including normalisation, log transformation, variable gene detection, and regressing out unwanted sources of variation (n_counts and percent_mito).

For Multiome ATAC data (10x Genomics), the data processed with CellRanger-ARC were further analysed using ArchR ^28^. QC was performed, considering among others TSS enrichment, nucleosomal banding patterns, the number and fraction of fragments in peaks, reads falling into ENCODE blacklist regions as well as doublet scores computed by ArchR. For high-quality cells, reads were mapped to 500-bp bins across the reference genome (hg38) followed by dimensionality reduction using latent semantic indexing (LSI), batch correction and cell-clustering with a modularity optimization-based approach implemented in Seurat ^117^. Gene scores based on the chromatin accessibility around genes were computed from the tile matrix in ArchR to check their consistency with measured expression values.

### Data integration and cell-type annotation

All transcriptome data were integrated by using scVI ^11^ (v.0.14.5, n_hidden=128, n_latent=50, n_layers=3, dispersion=’gene-batch’) and scArches (v.0.5.5, n_hidden=128, n_latent=50, n_layers=3, dispersion=’gene-batch’) with correcting batch effects (donor, cells-or-nuclei, and 10X Genomics library generation kits) and removing unwanted source of variations (total counts, percent mitochondrial genes, and percent ribosomal genes for ‘continuous_covariate_keys’). Scanpy functions were used to compute a neighborhood graph of observations based on the scVI latent space (scanpy.pp.neighbors) and to perform dimensionality reduction (sc.tl.umap) and Leiden clustering (sc.tl.leiden, resolution=1.0). Clusters showing hybrid transcriptional signatures which also had high scrublet-score were removed. After re-clustering, cell lineages were annotated on the basis of major maker gene expressions and statistically identified marker gene expression for each cluster (sc.tl.rank_genes_groups).

To identify fine-grained cell states of the cardiac conduction system (CCS), atrial cardiomyocytes (CMs) of SAN were sub-clustered; thus we identified a cluster of pacemaker cells (SAN pacemaker cell; SAN_P_cell) which expresses canonical channel genes (*HCN1*, *HCN4*) ^7^ and transcription factor (*TBX3*)(**Fig. 1C, Supp. Fig. 3A**). Sub-clustering of AVN atrial and ventricular CMs showed two CCS cell state clusters (AVN pacemaker cell; AVN_P_cell and AV bundle cell; AV_bundle) (**Fig. 1D**). AV bundle cells formed a distinct cluster defined by enrichment *GJA5* (encoding the Cx40), *CRNDE*, *CNTN5*, which were previously identified as a marker of His bundle cells in the mouse heart^14^ (**Supp. Fig. 3C**). To identify Purkinje cells, cardiomyocyte populations from AVN samples were integrated and clustered with those from the apex. This showed one cluster which contained not only the CCS cells from the AVN, but also a population derived from the apex expressing Purkinje fibre marker genes (*GJA5, IRX3, KCNJ3, MYL4*)(**Fig. 1E, Supp. Fig. 3D,E**)^17^.

Cell states of other cell types and other regions were defined by label transfer^19^ using the published dataset^1^ as a reference with revised annotations. The new annotations include NC populations, which express pan-glial markers and lack core neuronal markers (**Supp. Fig. 3H**); therefore this compartment is best described as glia and will be described below with the _glial suffix. ‘FB4’ was renamed as ‘FB4_activated’ based on FB activation signature genes (*POSTN*, *TNC*) and genes encoding ECM proteins (*COL1A1*, *COL1A2*, *COL3A1, FN1*) (**Fig. 1G, Supp. Fig. 3I**). ‘vCM3’ was renamed as ‘vCM3_stressed’ based on the specific expression of *NPPB* which encodes B-type natriuretic peptide (BNP), which is a diagnostic marker for heart failure and valuable prognostic predictors for the entire spectrum of the disease severity and expressed in stressed cardiomyocytes ^26^ (**Fig. 1G, Supp. Fig. 3J**). ‘EC7_atria’ was renamed as ‘EC7_endocardial’ based on a recently published study^27^. For myeloid cells, dimensionality reduction and batch correction (scVI) with Leiden clustering were repeated to identify and annotate fine-grained cell states (**Supp. Fig. 3K,L**). The transferred cell state labels which were not consistent with the coarse-grained cell type labels based on the global clusters were replaced with ‘unclassified’ and ignored in the downstream analysis.

### Cell type/state label transfer to snATAC-Seq and multimodal data integration

We identified cell types and states in snATAC-Seq cells based on the sc/snRNA-Seq annotation and using multiome data as a bridge dataset. We applied the Signac package (Stuart et al, Nature 2021) for data processing and label transfer. We performed normalisation using SCTransform and reduced dimensionality with PCA. We processed DNA accessibility assays with latent semantic indexing (LSI). We utilised reciprocal LSI projection to integrate anchors between the multi-ome and the snATAC-Seq datasets. We integrated the low-dimensional cell embeddings (LSI coordinates) across the datasets. We employed the multi-ome dataset as a reference and mapped the snATAC-Seq dataset to it using the FindTransferAnchors() and MapQuery() functions from Seurat (Hao et al, Cell 2021). We integrated all modalities into a merged low-dimensional embedding by applying the MultiVI package (Ashuach, Biorxiv 2022) and annotated the cells as described above. We confirmed the agreement between the Leiden clustering of the multimodal dataset and the cell annotations using metrics to assess clustering performance (AMI, homogeneity, completeness, Fowlkes-Mallows scores).

### Spatial mapping of cell states with cell2location

To spatially map heart cell states defined by single-cell transcriptomic data analysis in the Visium data, we used our cell2location method^118^. Briefly, we first estimated reference signatures of cell states per each region using a negative binomial regression model provided in the cell2ocation package. For the cell states which had low cell/nuclei numbers per region (fewer than 100), we used the cells from all regions for better signature inference. The inferred reference cell state signatures were used for cell2location cell type mapping that estimates the abundance of each cell state in each Visium spot by decomposing spot mRNA counts. The H&E images of the Visium slides were used to determine the average number of nuclei per Visium spot (=7) in the tissue and used as a hyperparameter in the cell2location pipeline. Cell state proportions in each Visium spot were calculated based on the estimated cell state abundances.

### Cell state spatial enrichment analysis

Anatomical microstructures of spatial transcriptomic data were manually annotated using the paired histology images as follows: epicardium, subepicardium, endocardium, myocardium, vessel, nerve, adipose tissue, cardiac_skeleton, fibrosis, node and AV bundle. Cell state proportions per spot were calculated based on the estimated abundances of cell states (cell2location). Cell state enrichments (odds ratio) in each structure were calculated by dividing the odds of target cell state proportions by the odds of the other cell state proportions. Odds of cell proportions were calculated as the ratio of cell proportion in the spots of a structure of interest to that in the other spots. Statistical significance was obtained by chi-square analysis (scipy.stats.chi2_contingency) and the *p*-value was corrected with Benjamini-Hochberg method.

### Cellular microenvironment discovery

The NMF analysis implemented in the cell2location pipeline was performed on spatial mapping results of each anatomical region of the heart. The NMF model was trained for a range of cell state combinations (number of factors: n_fact) N={5,…,14} and the effect size of the cell state group abundance between the spots within a given structure against the spots in the other areas was calculated for each factor (95 factors in total)(**Supp. Fig. 7, step1**). To test the significance, we permuted the annotation labels of all spots and generated a null distribution of the effect size. The *P* values were calculated on the basis of the proportion of the value that is as high as or higher than the actual effect size. For each given structure, first we selected the factor which has the highest significant effect size (*best-factor*)(**Supp. Fig. 7, step2**). Next we selected the n_fact which had multiple numbers of factors (*fine-factors*) with an effect size more than an arbitrary proportion (0.5) of the *best-factor* (**Supp. Fig. 7, step3**). We can consider the *fine-factors* as refined microenvironments which were identified with the NMF method (and not with the knowledge-based structural annotations).

For the myocardial-stress niche, first the estimated abundances (cell2location) of FB4_activated and vCM3_stressed were multiplied. Based on the multiplied abundance, clusters of neighbouring spots (n>5) which had higher than a value (0.03) were selected.

### CellPhoneDB NeuroGPCR expansion module

Using the HGNC library of GPCRs as a master list (HGNC group 139) we used publicly available databases (UniProt, Reactome, IUPHAR and GPCRdb (https://gpcrdb.org/)^119^ to generate a set of GPCRs with known ligands. In order to generate Ligand-Receptor interactions for these GPCRs we used ligand genes (for gene-encoded ligands). For non-gene-encoded ligands (such as small molecule ligands) we used ligand-proxies in the form of specific biosynthetic enzymes or transporter genes. Additionally, we added new trans-synaptic adhesion molecule interactions ^57,120,121^. Together this formed over 800 new interactions (**Supp. Table 3,4**) which we used with CellPhoneDB’s user-defined ‘database generate’ function ^8^.

### Spatially resolved cell-cell interaction analysis

CellPhoneDB analyses with a custom GPCR/neuronal ligand-receptor database were performed on the identified niches and the cell state components. Overall, the cell-cell interaction inference were performed using single-cell transcriptomic data of each anatomical region and by restricting to the cell states which were co-localised in the identified cellular niches.

For CCS and myocardial-stress niches, we selected the cell states that were either in the node of SAN or AVN and retrieved the interacting pairs of ligands and receptors satisfying the following criteria: (1) all the members were expressed in at least 10% of the cells in the cell states and (2) ligand-receptor complexes specific to two cell states inferred by the statistical method framework in CellPhoneDB (‘statistical_analysis’, P-value threshold=0.05).

For epicardial-subepicardial niches, the ligand-receptor interactions of the co-locating cell states were retrieved based on the following criteria: (1) all the members were expressed in at least 10% of the cell states and (2) at least one of the members in the ligand or the receptor is a differentially expressed gene compared with other cell states (SCANPY sc.tl.rank_genes_groups() function, P-value threshold=0.05, Fold change threshold=0.1). The ligand-receptor interactions were further selected based on mean expressions and the biological questions as indicated in the Result section and the figure legends.

The cell states used in each analysis: SAN (SAN_P_cell, FB2, FB4_activated, FB5, FB6, NC2_glial_NGF+, LYVE1+IGF1+MP), AVN (AVN_P_cell, aCM2, FB1, FB2, FB5, SMC1_basic, SMC2_art, NC2_glial_NGF+, LYVE1+IGF1+MP, Mast), epicardial-subepicardial niche (Meso, EC6_ven, EC8_ln, PC2_atria, LYVE1+IGF1+MP, B_plasma, T/NK_cycling, FB2, FB3, FB5, NC1), and myocardial-stress niche (FB3, FB4_activated, FB5, vCM3_stressed, EC2_cap, EC3_cap, EC4_immune, EC6_ven, PC3_str, SMC2_art, LYVE1+IGF1+MP, MoMP, NK_CD16hi).

### Ion channel and GPCR profile

Differential gene expression analysis with t-test method was performed using the SCANPY sc.tl.rank_genes_groups() function. Only multiome gene expression data was used to avoid the technical batch effects from kit differences. P value correction was performed using the Benjamini–Hochberg method. Each of the CCS cells (SAN_P_cell, AVN_P_cell, AV_bundle_cell, and Purkinje cells) was compared with non-CCS aCMs as a reference (**Supp. Table 2**). Genes were deemed DE with an adjusted P value of <0.05. DEGs encoding for ion channels and GPCRs were selected based on HGNC groups 177 and 139, respectively. Upregulated (log_2_FC>0) DEGs were depicted in the GPCR overview schematic (**Fig. 3B**).

### Mouse-Human DEG comparison

We took the ion channel genes which were differentially expressed in human SAN_P_cells (compared to atrial cardiomyocytes), and plotted differential expression data (log_2_FC, adjusted p value) for those alongside the orthologous ion channel gene from two mouse single cell studies where it was available ^38,39^. Orthologs were identified using NCBI Orthologs database.

### Identification of ligands in Visium data

Four spatial transcriptomics slides were identified as containing the right atrial ganglionated plexus (RAGP) by an expert anatomist (YH). Spot counts from the Visium-FFPE slides which contain RAGP were normalised, and spots were scored for expression of four generic pan-neuronal cytoskeletal markers (*PRPH*, *NEFL*, *NEFM*, *NEFH*) using the SCANPY sc.tl.gene_score() function. Correlation of individual gene expression profiles with this score was calculated (Pearson r and p value for each gene). The ligand/ligand-proxy list created as part of the CellPhoneDB NeuroGPCR module was used to identify ‘Ligand’ genes amongst the set of correlated genes.

### Gene regulatory network

The Scenic pipeline (Aibar et al., 2017; Van de Sande et al., 2020) was used (pySCENIC version 0.11.2) to predict transcription factors (TFs) and putative target genes regulated in the P cells. First, gene regulatory interactions were calculated based on co-expression across the single cell transcriptome data sets of atrial cardiomyocytes (using only Multiome data) with GRNBoost2 (Moerman et al., 2019). This was followed by pruning interactions using known TF binding motifs, and the construction of dataset-specific regulatory modules (regulons) (Imrichová et al., 2015). Regulons were then scored in each individual cell using AUCell. P cell relevant TFs and target genes were retrieved based on the following criteria: (1) regulator TFs which are differentially expressed genes in P cells compared with other atrial CMs (‘scanpy.tl.rank_genes_groups’ with only Multiome data, P-value threshold=0.05, Fold change threshold=0.5) and (2) target genes which were expressed in at least 10% and differentially expressed in P cells compared to other atrial CMs (same criteria as TF selection). A network of regulatory TFs and target genes was then constructed by linking individual regulons to create a graph (NetworkX, v.2.6.3)(**Fig. 3C**). The node colour of the target genes are based on the class: GPCR (HGNC, GID139), ion channel (HGNC, GID177), or TFs^122^.

The interactions of regulatory TFs and target genes were also inferred by using the ATAC data and ArchR (v.1.0.2)^28^. Prior to peak calling, pseudo-bulk replicates were generated (*addGroupCoverages*) for each fine-grain cell state annotated based on the gene expression data. Peak calling was performed for each cell state and the peak sets were merged to obtain a unified peak set (*addReproduciblePeakSet*). TF-binding motifs in the identified peaks were searched (*addMotifAnnotations*, motifSet=“cisbp”), and correlations between peak accessibility and gene expression were analysed (*addPeak2GeneLinks* and *getPeak2GeneLinks*, correlation > 0.4, FDR < 1e-04). TFs and potential target gene interactions were obtained by combining the two results (**Supp. Fig. 12B**). The inferred interactions are highlighted in red in the network graph (**Fig. 3C**).

### Drug2cell

Drug and target genes information were obtained from ChEMBL (version 30). Drugs were filtered based on targeting organisms (Homo sapiens), achieved phase in a clinical trial (max_phase=4, clinically approved), and functionally active or not. The activity (pChEMBL) threshold was specifically set for each family of target molecules according to the IDG project (https://druggablegenome.net/ProteinFam) (Kinases: ≦ 30nM, GPCRs: ≦ 100nM, Nuclear Hormone Receptors: ≦ 100nM, Ion Channels: ≦ 10μM, Others: ≦ 30nM) (**Supp. Table 7**). Clinically approved drugs are categorised based on the WHO ATC classification (https://www.who.int/tools/atc-ddd-toolkit/atc-classification). Drug scores in each single cell were calculated based on the target gene expressions. Score can be obtained by taking the mean of a set of target genes either without (method=’mean’) or with (method=’seurat’) subtracting with the mean expression of a reference set of genes (Seurat ref). The reference set is randomly sampled from the gene pool for each binned expression value. For the drug repurposing analysis, all the drugs are tested and select ones which have the statistically highest score in a cell type of interest by testing with wilcoxon sum test and p value were adjusted using the Benjamini–Hochberg method. For the drugs targeting GPCRs or ion channels, we searched the clinically approved (Max Phase: 4) and preclinical bioactive molecules with drug-like properties (Max Phase: 1-3) which target genes encoding GPCRs (HGNC GID:139) or ion channels (HGNC GID:177). The drug2cell python package is available at (https://github.com/Teichlab/drug2cell).

### Single-molecule fluorescence *in situ* hybridization

The FFPE-embedded heart tissue was sectioned onto SuperFrost Plus slides at a thickness of 5 μm. Staining with the RNAscope Multiplex Fluorescent Reagent Kit v2 Assay (Advanced Cell Diagnostics, Bio-Techne) was automated using a Leica BOND RX, according to the manufacturers’ instructions. The tissues were baked and dewaxed on the Leica Bond RX, followed by the application of a heat induced epitope retrieval (HIER) step with epitope retrieval 2 (ER2) for 15 min at 95℃ and protease digestion with Protease III for 15 min. Subsequent processing included RNAscope probe hybridization and channel development with Opal 520, Opal 570, and Opal 650 dyes (Akoya Biosciences) at a concentration of 1:1000, and streptavidin-conjugated Atto-425 (Bio Trend) at a concentration of 1:400 using TSA-biotin (TSA Plus Biotin Kit, Perkin Elmer). All nuclei were DAPI-stained. Stained sections were imaged with a Perkin Elmer Opera Phenix High-Content Screening System, using a 20× water-immersion objective (NA 0.16, 0.299 μm per pixel). Channels: DAPI (excitation 375 nm, emission 435–480 nm), Opal 520 (ex. 488 nm, em. 500–550 nm), Opal 570 (ex. 561 nm, em. 570–630 nm), Opal 650 (ex. 640 nm, em. 650–760 nm) and Atto 425 (ex. 425 nm, em. 463-501 nm). Stained sections were also imaged on the Hamamatsu S60 using a 40x objective (0.23 μm per pixel).

### RNA scope quantification

Quantification of RNAscope images was performed using imageJ. In order to remove background, each channel was initially normalised by: 1) subtracting the raw image with a Gaussian blur transformation with σ=50; 2) Performing a background subtraction with rolling ball radius of 50 pixels; 3) Set every pixel with an intensity lower than 40 (of an 8-bit image) to 0. Following normalisation, each section was quantified with sequential Regions of Interest (ROI) of 200×200μm. To avoid any bias due to the placement of the initial ROI, quantification was performed over 3 rounds, with the initial ROI displaced by 50μm in the x and y axis in each round. The following parameters were recorded per ROI: number of nuclei; *COL1A1* area; *NPPB* area. For data analysis, in order to avoid variation due to cell density, for each ROI, *COL1A1* and *NPPB* areas were normalised by dividing the area value by the number of nuclei in the ROI. Only ROIs with more than 10 nuclei were considered for analysis. An ROI was only considered to contain be a *COL1A1*/*NPPB* niche, if the staining area was equal to or higher than 1μm^2^. To avoid any confounding effect due to different section sizes, we quantified the increase of the number of *COL1A1*/*NPPB* niches via the Average ROI percentage, consisting of the number of *COL1A1*/*NPPB* niche ROI divided by the total number of ROI, averaged over the 3 rounds of quantitation. To quantify the expression of *COL1A1* and *NPPB*, we averaged the normalised area of each ROI per niche over the 3 rounds of quantitation.

### Immunofluorescence

The FFPE-embedded heart tissue samples were sectioned at 6 µm thickness. The sections were immersed in Xylene for 2×10 minutes, 100% ethanol for 2×10 minutes, 95% ethanol for 5 minutes, 70% ethanol for 5 minutes, 50% ethanol for 5 minutes, and incubated in deionized water until use. Antigen retrieval was performed using the proteinase K kit from abcam for 5 minutes at RT. (ab64220). Following antigen retrieval, sections were permeabilized and blocked in 0.1M TRIS, containing 0.1% Triton (Sigma), 1% normal mouse serum, 1% normal goat serum and 1% BSA (R&D Systems). Samples were stained for 2h at RT in a wet chamber with the appropriate antibodies, washed 3 times in PBS and mounted in Fluoromount-G® (Southern Biotech). Images were acquired using a TCS SP8 (Leica, Milton Keynes, UK) inverted confocal microscope, with a 40x 1.1NA water objective. Raw imaging data were processed using Imaris (Bitplane). The staining set-up and antibody information is in Supplementary Table.

## Acknowledgements

This paper is dedicated to the memory of our dear friend and colleague Dr. Daniele Muraro who contributed to this work. The authors extend their deep gratitude to the donors and relatives who provided cardiac tissues for research. We thank NHS Blood and Transplant, the Cambridge Biorepository for Translational Medicine (CBTM) and the Cardiovascular Research Centre Biobank at the Royal Brompton and Harefield hospitals. We thank the Facility for Imaging by Light Microscopy and S. Rothery and L. Lawrence from the Research Histology Facility at Imperial College. We gratefully acknowledge support from the Wellcome Sanger Cytometry Core Facility, Cellular Genetics Informatics team, Cellular Generation and Phenotyping (CGaP) team, Core DNA Pipelines team, and Imperial College Healthcare NHS Trust Tissue Bank. Vitalii Kleshchevnikov for his help on cell2location analysis. Kevin Troulé Lozano and Luz Garcia-Alonso for their help on CellPhoneDB neural-GPCR module curation. This work was made possible by a partnership between the Wellcome Sanger Institute, Imperial College London, and Newcastle University.

## Funding

This project has been made possible in part by the Wellcome Trust (WT206194, S.A.T); Chan Zuckerberg Foundation (2019-002431, 2021-237882) to M.N., and S.A.T.; the British Heart Foundation and Deutsches Zentrum für Herz-Kreislauf-Forschung (BHF/DZHK: SP/19/1/34461) to M.N., and S.A.T.; the National Institute for Health and Care Research (NIHR) Blood and Transplant Research Unit in Organ Donation and Transplantation (NIHR203332), a partnership between NHS Blood and Transplant, University of Cambridge and Newcastle University, to J.D., and L.W.; the Wellcome Trust Clinical PhD Fellowship to J.C.; the Overseas Research Fellowship of the Takeda Science Foundation to K.K.; the National Heart and Lung Institute PhD studentship to S. N. B.; the British Society for Heart Failure Research Fellowship to L.M. and Alexander Jansons Myocarditis UK to L.M. Imperial College Healthcare NHS Trust Tissue Bank is funded by the National Institute for Health Research (NIHR) Biomedical Research Centre.

## Author contributions

Project design: K.K., J.C., M.N., S.A.T.; Data generation: K.K., J.C., A.M.A.M., M.L., L.R., L.M., M.D., N.R., S.N.B., S.P. (Shani Perera), A.W.C., I.M., K.T.M., L.B., L.M., L.T., L.W., M.M.H., S.P. (Sophie Pritchard), J.D., K.S.P., R.A.C., M.N.; Structural annotation: J.C., S.Y.H.; Data analysis: K.K., J.C., D.M., A.M.A.M., J.P.P., N.K., K.P., C.T.L.; drug2cell package: K.P., K.K.; CellPhoneDB neural-GPCR module: J.C.; Web portal: M.P.; Writing: K.K., J.C., M.N., S.A.T.; Review and editing: A.M.A.M., J.P.P., C.T.L., S.P., I.M.; Supervision: J.D., K.S.P., M.P., M.R.C., N.H., M.N., S.A.T.

## Competing interests

In the past three years, S.A.T. has consulted or been a member of scientific advisory boards at Roche, Genentech, Biogen, GlaxoSmithKline, Qiagen and ForeSite Labs and is an equity holder of Transition Bio. The remaining authors declare no competing interests.

## Data availability

Sequencing data for Multiome and Visium will be available on Human Cell Atlas Data Portal. Processed data of sc/snRNAseq and Visium data will be available for browsing gene expression and download via heartcellatlas.org.

---

Supp. Fig. 1 Dissection protocols for cardiac conduction system Workflow to capture SAN and AVN regions using a two-stage dissection. For the SAN, a region of the posterolateral RA parallel to and spanning the *crista terminalis* (CT) was isolated (dashed box). For the AVN and AV bundle, a single region incorporating the Triangle of Koch (bordered by Todaro’s tendon, the coronary sinus ostium and tricuspid valve annulus) as well as the basal septum (spanning from interatrial to interventricular septum, and including the membranous septum) was isolated (dashed box). These were then subdivided into several smaller samples each ∼4mm thick, which were individually frozen and embedded in OCT. Cryosections were stained (HE) with expert histologist review to determine the presence of SAN, AVN and AV bundle (yellow bordered zone). CCS-containing blocks (green arrows) then proceed to Multiome, Visium and RNAscope. **Abbreviations:** SAN (sinoatrial node); AVN (atrioventricular node); RA (right atrium); SVC (superior vena cava); IVC (inferior vena cava); MS (membranous septum); TV (tricuspid valve); CCS (cardiac conduction system).

Supp. Fig. 2 Profiling of cardiac cells A,B: UMAP representation of gene expression data from the eight regions showing major batch keys: ‘donor’, ‘cell_or_nuclei’, and ‘kit_10x’ labels (A) or highlighting each region (B). C: Major marker expression in the identified 12 cell types. D: UMAP representation of gene expression data from the eight regions showing the 75 fine-grained cell state labels. **Abbreviations:** CCS (cardiac conduction system); SAN (sinoatrial node); AVN (atrioventricular node); AVB (AV bundle); EPI (epicardium); ENDO (endocardium); LV (left ventricle); RV (right ventricle); SP (septum); AX (apex); CM (cardiomyocyte); FB (fibroblast); EC (endothelial cell); SMC (smooth muscle cell); P_cell (pacemaker cell)

Supp. Fig. 3 Identification of fine-grained cell states A: Dot plot showing the marker gene expressions of SAN and AVN P cells. B: UMAP representation of gene expression data of AVN atrial and ventricle CMs. Dimensional reduction and leiden clustering were repeated for atrial CMs and CCS cell clusters (colored in black) (Fig. 1D). C: Dot plot showing the marker gene expressions of AV bundle cells. D: UMAP representation of gene expression data of AX and AVN atrial and ventricle CMs. With the labels of leiden clusters, regions, and AVN CCS cells. E: Dot plot showing the marker gene expressions of Purkinje cells. F: Donor proportions of CCS cell. G: H&E morphology used of the spatial transcriptomic samples of the CCS (Fig. 1F) showing the SAN, AVN, AV bundle, and Purkinje cells. Histologically identified CCS regions are in dashed lines. H: Dot plot showing the expressions of glial, schwann, and neuronal cell marker genes in neural cell populations and other cell types. I: Dot plot showing the FB activation and extracellular matrix gene signature expressions in FB cell states. J: Dot plot showing the marker genes of cardiomyocyte stress in ventricular CM cell states. K,L: Fine-grained cell state annotation of myeloid compartment. UMAP representation of gene expression of myeloid cells from the eight regions (K). Cell states were annotated based on the clusters and myeloid marker expression (L).

Supp. Fig. 4 Profiling of cardiac cells with ATAC data A: UMAP representation of chromatin accessibility data from the eight regions showing the 60 fine-grained cell state labels. B: Dot plot showing the gene scores of differentially accessible genes in 12 major cell types. C: Dot plot showing the gene scores of the cardiomyocyte stress signature in ventricular CM cell states. D: Dot plot showing the gene scores of the marker genes of SAN and AVN P cells.

Supp. Fig. 5 Cell state enrichment in the manually annotated structures Cell state enrichments (odds ratio) in the ‘vessel’ (A), ‘fibrosis’ (B), ‘endocardium’ (C), and ‘nerve’ (D) of the free wall of the six regions: RA, LA, RV, LV, SP, and AX. Statistically significant enrichments (chi-square test, p<0.05) are shown in magenta-pink. Data show value ±SD.

Supp. Fig. 6 Cell state enrichment in the manually annotated CCS structures and the epicardium A: Cell state enrichment (odds ratio) in the CCS structures: ‘node’ of SAN and AVN, and ‘AV bundle’ of AVN. Statistically significant enrichments (chi-square test, p<0.05) are shown in magenta-pink. Data show value ±SD. B: Abundance of the cell states (mapped by cell2location) enriched in the ‘node’ of SAN. C,D: H&E image, manual structural annotations, and abundance of the cell states enriched in the ‘node’ (C) or ‘AV bundle’ (D) of AVN. E: Dot plot showing expression of connexin genes in MP and monocyte cell states from the SAN and AVN regions. F: Cell state enrichments (odds ratio) in the ‘epicardium-subepicardium’ of the free wall of the four regions: RV, LV, RA, and LA. Statistically significant enrichments (chi-square test, p<0.05) are shown in magenta-pink. Data show value ±SD.

Supp. Fig. 7 Decomposing manual structural annotations to identify cellular niches The steps of the similarity assessment of each factor with each manually annotated structure. Effect size (Cohen’s d) of the factor loadings of the spots in a given structure against spots in the other areas is calculated (step 1). For each given structure, the factor which has the highest significant effect size is selected (best-factor) (step2). Next, multiple numbers of factors within a ‘n_fact’ which had effect size more than an arbitrary proportion (0.5) of the best-factor (fine-factors) are selected (step3).

Supp. Fig. 8 Cellular niche identification in the node A, B_(i)_, C_(i)_: Selected factors which had high effect size in the ‘node’ of SAN visium sections (as indicated by the arrows): donor AH1 (A), AV14-slide 1 (B_(i)_), and AV14-slide 2 (C_(i)_). The factor names were assigned based on Fig. 2C, Supp. Fig. 8B_(iii)_, and Supp. Fig. 8C_(iii)_. B, C: Cellular microenvironment identification in the SAN. Manual structural annotations were performed based on the H&E images (B_(ii)_, C_(ii)_). Factors from cell2location NMF analysis which showed high similarity (Cohen’s d) with individual manual structural annotation were selected (as described in Methods). Factor loadings across locations (abundance of cell state group) are shown in spatial coordinates for selected factors (B_(iii)_, C_(iii)_). The accompanying dot plot illustrates factor loadings normalised per cell type (dot size and colour). Cell states with more than 0.4 in the selected factors are shown (B_(iii)_, C_(iii)_). Estimated abundance of representative cell states in the central node of SAN slide (B_(iv)_, C_(iv)_).

Supp. Fig. 9 Cellular localisations in the node A-D: Analyses on Visium-FFPE slides of the SAN region: HE image and manual structural annotations (A), cell state enrichments in the ‘node’ of SAN (B), and the proportions of the cell states (mapped by cell2location) enriched in the ‘node’ (C,D). Data in (B) show value ±SD. E: Expression of extracellular cellular matrix component genes which were differentially expressed in the central nodal niche compared with the peripheral niche. Each niche was defined based on the NMF analysis and the corresponding spots were selected by thresholding the factor loadings (top 10% of all spots). Differentially expressed genes were obtained by using a scanpy function (‘scanpy.tl.rank_genes_groups’ with visium spots of the nodes, P-value threshold=0.05, Fold change threshold=0.5).

Supp. Fig. 10: Epicardial cellular niches in the LA. A: Selected factors (n_fact = 5) which had high effect size in the ‘epicardium-subepicardium’ of RV. The factor names were assigned based on Fig. 2G. B: Selected factors (n_fact = 6) which had high effect size in the ‘epicardium-subepicardium’ of LA. The factor names were assigned based on Supp. Fig. 10D. C: Manual structural annotations based on the HE images. D: The factor loadings across locations (abundance of cell state group) are shown in spatial coordinates for the selected factors in (B). The accompanying dot plot illustrates factor loadings normalised per cell type (dot size and colour). Cell states with more than 0.4 in the selected factors are shown.

Supp. Fig. 11: Characterisation of human pacemaker cells and their niche. A: Dot plot comparing profiles of mouse and human SAN pacemaker cells. Publicly available differential expression data for mouse sinoatrial myocytes (Liang 2021, Linscheid 2019) were obtained and plotted with the DEGs encoding for ion channels in human SAN P cells. B: Dot plot shows the expression of the DEGs (log2FC>0, adj_pval<0.05) encoding for ion channels (HGNC GID:177) in any of the CCS cell states compared with the other aCMs. C: Dot plot shows the expression of the DEGs (log2FC>0, adj_pval<0.05) encoding for GPCRs (HGNC GID:139) in any of the CCS cell states compared with the other aCMs. D: Dot plot shows the expression of genes encoding for catecholamine β receptors. E,F: Inferred spatial cell-cell interactions of GPCRs (CellPhoneDB with neural-GPCR module) in the central node of SAN (E) or the ‘node’ structure of AVN (F), with P cell as the receiver. G: Visium plot showing *AGT* expression across a SAN region slide. **Abbreviations:** DEG (differentially expressed gene); GPCR (G protein-coupled receptors)

Supp. Fig. 12: Regulator TFs of P cells A: Dot plot showing regulator TF expressions in atrial CM cell states from multiome data. TFs were selected based on the differential expressions in P cells compared with other atrial CMs (Methods). B: Workflow for selecting regulator TF and target gene interactions. The network was constructed based on gene expression data and pySCENIC analysis. The interactions obtained from ATAC data were used for highlighting the interactions (Methods). C: Peak-to-gene linkage plot of *HCN1* (ArchR). One of the peaks linked to *HCN1* has FOXP2 binding motif as indicated.

Supp. Fig. 13: Cell-cell interactions in the node A: Inferred cell-cell trans-synaptic interactions (CellPhoneDB neural-GPCR module) in the AVN, spatially refined by cell states enriched in the ‘node’ structure, with AVN_P_cells as the receiver cells with expression receptors (LR mean > 1). B,C: Inferred spatial cell-cell interactions of LGICs in the central node of SAN (B) or the ‘node’ structure of AVN (C), with P cell as the receiver. **Abbreviations**: LR mean (mean expressions of the interacting ligand-receptor partners).

Supp. Fig. 14: Characterisation of human RAGP A: Right Atrial Ganglionated Plexus: Plot of ‘pan-neuronal cytoskeleton score’ (see Methods) in spatial coordinates matches RAGP (inset showing RAGP in associated HE image). B: Several GPCR ligands correlate with the pan-neuronal nerve cytoskeleton score. Correlation (Pearson r vs p value) of individual ligand genes with the pan-neuronal nerve cytoskeleton score (see Methods). Ligand genes with p value <0.05 and Pearson r > 0.1 are labelled. C: Plots of significantly correlated ligand genes in spatial coordinates. **Abbreviations**: RAGP (right atrial ganglionated plexus)

Supp. Fig. 15: Drug target exploration at the single-cell level Dotplot shows drugs which target GPCRs or Ion channels and genes encoding for their molecular targets. Dots pinpoint the drug-target pairs predicted in the P cells, and the dot colours show the maximum phase of the clinical trials for each drug. Target genes which are expressed in at least 10% of P cells are highlighted in red.

Supp. Fig. 16: An immune niche in the epicardium A: Abundance of the co-locating cell states (mapped by cell2location) in the ‘epicardium-subepicardium’ structure of RV (Fig. 2G). B: Dot plot showing expressions of IGH genes significantly enriched in the ‘epicardium-subepicardium’ compared with other manually annotated structures (‘scanpy.tl.rank_genes_groups’ with visium spots of structures, P-value threshold=0.05, Fold change threshold=0.5). C: Workflow of spatial CellPhoneDB analysis focusing on B_plasma cells. D: Inferred spatial cell-cell interactions of TGF-beta superfamily, spatially refined by niche partner cell states, with plasma B cells as the sender cell expressing ligands. E: Dot plot showing *TGFB1* expression in the cell states localised in the epicardial niches. F: Dot plot showing the expressions of ‘antimicrobial humoral response’ genes (Gene Ontology Term, GO:0019730) which were expressed significantly higher in mesothelial cells compared to other cells (‘scanpy.tl.rank_genes_groups’, log2FC>1, adj_pval<0.05). G: Expression of antimicrobial response genes, *SLPI* and *RARRES2*, in spatial transcriptomics data. **Abbreviations**: IGH (immunoglobulin heavy chain); LR mean (mean expressions of the interacting ligand-receptor partners).

Supp. Fig. 17: Ventricular myocardial-stress niche A,B: Comparisons of FB4_activated proportions amongst fibroblasts (A) and vCM3_stressed amongst ventricular cardiomyocytes (B) between control and diseased samples in publicly available diseased datasets (Chaffin *et al*., *Nature*, 2022; Reichart *et al*., *Science*, 2022). The cell state labels were transferred from the healthy dataset by using scNym (https://github.com/calico/scnym). *P < 0.05, **P < 0.01, and ***P < 0.001 (Wilcoxon rank-sum test with p-value collection). C: Single-molecule fluorescence *in situ* hybridization of left ventricular regions (LV, SP, and AX) stained with RNAscope probes against *NPPB* and *COL1A1* and DAPI. The proportions of the *COL1A1*/*NPPB* co-location and the expressions of each gene in the *COL1A1*/*NPPB* co-location were quantified for control (n=2 donors x 3 regions) and DCM samples (n=2 donors x 3 regions)(Methods). D,E: Visium plots of interventricular septum showing localisation of FB4_activated and vCM3_stressed (mapped by cell2location), a niche comprised of the two cell states (C), and expression of FB4_activated and vCM3_stressed markers *NPPB* and *COL1A1* (D). The myocardial-stress niche was defined based on the FB4_activated and vCM3_stressed abundance (Methods). F: Cell states enriched in the myocardial-stress niche, ordered by mean abundance. G,H: Inferred cell-cell interactions of cytokines (HGNC, GID:599,602,1932,781,1264) spatially refined by myocardial-stress niche cell states, with FB4_activated (G) or vCM3_stressed (H) as the receiver. I: Dotplot showing inflammatory cytokine receptor expressions in vCM3_stressed and other ventricular CMs of left ventricular regions (LV, SP, and AX). J: Myocardial-stress niche in the ventricle. TGFβ superfamily interactions from immune cells, such as MPs and NK cells, and vasculature cells to FB4_activated and vCM3_stressed may cause pathogenic fibrosis. Vasculature cells also express inflammatory cytokines, including IL-6, TNFSF12, that may directly affect ventricular CMs and lead to adverse cardiac remodelling. **Abbreviations**: vCM (ventricular cardiomyocyte); DCM (dilated cardiomyopathy); TGF (transforming growth factor)

Supp. Table 1: Anonymised Donor Information Anonymised donor information including clinical metadata

Supp. Table 2: DE test result of CCS cells Result of differential gene expression analysis of CCS cells (SAN_P_cell, AVN_P_cell, AV_bundle_cell, and Purkinje_cell) compared with the non-CCS aCM (t-test). P value correction was performed using the Benjamini–Hochberg method.

Supp. Table 3: CellPhoneDB neural-GPCR module, interactions input Table of CellPhoneDB neural-GPCR module interactions. Interaction partners use Uniprot ID (with protein name also listed) unless they are complexes.

Supp. Table 4: CellPhoneDB neural-GPCR module, complex input Table of complexes of CellPhoneDB neural-GPCR module interactions. Complex names are listed, with complex constituents in the same row (using Uniprot IDs as identifiers).

Supp. Table 5: Visium gene expression correlation with ‘pan-neuronal cytoskeleton score’ Correlation of genes with the pan-neuronal cytoskeleton score in visium data (Methods). Pearson r correlation coefficient, associated p value and mean expression across all visium spots.

Supp. Table 6: Information of the samples used in the analyses

Supp. Table 7: Filtered drugs and targets Drugs and target genes from ChEMBL (version 30) were filtered as described in the Methods.

## Notes

### Summary of Updates

Author list and affiliations updated.

