## Supplementary figures for "Spatially resolved multiomics of human cardiac niches"

Supp Fig 1

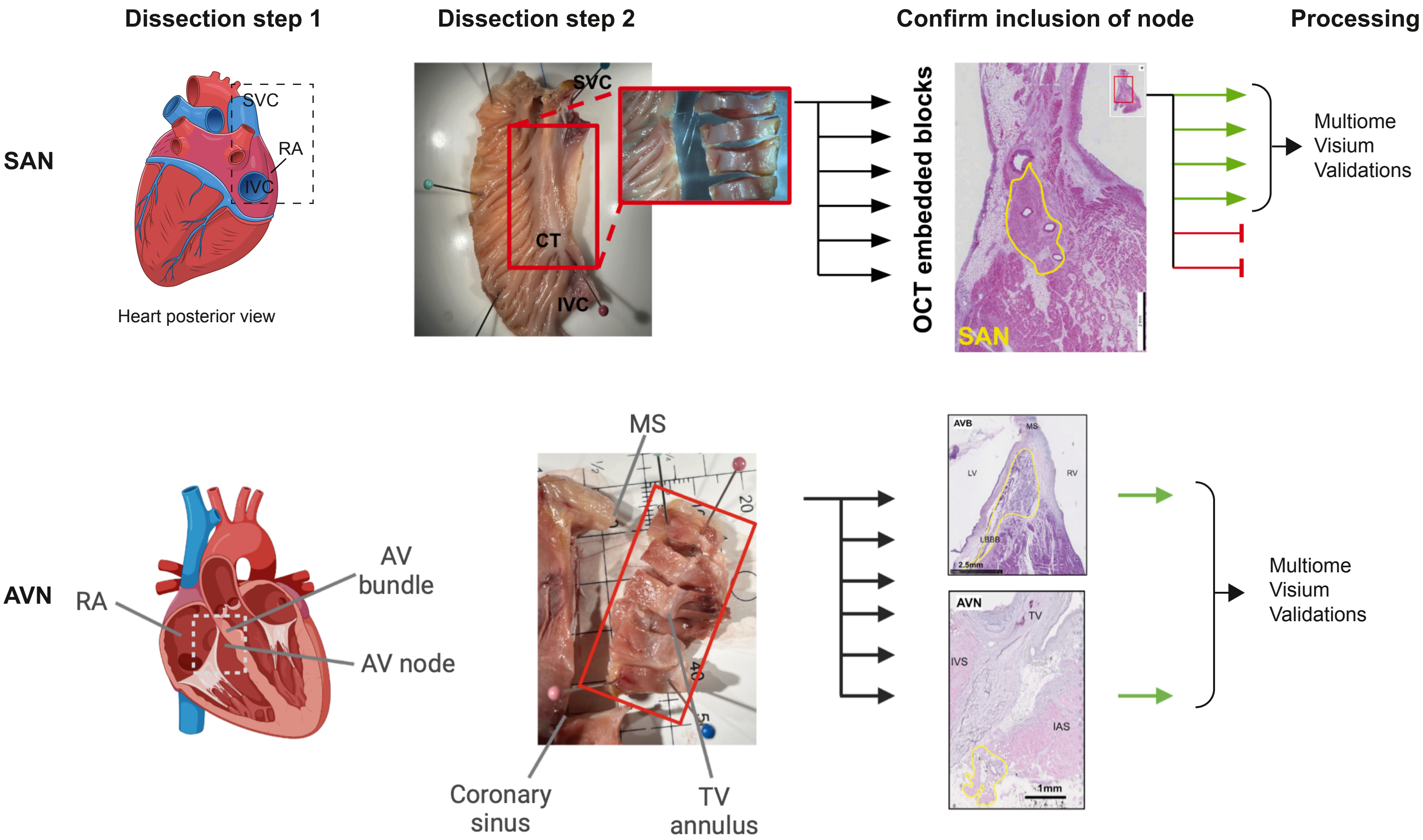

Dissection step 1
Dissection step 2
Confirm inclusion of node
Processing
SAN
SVC
RA
IVC
Heart posterior view
SVC
CT
IVC
OCT embedded blocks
SAN
Multiome
Visium
Validations
AVN
RA
AV
bundle
AV node
MS
Coronary
sinus
TV
annulus
AVB
MS
LV
RV
LBBB
2.5mm
AVN
TV
IVS
IAS
1mm
Multiome
Visium
Validations

Supp Fig 2

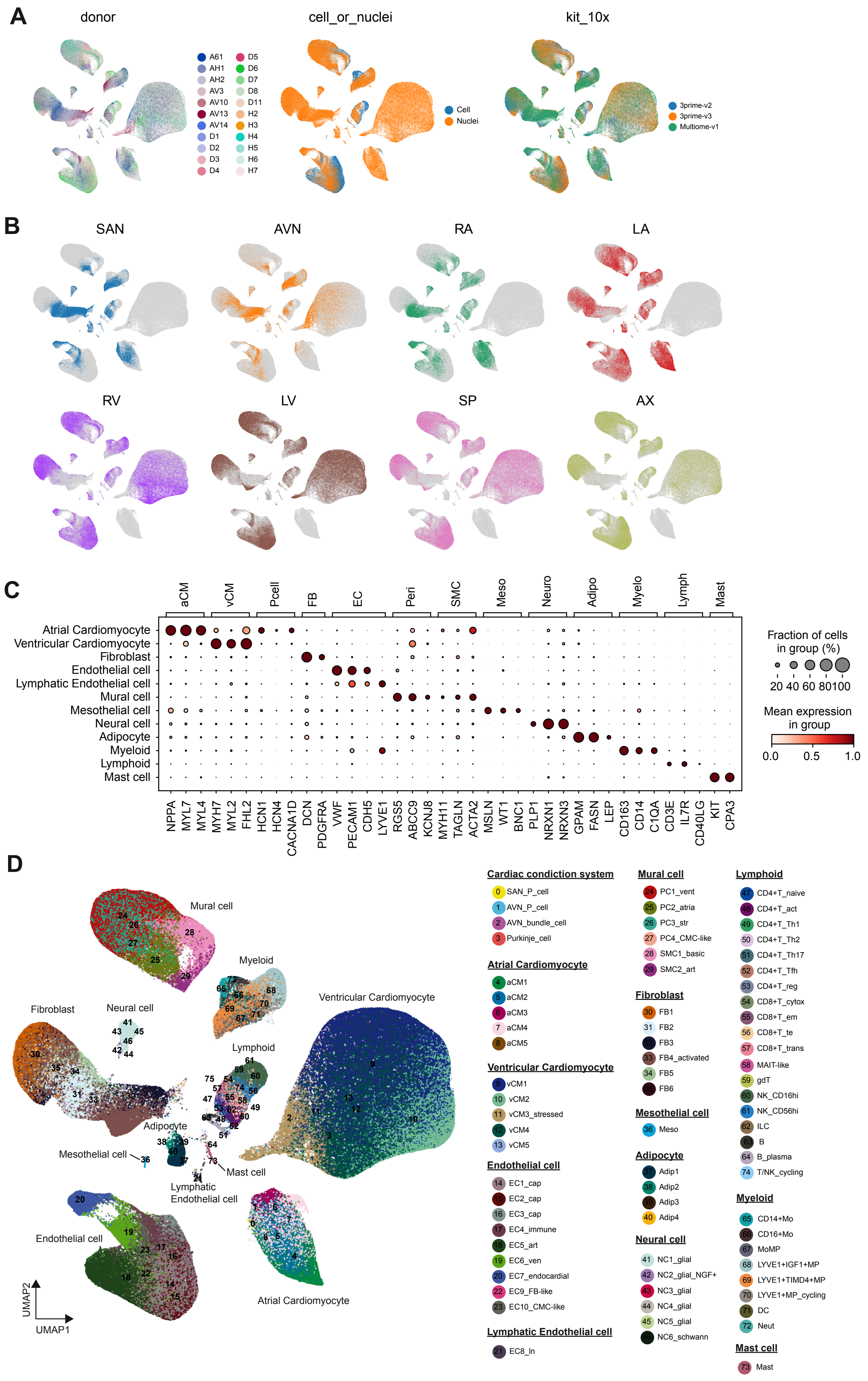
A
donor
A61
AH1
AH2
AV3
AV10
AV13
AV14
D1
D2
D3
D4
D5
D6
D7
D8
D11
H2
H3
H4
H5
H6
H7
cell_or_nuclei
Cell
Nuclei
kit_10x
3prime-v2
3prime-v3
Multiome-v1
B
SAN
AVN
RA
LA
RV
LV
SP
AX
C
aCM
vCM
Pcell
FB
EC
Peri
SMC
Meso
Neuro
Adipo
Myelo
Lymph
Mast
Atrial Cardiomyocyte
Ventricular Cardiomyocyte
Fibroblast
Endothelial cell
Lymphatic Endothelial cell
Mural cell
Mesothelial cell
Neural cell
Adipocyte
Myeloid
Lymphoid
Mast cell
NPPA
MYL7
MYL4
MYH7
MYL2
FHL2
HCN1
HCN4
CACNA1D
DCN
PDGFRA
VWF
PECAM1
CDH5
LYVE1
RGS5
ABCC9
KCNJ8
MYH11
TAGLN
ACTA2
MSLN
WT1
BNC1
PLP1
NRXN1
NRXN3
GPAM
FASN
LEP
CD163
CD14
C1QA
CD3E
IL7R
CD40LG
KIT
CPA3
Fraction of cells
in group (%)
20 40 60 80100
Mean expression
in group
0.0
0.5
1.0
D
Mural cell
Myeloid
Fibroblast
Neural cell
Ventricular Cardiomyocyte
Lymphoid
Adipocyte
Mesothelial cell
Mast cell
Lymphatic
Endothelial cell
Endothelial cell
Atrial Cardiomyocyte
UMAP2
UMAP1
Cardiac condiction system
0 SAN_P_cell
1 AVN_P_cell
2 AVN_bundle_cell
3 Purkinje_cell
Atrial Cardiomyocyte
4 aCM1
5 aCM2
6 aCM3
7 aCM4
8 aCM5
Ventricular Cardiomyocyte
9 vCM1
10 vCM2
11 vCM3_stressed
12 vCM4
13 vCM5
Endothelial cell
14 EC1_cap
15 EC2_cap
16 EC3_cap
17 EC4_immune
18 EC5_art
19 EC6_ven
20 EC7_endocardial
22 EC9_FB-like
23 EC10_CMC-like
Lymphatic Endothelial cell
21 EC8_ln
Mural cell
24 PC1_vent
25 PC2_atria
26 PC3_str
27 PC4_CMC-like
28 SMC1_basic
29 SMC2_art
Fibroblast
30 FB1
31 FB2
32 FB3
33 FB4_activated
34 FB5
35 FB6
Mesothelial cell
36 Meso
Adipocyte
37 Adip1
38 Adip2
39 Adip3
40 Adip4
Neural cell
41 NC1_glial
42 NC2_glial_NGF+
43 NC3_glial
44 NC4_glial
45 NC5_glial
46 NC6_schwann
Lymphoid
47 CD4+T_naive
48 CD4+T_act
49 CD4+T_Th1
50 CD4+T_Th2
51 CD4+T_Th17
52 CD4+T_Tfh
53 CD4+T_reg
54 CD8+T_cytox
55 CD8+T_em
56 CD8+T_te
57 CD8+T_trans
58 MAIT-like
59 gdT
60 NK_CD16hi
61 NK_CD56hi
62 ILC
63 B
64 B_plasma
74 T/NK_cycling
Myeloid
65 CD14+Mo
66 CD16+Mo
67 MoMP
68 LYVE1+IGF1+MP
69 LYVE1+TIMD4+MP
70 LYVE1+MP_cycling
71 DC
72 Neut
Mast cell
73 Mast

### Supp Fig 3

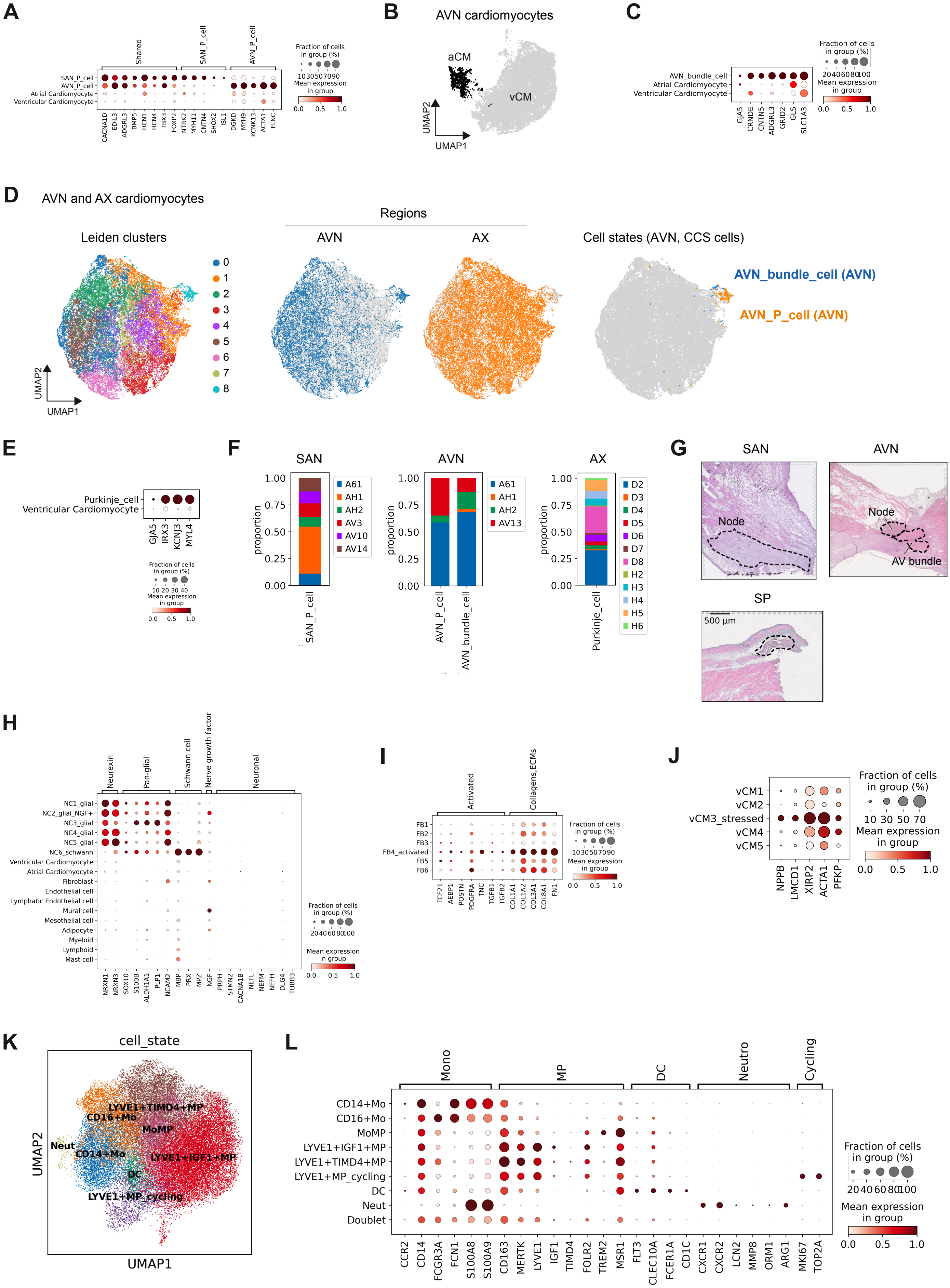

A
B
AVN cardiomyocytes
aCM
vCM
UMAP1
UMAP2
C
D
AVN and AX cardiomyocytes
Regions
Leiden clusters
AVN
AX
Cell states (AVN, CCS cells)
AVN_bundle_cell (AVN)
AVN_P_cell (AVN)
E
F
SAN
AVN
AX
G
SAN
AVN
Node
AV bundle
SP
500 µm
H
I
J
K
cell_state
L

### Supp Fig 4

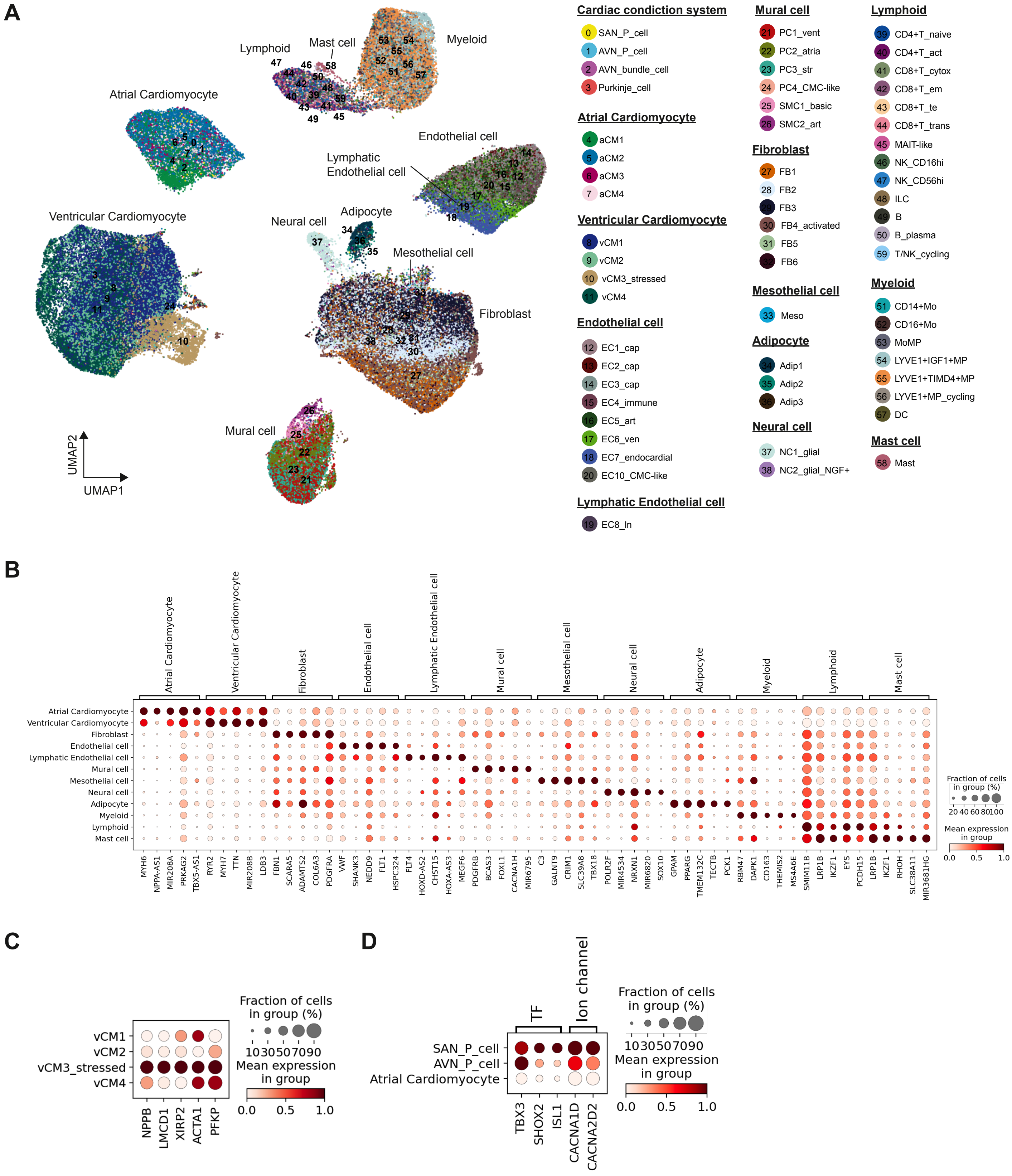
A
Lymphoid
Mast cell
Myeloid
Atrial Cardiomyocyte
Endothelial cell
Lymphatic Endothelial cell
Ventricular Cardiomyocyte
Neural cell
Adipocyte
Mesothelial cell
Fibroblast
Mural cell
UMAP2
UMAP1
Cardiac condiction system
0 SAN_P_cell
1 AVN_P_cell
2 AVN_bundle_cell
3 Purkinje_cell
Atrial Cardiomyocyte
4 aCM1
5 aCM2
6 aCM3
7 aCM4
Ventricular Cardiomyocyte
8 vCM1
9 vCM2
10 vCM3_stressed
11 vCM4
Endothelial cell
12 EC1_cap
13 EC2_cap
14 EC3_cap
15 EC4_immune
16 EC5_art
17 EC6_ven
18 EC7_endocardial
20 EC10_CMC-like
Lymphatic Endothelial cell
19 EC8_ln
Mural cell
21 PC1_vent
22 PC2_atria
23 PC3_str
24 PC4_CMC-like
25 SMC1_basic
26 SMC2_art
Fibroblast
27 FB1
28 FB2
29 FB3
30 FB4_activated
31 FB5
32 FB6
Mesothelial cell
33 Meso
Adipocyte
34 Adip1
35 Adip2
36 Adip3
Neural cell
37 NC1_glial
38 NC2_glial_NGF+
Lymphoid
39 CD4+T_naive
40 CD4+T_act
41 CD8+T_cytox
42 CD8+T_em
43 CD8+T_te
44 CD8+T_trans
45 MAIT-like
46 NK_CD16hi
47 NK_CD56hi
48 ILC
49 B
50 B_plasma
59 T/NK_cycling
Myeloid
51 CD14+Mo
52 CD16+Mo
53 MoMP
54 LYVE1+IGF1+MP
55 LYVE1+TIMD4+MP
56 LYVE1+MP_cycling
57 DC
Mast cell
58 Mast
B
Atrial Cardiomyocyte
Ventricular Cardiomyocyte
Fibroblast
Endothelial cell
Lymphatic Endothelial cell
Mural cell
Mesothelial cell
Neural cell
Adipocyte
Myeloid
Lymphoid
Mast cell
MYH6 NPPA-AS1 MIR208A PRKAG2 TBX5-AS1 RYR2 MYH7 TTN MIR208B LDB3 FBN1 SCARA5 ADAMTS2 COL6A3 PDGFRA VWF SHANK3 NEDD9 FLT1 HSPC324 FLT4 HOXD-AS2 CHST15 HOXA-AS3 MEGF6 PDGFRB BCAS3 FOXL1 CACNA1H MIR6795 C3 GALNT9 CRIM1 SLC39A8 TBX18 POLR2F MIR4534 NRXN1 MIR6820 SOX10 GPAM PPARG TMEM132C TECTB PCK1 RBM47 DAPK1 CD163 THEMIS2 MS4A6E SMIM11B LRP1B IKZF1 EYS PCDH15 LRP1B IKZF1 RHOH SLC38A11 MIR3681HG
Fraction of cells in group (%)
20 40 60 80 100
Mean expression in group
0.0 0.5 1.0
C
vCM1
vCM2
vCM3_stressed
vCM4
NPPB LMCD1 XIRP2 ACTA1 PFKP
Fraction of cells in group (%)
10 30 50 70 90
Mean expression in group
0.0 0.5 1.0
D
TF
Ion channel
SAN_P_cell
AVN_P_cell
Atrial Cardiomyocyte
TBX3 SHOX2 ISL1 CACNA1D CACNA2D2
Fraction of cells in group (%)
10 30 50 70 90
Mean expression in group
0.0 0.5 1.0

Supp Fig 5

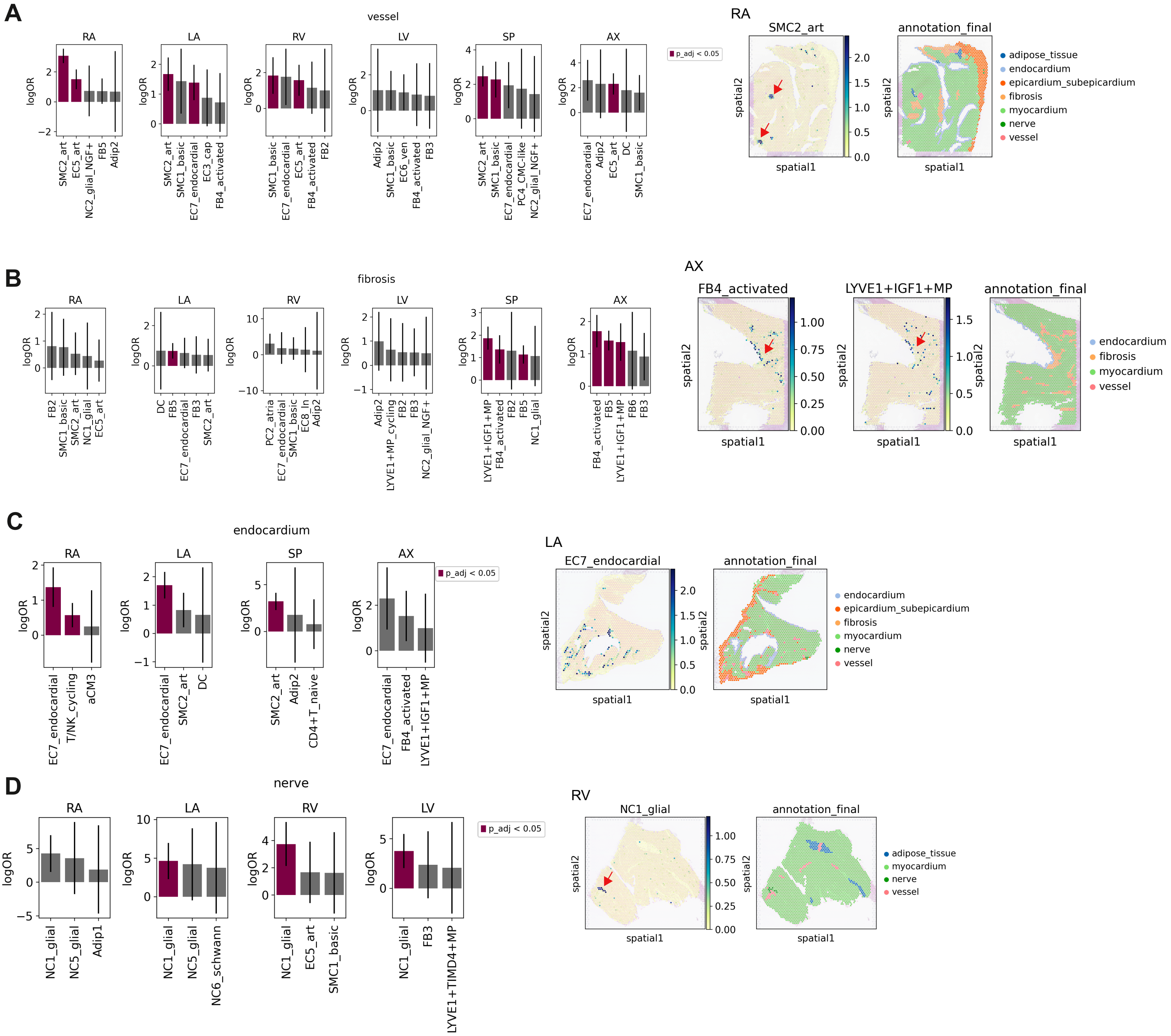
A
vessel
RA
LA
RV
LV
SP
AX
logOR
SMC2_art
EC5_art
NC2_glial_NGF+
FB5
Adip2
SMC1_basic
EC7_endocardial
EC3_cap
FB4_activated
FB2
EC6_ven
FB3
PC4_CMC-like
DC
p_adj < 0.05
RA
SMC2_art
annotation_final
spatial1
spatial2
adipose_tissue
endocardium
epicardium_subepicardium
fibrosis
myocardium
nerve
vessel
B
fibrosis
RA
LA
RV
LV
SP
AX
logOR
FB2
SMC1_basic
SMC2_art
NC1_glial
EC5_art
DC
FB5
EC7_endocardial
FB3
PC2_atria
EC8_ln
Adip2
LYVE1+MP_cycling
NC2_glial_NGF+
LYVE1+IGF1+MP
FB4_activated
FB6
AX
FB4_activated
LYVE1+IGF1+MP
annotation_final
spatial1
spatial2
endocardium
fibrosis
myocardium
vessel
C
endocardium
RA
LA
SP
AX
logOR
EC7_endocardial
T/NK_cycling
aCM3
SMC2_art
DC
Adip2
CD4+T_naive
FB4_activated
LYVE1+IGF1+MP
p_adj < 0.05
LA
EC7_endocardial
annotation_final
spatial1
spatial2
endocardium
epicardium_subepicardium
fibrosis
myocardium
nerve
vessel
D
nerve
RA
LA
RV
LV
logOR
NC1_glial
NC5_glial
Adip1
NC6_schwann
EC5_art
SMC1_basic
FB3
LYVE1+TIMD4+MP
p_adj < 0.05
RV
NC1_glial
annotation_final
spatial1
spatial2
adipose_tissue
myocardium
nerve
vessel

Supp Fig 6

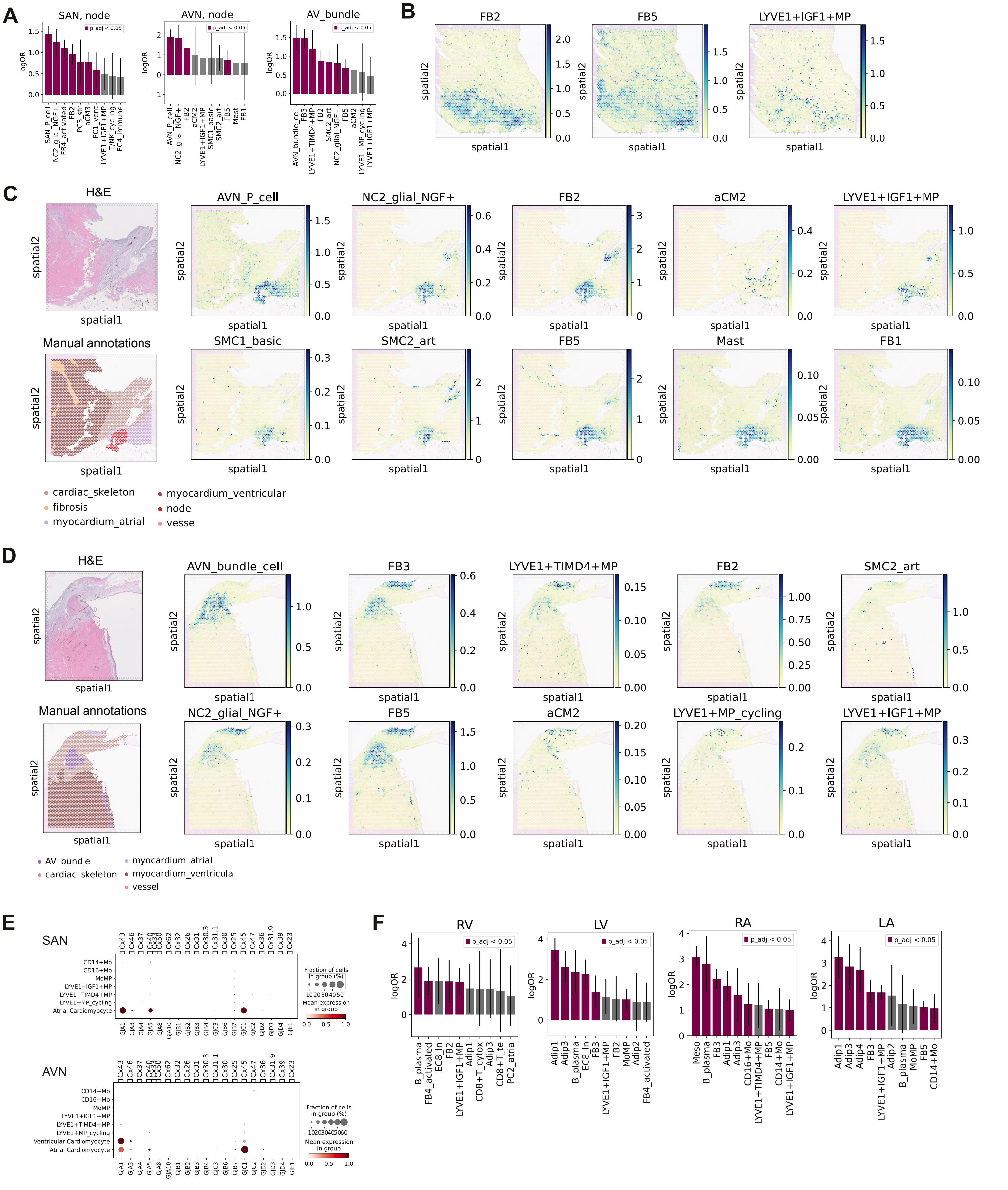

A
SAN, node
AVN, node
AV_bundle
p_adj < 0.05
logOR
SAN_P_cell
NC2_glial_NGF+
FB4_activated
FB2
PC3_str
aCM3
PC1_vent
LYVE1+IGF1+MP
T/NK_cycling
EC4_immune
AVN_P_cell
aCM2
SMC1_basic
SMC2_art
FB5
Mast
FB1
AVN_bundle_cell
FB3
LYVE1+TIMD4+MP
LYVE1+MP_cycling
B
FB2
FB5
LYVE1+IGF1+MP
spatial1
spatial2
C
H&E
Manual annotations
cardiac_skeleton
fibrosis
myocardium_atrial
myocardium_ventricular
node
vessel
D
AVN_bundle_cell
AV_bundle
myocardium_ventricula
E
SAN
AVN
CD14+Mo
CD16+Mo
MoMP
LYVE1+IGF1+MP
LYVE1+TIMD4+MP
LYVE1+MP_cycling
Atrial Cardiomyocyte
Ventricular Cardiomyocyte
Fraction of cells in group (%)
Mean expression in group
F
RV
LV
RA
LA
B_plasma
EC8_ln
Adip1
Adip2
Adip3
Adip4
CD8+T_cytox
CD8+T_te
PC2_atria
Meso
CD16+Mo
CD14+Mo

Supp. Fig 7

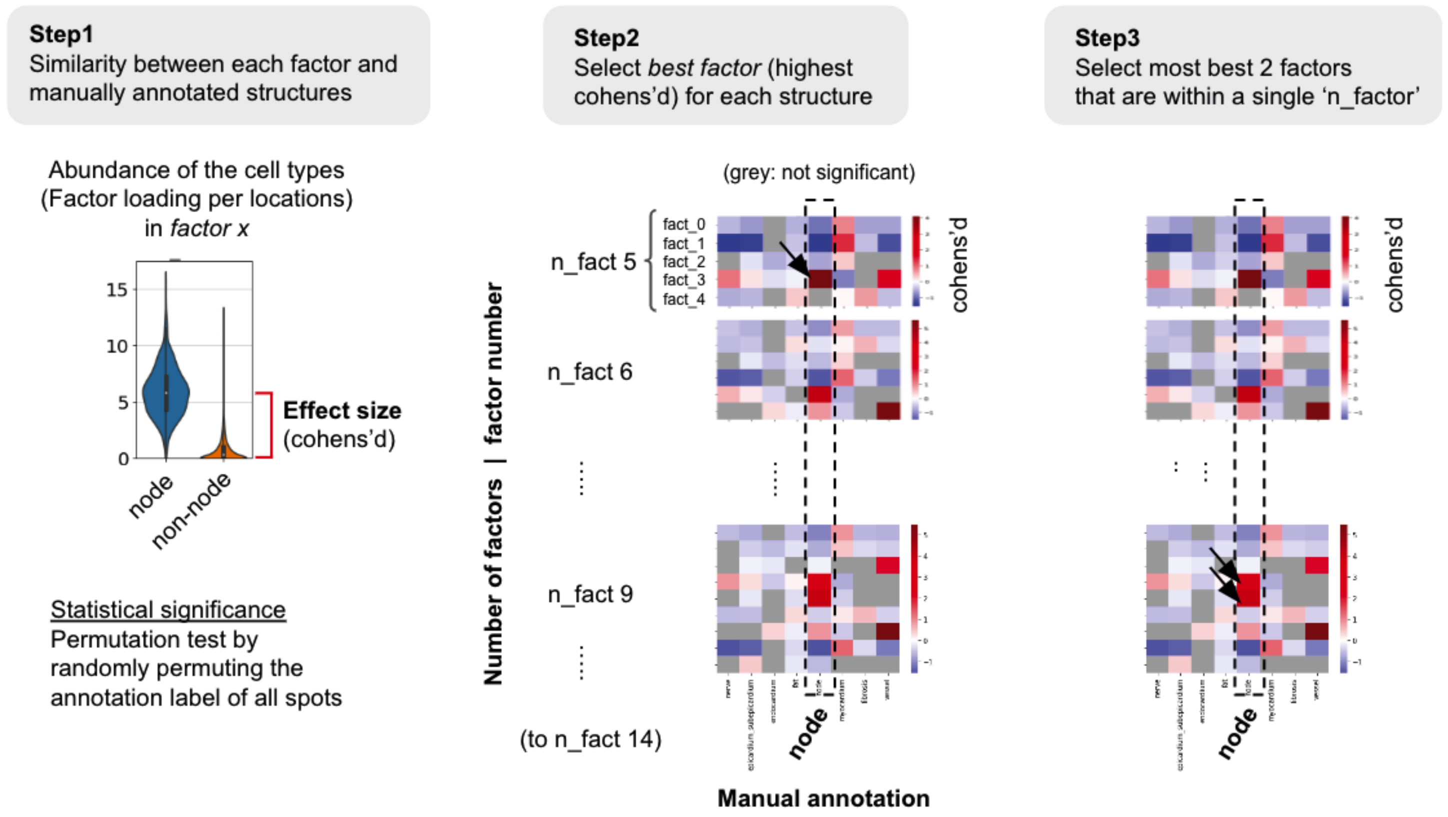
Step1
Similarity between each factor and manually annotated structures
Abundance of the cell types
(Factor loading per locations)
in factor x
15
10
5
0
node
non-node
Effect size
(cohens'd)
Statistical significance
Permutation test by randomly permuting the annotation label of all spots
Step2
Select best factor (highest cohens'd) for each structure
(grey: not significant)
fact_0
fact_1
fact_2
fact_3
fact_4
n_fact 5
n_fact 6
n_fact 9
(to n_fact 14)
Number of factors | factor number
cohens'd
node
Manual annotation
Step3
Select most best 2 factors that are within a single 'n_factor'
cohens'd
node

Supp. Fig 8

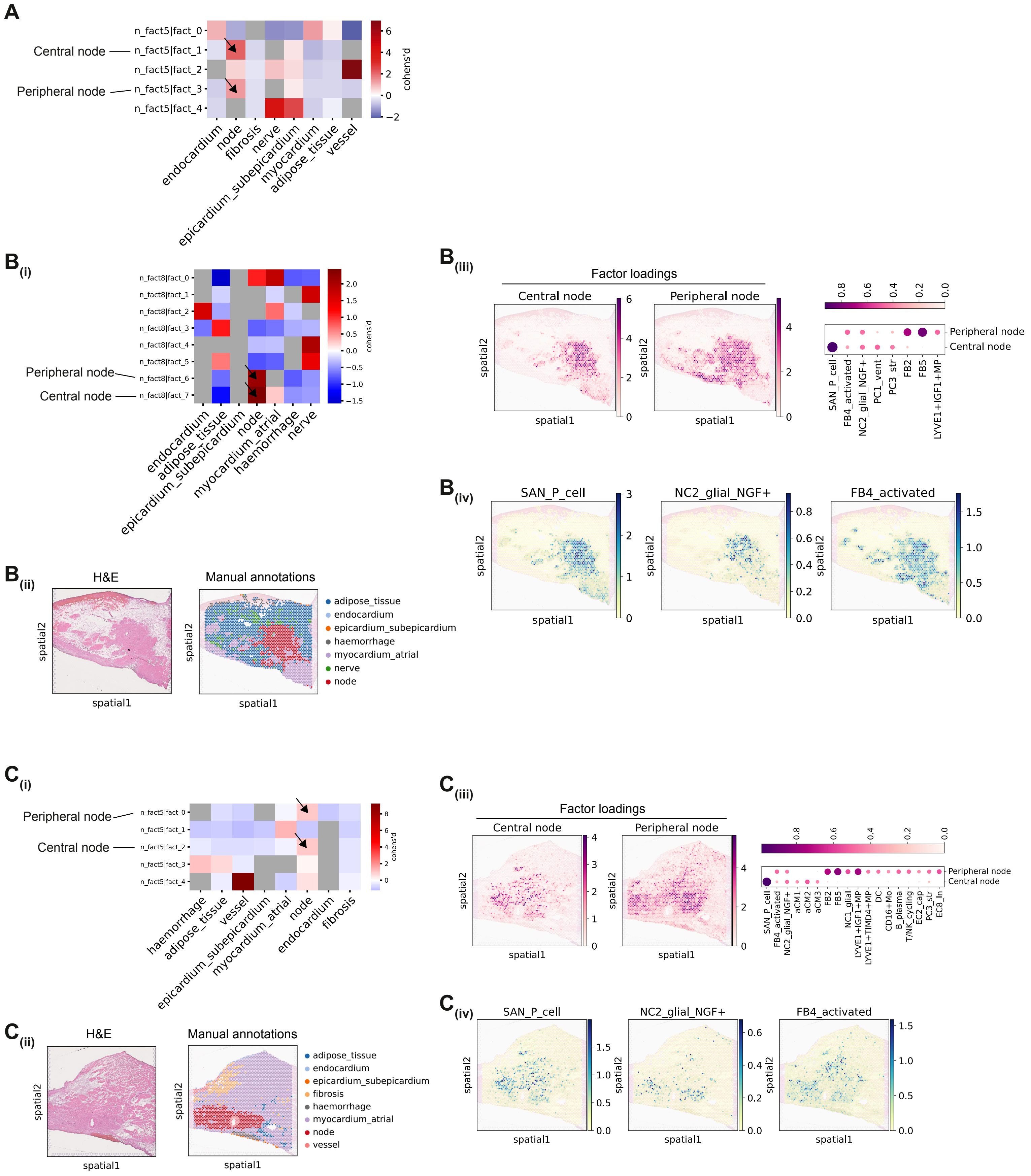
A
Central node
Peripheral node
n_fact5|fact_0
n_fact5|fact_1
n_fact5|fact_2
n_fact5|fact_3
n_fact5|fact_4
cohens'd
endocardium
node
fibrosis
nerve
epicardium_subepicardium
myocardium
adipose_tissue
vessel
B(i)
Peripheral node
Central node
n_fact8|fact_0
n_fact8|fact_1
n_fact8|fact_2
n_fact8|fact_3
n_fact8|fact_4
n_fact8|fact_5
n_fact8|fact_6
n_fact8|fact_7
cohens'd
endocardium
adipose_tissue
epicardium_subepicardium
node
myocardium_atrial
haemorrhage
nerve
B(ii)
H&E
Manual annotations
spatial1
spatial2
adipose_tissue
endocardium
epicardium_subepicardium
haemorrhage
myocardium_atrial
nerve
node
B(iii)
Factor loadings
Central node
Peripheral node
spatial1
spatial2
Peripheral node
Central node
SAN_P_cell
FB4_activated
NC2_glial_NGF+
PC1_vent
PC3_str
FB2
FB5
LYVE1+IGF1+MP
B(iv)
SAN_P_cell
NC2_glial_NGF+
FB4_activated
spatial1
spatial2
C(i)
Peripheral node
Central node
n_fact5|fact_0
n_fact5|fact_1
n_fact5|fact_2
n_fact5|fact_3
n_fact5|fact_4
cohens'd
haemorrhage
adipose_tissue
vessel
epicardium_subepicardium
myocardium_atrial
node
endocardium
fibrosis
C(ii)
H&E
Manual annotations
spatial1
spatial2
adipose_tissue
endocardium
epicardium_subepicardium
fibrosis
haemorrhage
myocardium_atrial
node
vessel
C(iii)
Factor loadings
Central node
Peripheral node
spatial1
spatial2
Peripheral node
Central node
SAN_P_cell
FB4_activated
NC2_glial_NGF+
aCM1
aCM2
aCM3
FB2
FB5
NC1_glial
LYVE1+IGF1+MP
LYVE1+TIMD4+MP
DC
CD16+Mo
B_plasma
T/NK_cycling
EC2_cap
PC3_str
EC8_ln
C(iv)
SAN_P_cell
NC2_glial_NGF+
FB4_activated
spatial1
spatial2

Supp. Fig 9

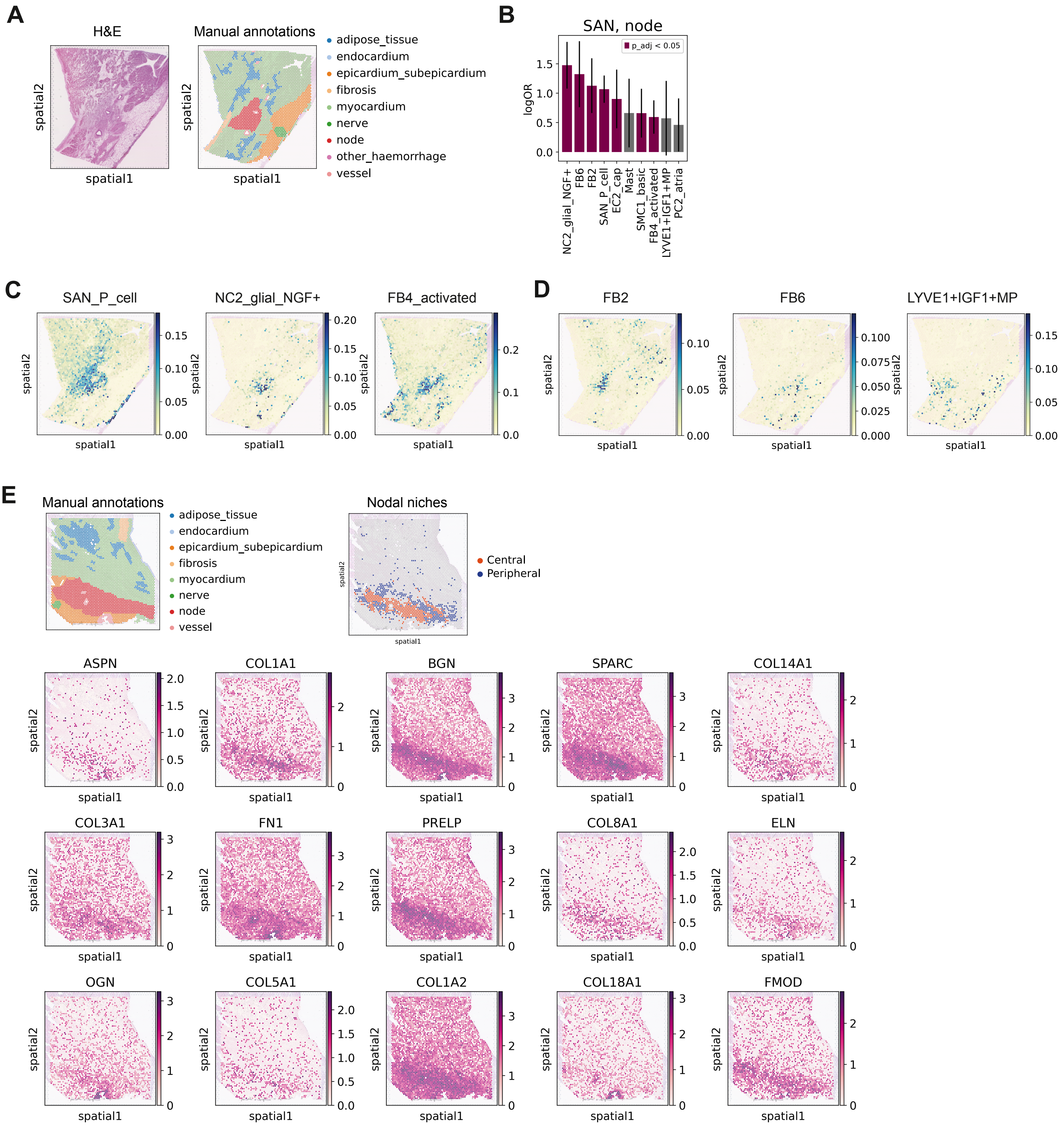
A
H&E
Manual annotations
spatial2
spatial1
adipose_tissue
endocardium
epicardium_subepicardium
fibrosis
myocardium
nerve
node
other_haemorrhage
vessel
B
SAN, node
p_adj < 0.05
logOR
NC2_glial_NGF+
FB6
FB2
SAN_P_cell
EC2_cap
Mast
SMC1_basic
FB4_activated
LYVE1+IGF1+MP
PC2_atria
C
SAN_P_cell
NC2_glial_NGF+
FB4_activated
D
FB2
FB6
LYVE1+IGF1+MP
E
Manual annotations
Nodal niches
Central
Peripheral
ASPN
COL1A1
BGN
SPARC
COL14A1
COL3A1
FN1
PRELP
COL8A1
ELN
OGN
COL5A1
COL1A2
COL18A1
FMOD

Supp Fig 10

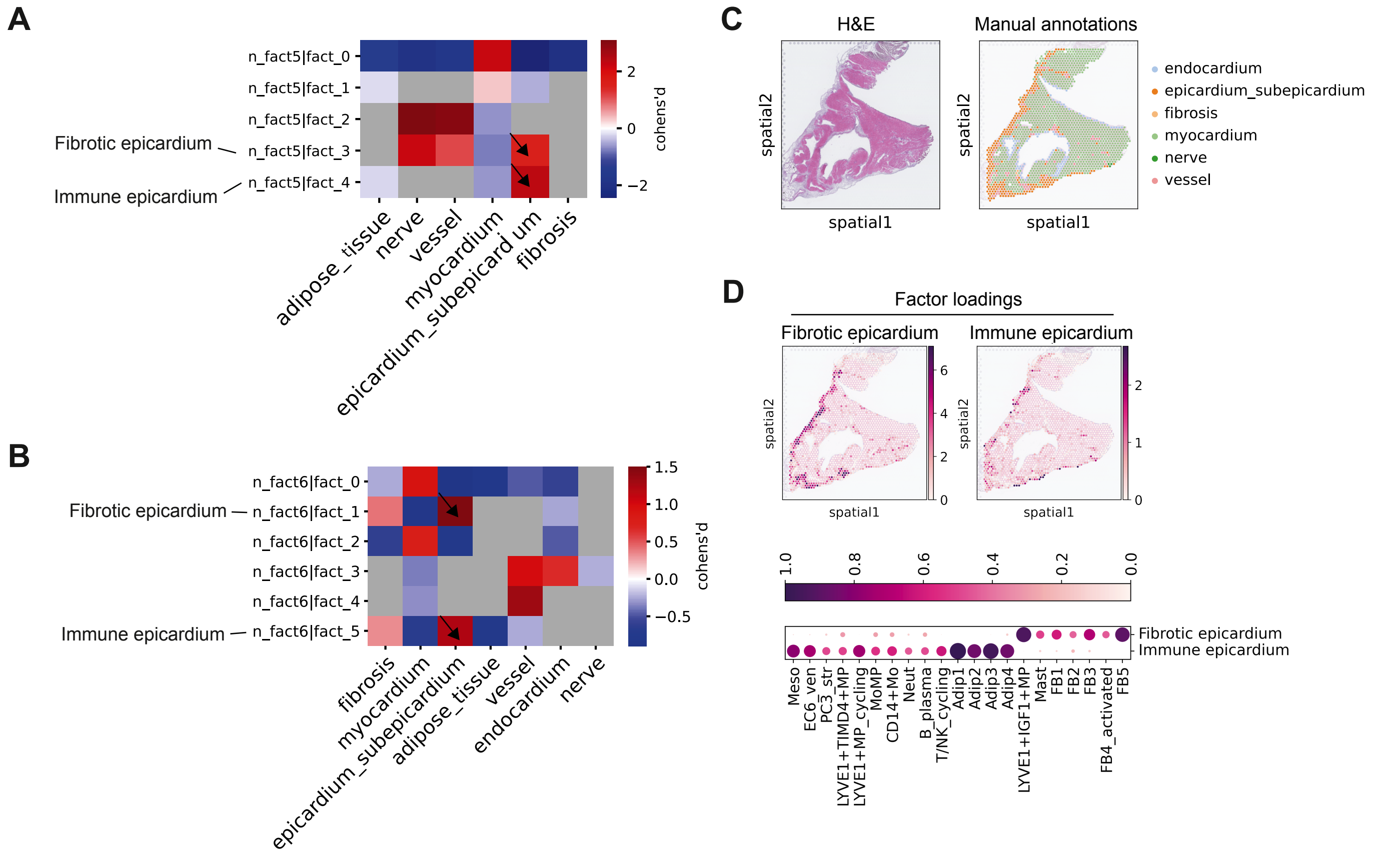
A
n_fact5|fact_0
n_fact5|fact_1
n_fact5|fact_2
n_fact5|fact_3
n_fact5|fact_4
Fibrotic epicardium
Immune epicardium
adipose_tissue
nerve
vessel
myocardium
epicardium_subepicardium
fibrosis
cohens'd
B
n_fact6|fact_0
n_fact6|fact_1
n_fact6|fact_2
n_fact6|fact_3
n_fact6|fact_4
n_fact6|fact_5
Fibrotic epicardium
Immune epicardium
fibrosis
myocardium
epicardium_subepicardium
adipose_tissue
vessel
endocardium
nerve
cohens'd
C
H&E
Manual annotations
spatial1
spatial2
endocardium
epicardium_subepicardium
fibrosis
myocardium
nerve
vessel
D
Factor loadings
Fibrotic epicardium
Immune epicardium
spatial1
spatial2
Meso
EC6_ven
PC3_str
LYVE1+TIMD4+MP
LYVE1+MP_cycling
MoMP
CD14+Mo
Neut
B_plasma
T/NK_cycling
Adip1
Adip2
Adip3
Adip4
LYVE1+IGF1+MP
Mast
FB1
FB2
FB3
FB4_activated
FB5

Supp Fig 11

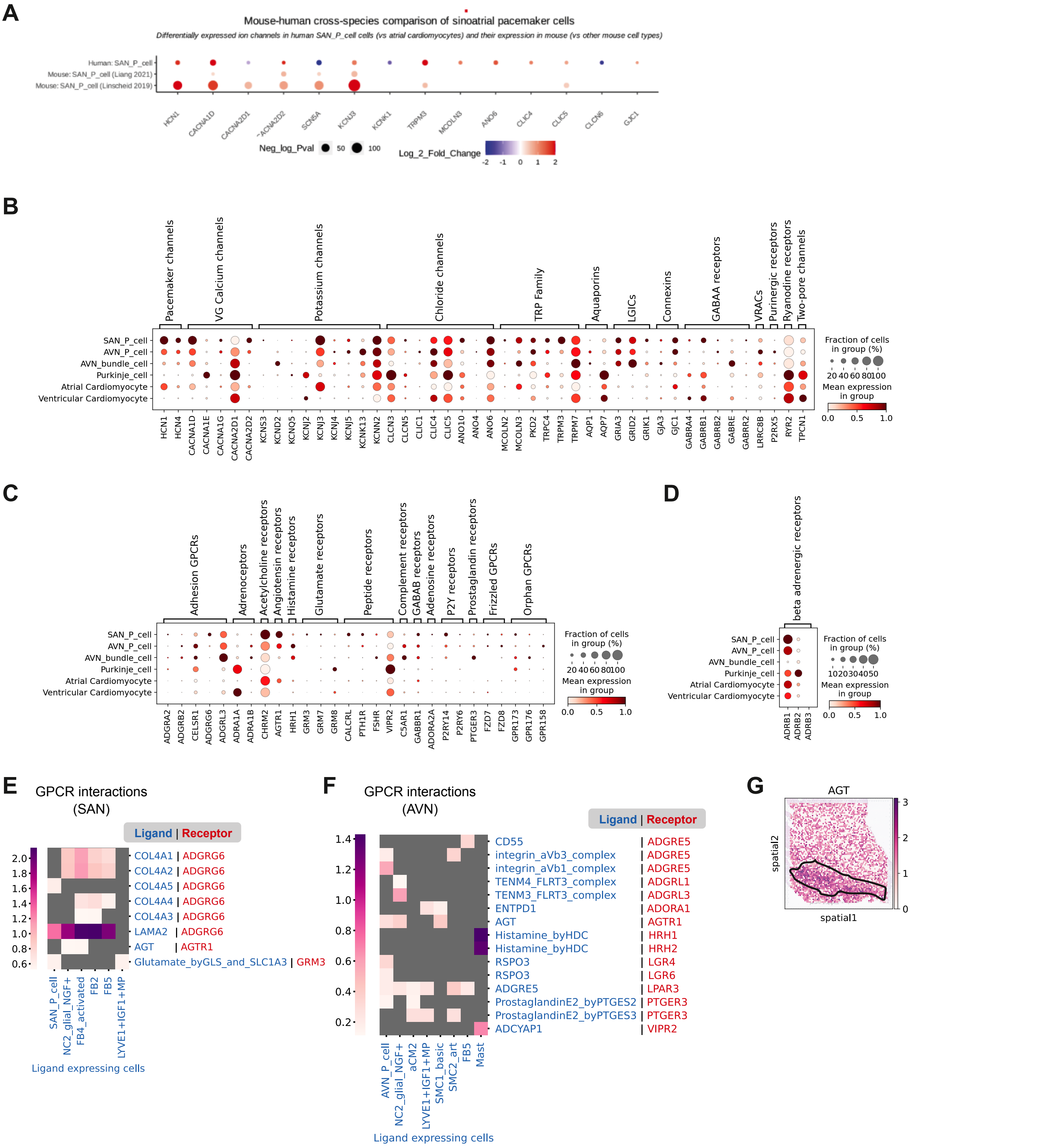
A
Mouse-human cross-species comparison of sinoatrial pacemaker cells
Differentially expressed ion channels in human SAN_P_cell cells (vs atrial cardiomyocytes) and their expression in mouse (vs other mouse cell types)
Human: SAN_P_cell
Mouse: SAN_P_cell (Liang 2021)
Mouse: SAN_P_cell (Linscheid 2019)
HCN1 CACNA1D CACNA2D1 CACNA2D2 SCN5A KCNJ3 KCNK1 TRPM3 MCOLN3 ANO6 CLIC4 CLIC5 CLCN6 GJC1
Neg_log_Pval 50 100
Log_2_Fold_Change -2 -1 0 1 2
B
Pacemaker channels
VG Calcium channels
Potassium channels
Chloride channels
TRP Family
Aquaporins
LGICs
Connexins
GABAA receptors
VRACs
Purinergic receptors
Ryanodine receptors
Two-pore channels
SAN_P_cell
AVN_P_cell
AVN_bundle_cell
Purkinje_cell
Atrial Cardiomyocyte
Ventricular Cardiomyocyte
HCN1 HCN4 CACNA1D CACNA1E CACNA1G CACNA2D1 CACNA2D2 KCNS3 KCND2 KCNQ5 KCNJ2 KCNJ3 KCNJ4 KCNJ5 KCNK13 KCNN2 CLCN3 CLCN5 CLIC1 CLIC4 CLIC5 ANO10 ANO4 ANO6 MCOLN2 MCOLN3 PKD2 TRPC4 TRPM3 TRPM7 AQP1 AQP7 GRIA3 GRID2 GRIK1 GJA3 GJC1 GABRA4 GABRB1 GABRB2 GABRE GABRR2 LRRC8B P2RX5 RYR2 TPCN1
Fraction of cells in group (%)
20 40 60 80 100
Mean expression in group
0.0 0.5 1.0
C
Adhesion GPCRs
Adrenoceptors
Acetylcholine receptors
Angiotensin receptors
Histamine receptors
Glutamate receptors
Peptide receptors
Complement receptors
GABAB receptors
Adenosine receptors
P2Y receptors
Prostaglandin receptors
Frizzled GPCRs
Orphan GPCRs
SAN_P_cell
AVN_P_cell
AVN_bundle_cell
Purkinje_cell
Atrial Cardiomyocyte
Ventricular Cardiomyocyte
ADGRA2 ADGRB2 CELSR1 ADGRG6 ADGRL3 ADRA1A ADRA1B CHRM2 AGTR1 HRH1 GRM3 GRM7 GRM8 CALCRL PTH1R FSHR VIPR2 C5AR1 GABBR1 ADORA2A P2RY14 P2RY6 PTGER3 FZD7 FZD8 GPR173 GPR176 GPR158
Fraction of cells in group (%)
20 40 60 80 100
Mean expression in group
0.0 0.5 1.0
D
beta adrenergic receptors
SAN_P_cell
AVN_P_cell
AVN_bundle_cell
Purkinje_cell
Atrial Cardiomyocyte
Ventricular Cardiomyocyte
ADRB1 ADRB2 ADRB3
Fraction of cells in group (%)
10 20 30 40 50
Mean expression in group
0.0 0.5 1.0
E
GPCR interactions (SAN)
Ligand | Receptor
COL4A1 | ADGRG6
COL4A2 | ADGRG6
COL4A5 | ADGRG6
COL4A4 | ADGRG6
COL4A3 | ADGRG6
LAMA2 | ADGRG6
AGT | AGTR1
Glutamate_byGLS_and_SLC1A3 | GRM3
2.0 1.8 1.6 1.4 1.2 1.0 0.8 0.6
SAN_P_cell
NC2_glial_NGF+
FB4_activated
FB2
FB5
LYVE1+IGF1+MP
Ligand expressing cells
F
GPCR interactions (AVN)
Ligand | Receptor
CD55 | ADGRE5
integrin_aVb3_complex | ADGRE5
integrin_aVb1_complex | ADGRE5
TENM4_FLRT3_complex | ADGRL1
TENM3_FLRT3_complex | ADGRL3
ENTPD1 | ADORA1
AGT | AGTR1
Histamine_byHDC | HRH1
Histamine_byHDC | HRH2
RSPO3 | LGR4
RSPO3 | LGR6
ADGRE5 | LPAR3
ProstaglandinE2_byPTGES2 | PTGER3
ProstaglandinE2_byPTGES3 | PTGER3
ADCYAP1 | VIPR2
1.4 1.2 1.0 0.8 0.6 0.4 0.2
AVN_P_cell
NC2_glial_NGF+
aCM2
LYVE1+IGF1+MP
SMC1_basic
SMC2_art
FB5
Mast
Ligand expressing cells
G
AGT
spatial2
spatial1
3 2 1 0

Supp Fig 12

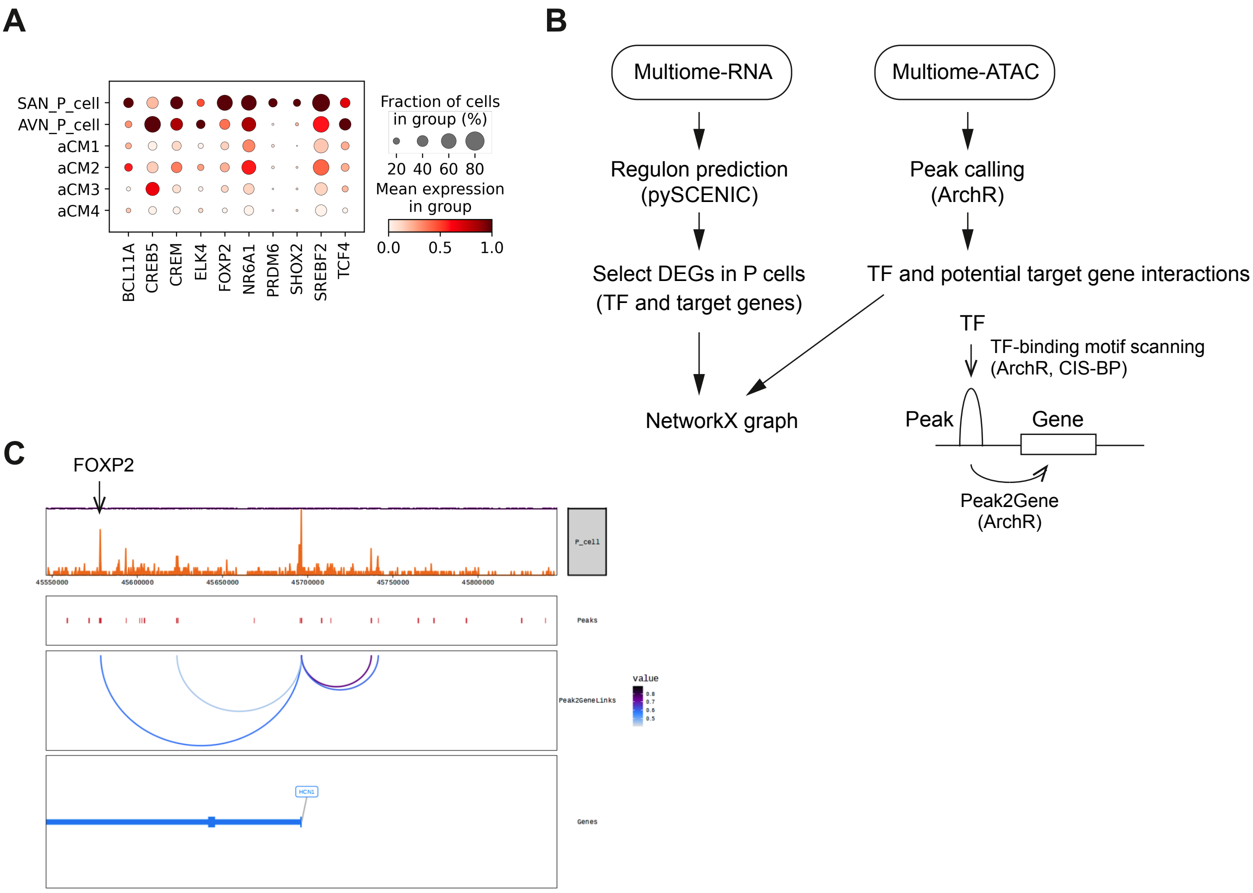
A
SAN_P_cell
AVN_P_cell
aCM1
aCM2
aCM3
aCM4
BCL11A
CREB5
CREM
ELK4
FOXP2
NR6A1
PRDM6
SHOX2
SREBF2
TCF4
Fraction of cells in group (%)
20 40 60 80
Mean expression in group
0.0 0.5 1.0
B
Multiome-RNA
Multiome-ATAC
Regulon prediction (pySCENIC)
Peak calling (ArchR)
Select DEGs in P cells (TF and target genes)
TF and potential target gene interactions
TF
TF-binding motif scanning (ArchR, CIS-BP)
NetworkX graph
Peak
Gene
Peak2Gene (ArchR)
C
FOXP2
P_cell
45550000
45600000
45650000
45700000
45750000
45800000
Peaks
Peak2GeneLinks
value
Genes

Supp Fig 13

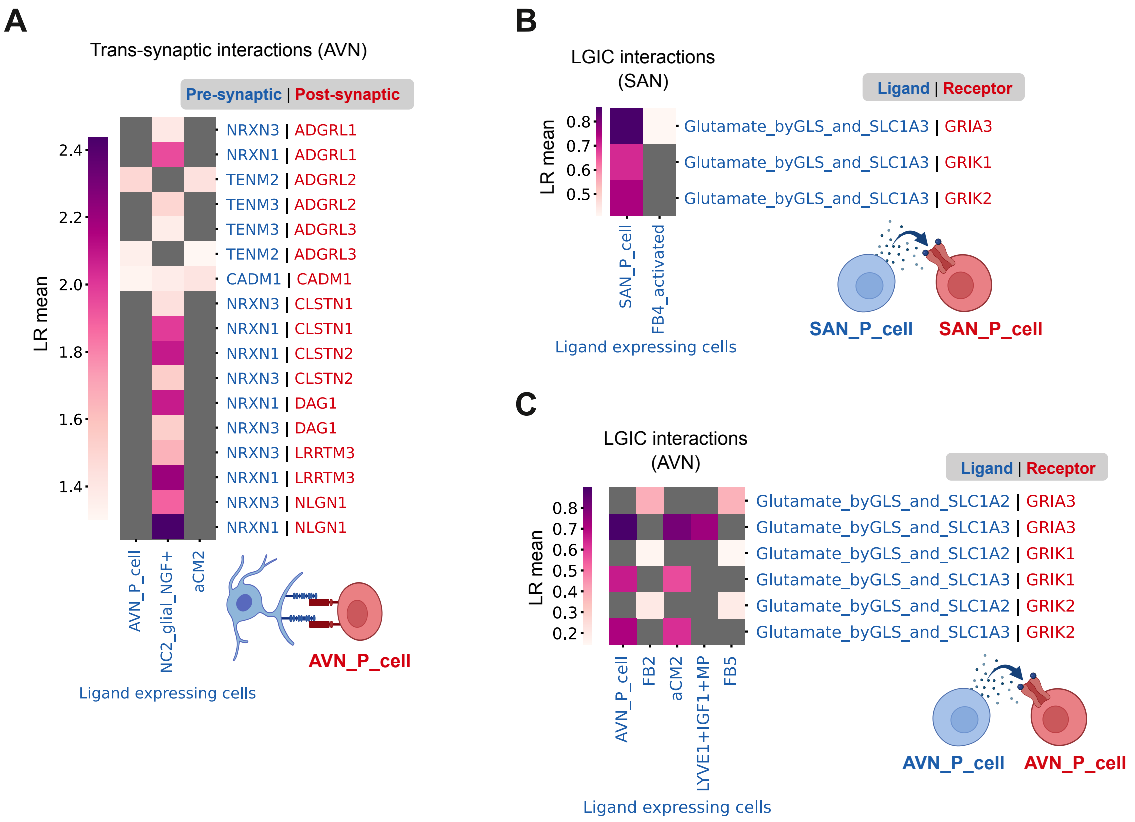
A
Trans-synaptic interactions (AVN)
Pre-synaptic | Post-synaptic
LR mean
2.4
2.2
2.0
1.8
1.6
1.4
NRXN3 | ADGRL1
NRXN1 | ADGRL1
TENM2 | ADGRL2
TENM3 | ADGRL2
TENM3 | ADGRL3
TENM2 | ADGRL3
CADM1 | CADM1
NRXN3 | CLSTN1
NRXN1 | CLSTN1
NRXN1 | CLSTN2
NRXN3 | CLSTN2
NRXN1 | DAG1
NRXN3 | DAG1
NRXN3 | LRRTM3
NRXN1 | LRRTM3
NRXN3 | NLGN1
NRXN1 | NLGN1
AVN_P_cell
NC2_glial_NGF+
aCM2
AVN_P_cell
Ligand expressing cells
B
LGIC interactions (SAN)
Ligand | Receptor
LR mean
0.8
0.7
0.6
0.5
Glutamate_byGLS_and_SLC1A3 | GRIA3
Glutamate_byGLS_and_SLC1A3 | GRIK1
Glutamate_byGLS_and_SLC1A3 | GRIK2
SAN_P_cell
FB4_activated
SAN_P_cell
SAN_P_cell
Ligand expressing cells
C
LGIC interactions (AVN)
Ligand | Receptor
LR mean
0.8
0.7
0.6
0.5
0.4
0.3
0.2
Glutamate_byGLS_and_SLC1A2 | GRIA3
Glutamate_byGLS_and_SLC1A3 | GRIA3
Glutamate_byGLS_and_SLC1A2 | GRIK1
Glutamate_byGLS_and_SLC1A3 | GRIK1
Glutamate_byGLS_and_SLC1A2 | GRIK2
Glutamate_byGLS_and_SLC1A3 | GRIK2
AVN_P_cell
FB2
aCM2
LYVE1+IGF1+MP
FB5
AVN_P_cell
AVN_P_cell
Ligand expressing cells

### Supp Fig 14

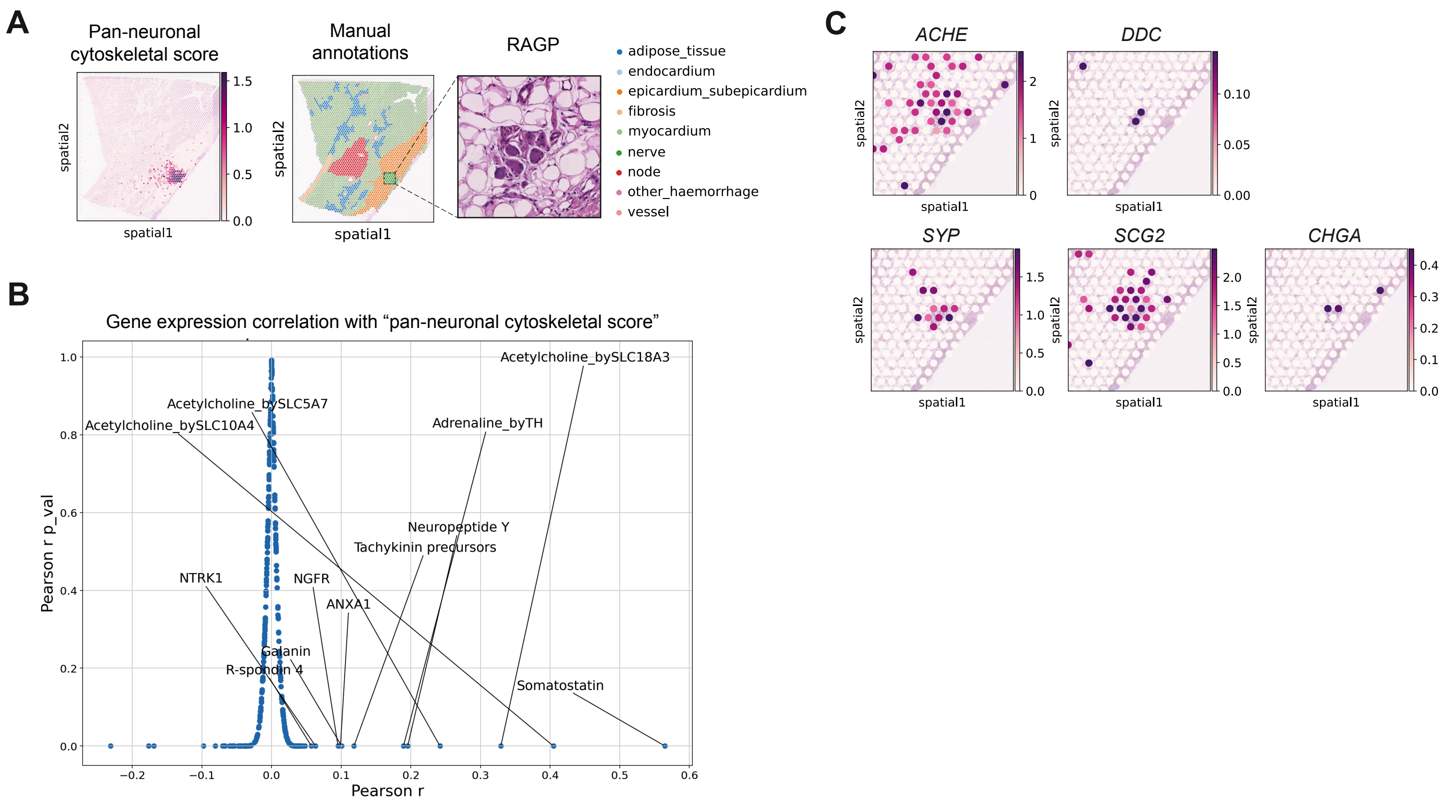
A
Pan-neuronal cytoskeletal score
Manual annotations
RAGP
spatial1
spatial2
adipose_tissue
endocardium
epicardium_subepicardium
fibrosis
myocardium
nerve
node
other_haemorrhage
vessel
B
Gene expression correlation with “pan-neuronal cytoskeletal score”
Pearson r p_val
Pearson r
Acetylcholine_bySLC18A3
Acetylcholine_bySLC5A7
Acetylcholine_bySLC10A4
Adrenaline_byTH
Neuropeptide Y
Tachykinin precursors
NTRK1
NGFR
ANXA1
Galanin
R-spondin 4
Somatostatin
C
ACHE
DDC
SYP
SCG2
CHGA

### Supp Fig 15

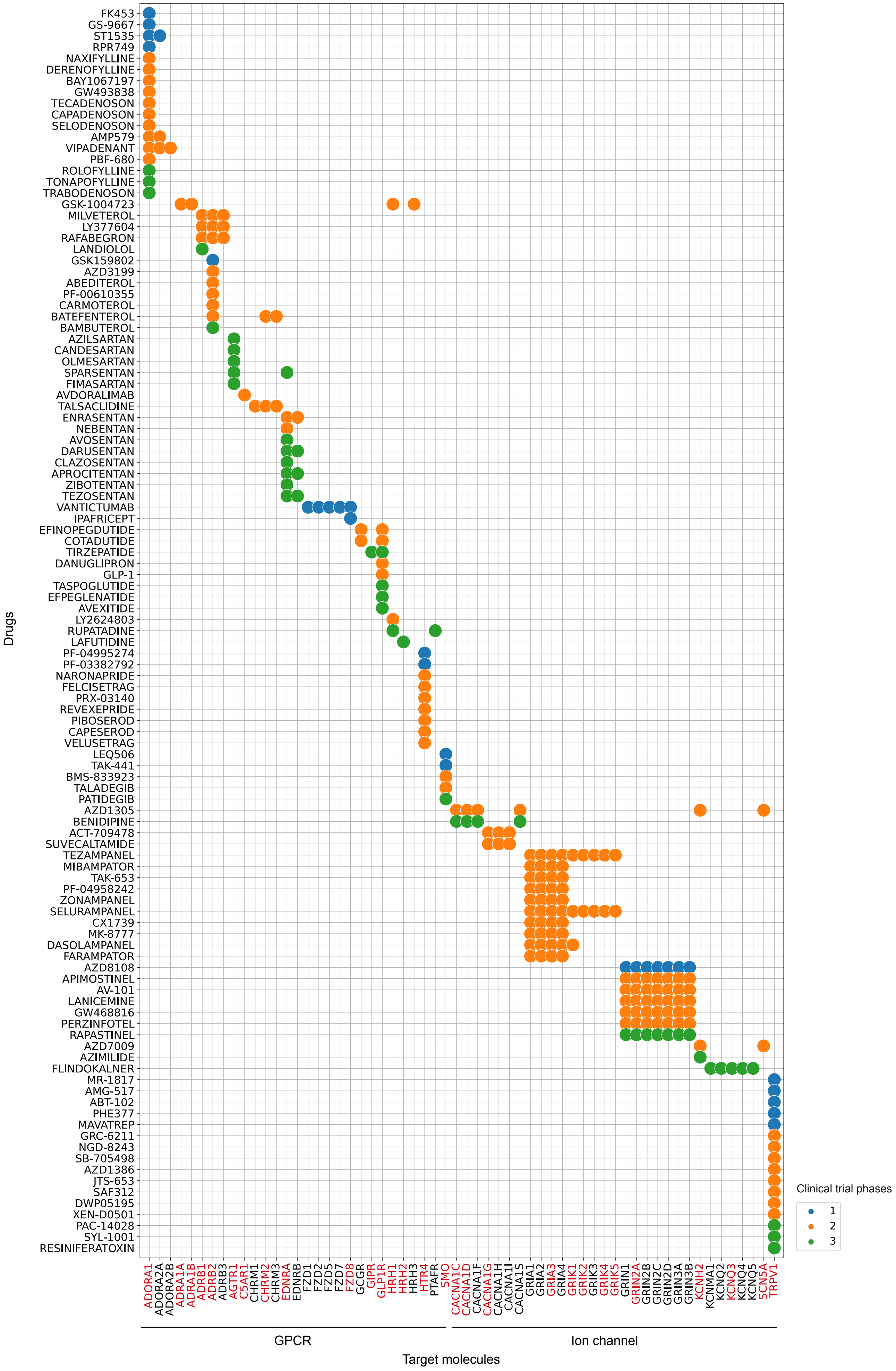
Drugs
FK453
GS-9667
ST1535
RPR749
NAXIFYLLINE
DERENOFYLLINE
BAY1067197
GW493838
TECADENOSON
CAPADENOSON
SELODENOSON
AMP579
VIPADENANT
PBF-680
ROLOFYLLINE
TONAPOFYLLINE
TRABODENOSON
GSK-1004723
MILVETEROL
LY377604
RAFABEGRON
LANDIOLOL
GSK159802
AZD3199
ABEDITEROL
PF-00610355
CARMOTEROL
BATEFENTEROL
BAMBUTEROL
AZILSARTAN
CANDESARTAN
OLMESARTAN
SPARSENTAN
FIMASARTAN
AVDORALIMAB
TALSACLIDINE
ENRASENTAN
NEBENTAN
AVOSENTAN
DARUSENTAN
CLAZOSENTAN
APROCITENTAN
ZIBOTENTAN
TEZOSENTAN
VANTICTUMAB
IPAFRICEPT
EFINOPEGDUTIDE
COTADUTIDE
TIRZEPATIDE
DANUGLIPRON
GLP-1
TASPOGLUTIDE
EFPEGLENATIDE
AVEXITIDE
LY2624803
RUPATADINE
LAFUTIDINE
PF-04995274
PF-03382792
NARONAPRIDE
FELCISETRAG
PRX-03140
REVEXEPRIDE
PIBOSEROD
CAPESEROD
VELUSETRAG
LEQ506
TAK-441
BMS-833923
TALADEGIB
PATIDEGIB
AZD1305
BENIDIPINE
ACT-709478
SUVECALTAMIDE
TEZAMPANEL
MIBAMPATOR
TAK-653
PF-04958242
ZONAMPANEL
SELURAMPANEL
CX1739
MK-8777
DASOLAMPANEL
FARAMPATOR
AZD8108
APIMOSTINEL
AV-101
LANICEMINE
GW468816
PERZINFOTEL
RAPASTINEL
AZD7009
AZIMILIDE
FLINDOKALNER
MR-1817
AMG-517
ABT-102
PHE377
MAVATREP
GRC-6211
NGD-8243
SB-705498
AZD1386
JTS-653
SAF312
DWP05195
XEN-D0501
PAC-14028
SYL-1001
RESINIFERATOXIN
ADORA1
ADORA2A
ADORA2B
ADRA1A
ADRA1B
ADRB1
ADRB2
ADRB3
AGTR1
C5AR1
CHRM1
CHRM2
CHRM3
EDNRA
EDNRB
FZD1
FZD2
FZD5
FZD7
FZD8
GCGR
GIPR
GLP1R
HRH1
HRH2
HRH3
HTR4
PTAFR
SMO
CACNA1C
CACNA1D
CACNA1F
CACNA1G
CACNA1H
CACNA1I
CACNA1S
GRIA1
GRIA2
GRIA3
GRIA4
GRIK1
GRIK2
GRIK3
GRIK4
GRIK5
GRIN1
GRIN2A
GRIN2B
GRIN2C
GRIN2D
GRIN3A
GRIN3B
KCNH2
KCNMA1
KCNQ2
KCNQ3
KCNQ4
KCNQ5
SCN5A
TRPV1
GPCR
Ion channel
Target molecules
Clinical trial phases
1
2
3

Supp Fig 16

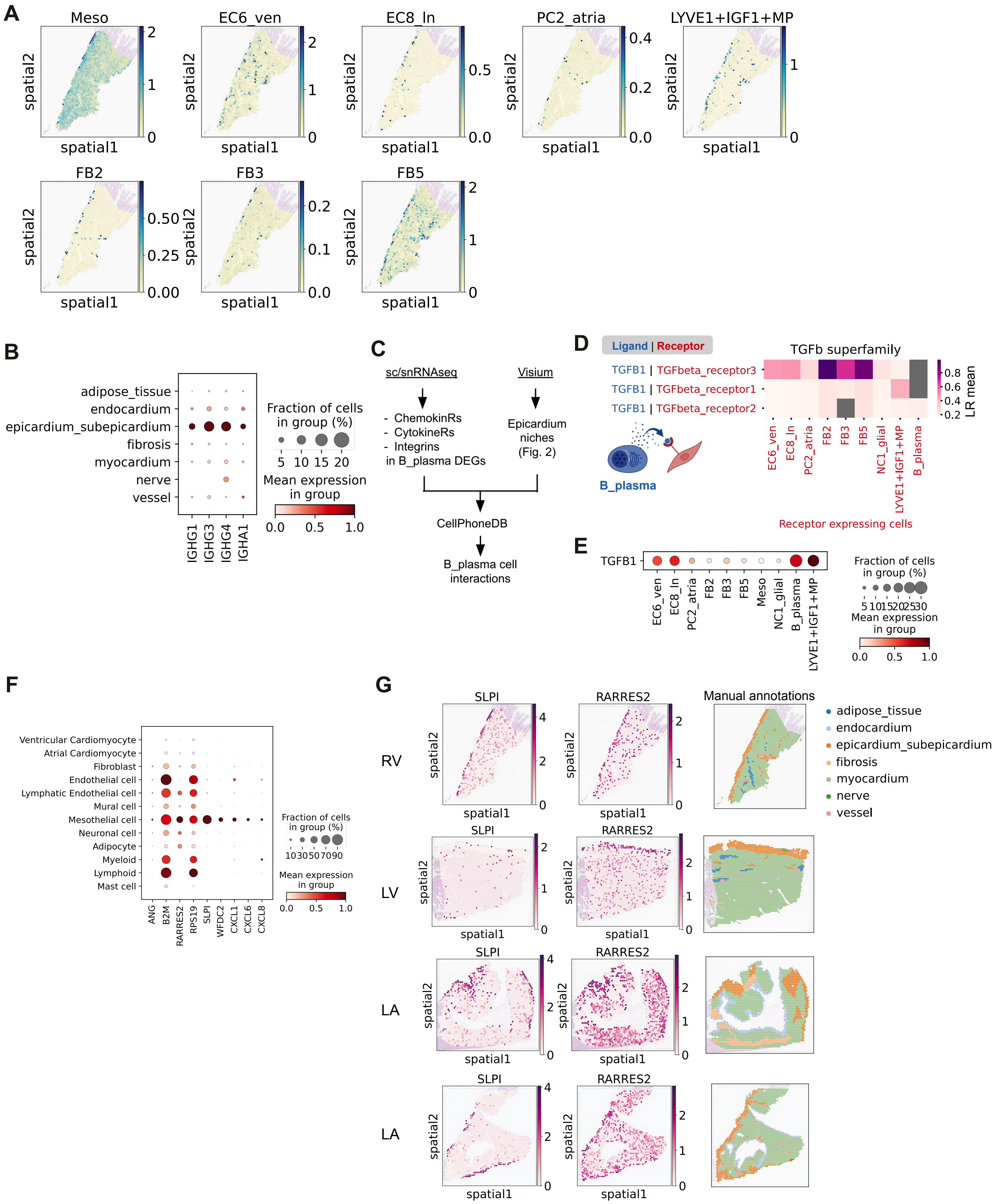
A
Meso
EC6_ven
EC8_ln
PC2_atria
LYVE1+IGF1+MP
FB2
FB3
FB5
spatial1
spatial2
B
adipose_tissue
endocardium
epicardium_subepicardium
fibrosis
myocardium
nerve
vessel
IGHG1
IGHG3
IGHG4
IGHA1
Fraction of cells in group (%)
5 10 15 20
Mean expression in group
0.0 0.5 1.0
C
sc/snRNAseq
Visium
- ChemokinRs
- CytokineRs
- Integrins
in B_plasma DEGs
Epicardium niches (Fig. 2)
CellPhoneDB
B_plasma cell interactions
D
Ligand | Receptor
TGFb superfamily
TGFB1 | TGFbeta_receptor3
TGFB1 | TGFbeta_receptor1
TGFB1 | TGFbeta_receptor2
B_plasma
EC6_ven
EC8_ln
PC2_atria
FB2
FB3
FB5
NC1_glial
LYVE1+IGF1+MP
B_plasma
LR mean
Receptor expressing cells
E
TGFB1
EC6_ven
EC8_ln
PC2_atria
FB2
FB3
FB5
Meso
NC1_glial
B_plasma
LYVE1+IGF1+MP
Fraction of cells in group (%)
5 10 15 20 25 30
Mean expression in group
0.0 0.5 1.0
F
Ventricular Cardiomyocyte
Atrial Cardiomyocyte
Fibroblast
Endothelial cell
Lymphatic Endothelial cell
Mural cell
Mesothelial cell
Neuronal cell
Adipocyte
Myeloid
Lymphoid
Mast cell
ANG
B2M
RARRES2
RPS19
SLPI
WFDC2
CXCL1
CXCL6
CXCL8
Fraction of cells in group (%)
10 30 50 70 90
Mean expression in group
0.0 0.5 1.0
G
RV
LV
LA
LA
SLPI
RARRES2
Manual annotations
adipose_tissue
endocardium
epicardium_subepicardium
fibrosis
myocardium
nerve
vessel

Supp Fig 17

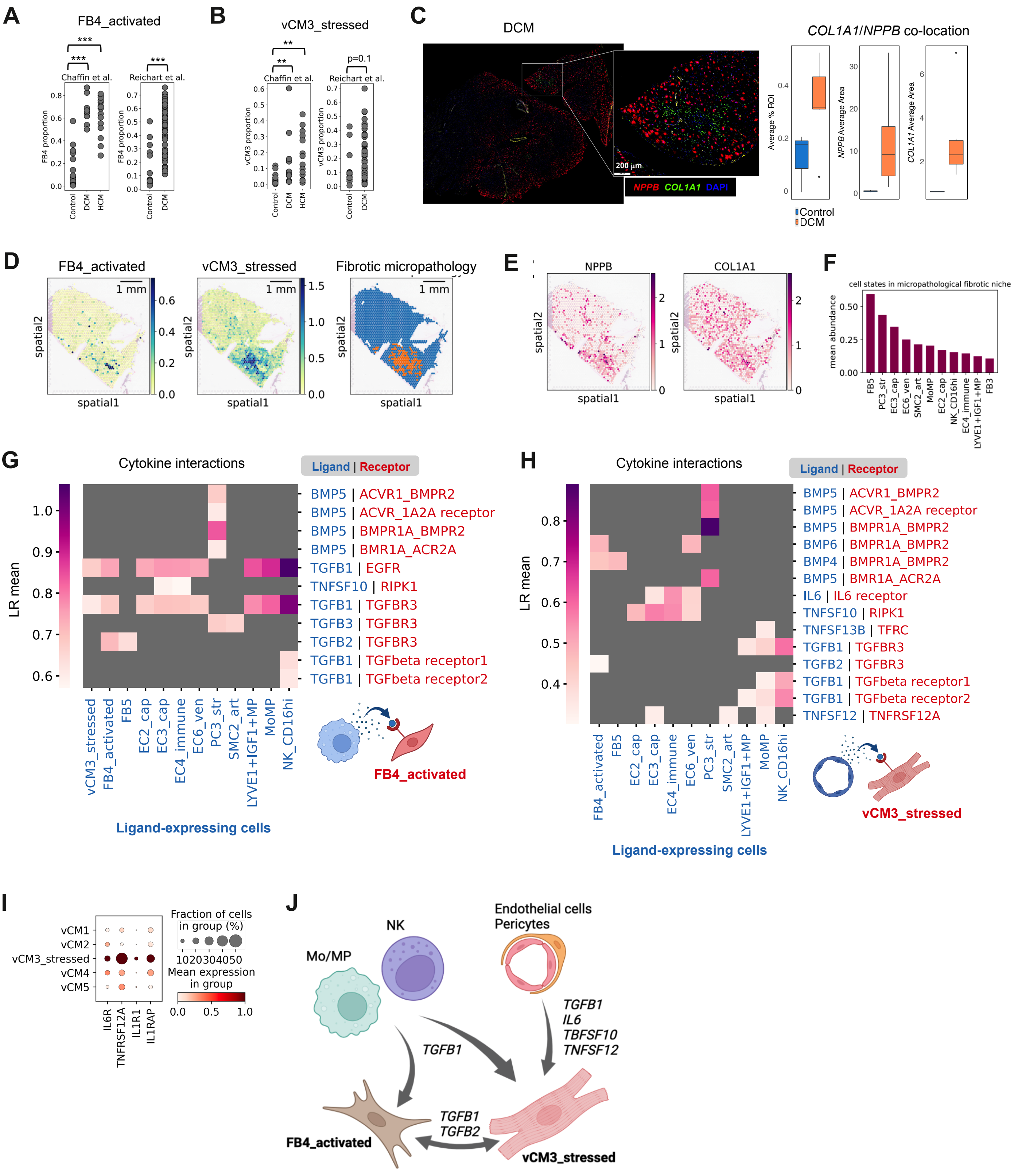
A
FB4_activated
Chaffin et al.
Reichart et al.
FB4 proportion
Control
DCM
HCM
B
vCM3_stressed
Chaffin et al.
Reichart et al.
p=0.1
vCM3 proportion
C
DCM
200 µm
NPPB COL1A1 DAPI
COL1A1/NPPB co-location
Average % ROI
NPPB Average Area
COL1A1 Average Area
Control
DCM
D
FB4_activated
vCM3_stressed
Fibrotic micropathology
1 mm
spatial1
spatial2
E
NPPB
COL1A1
F
cell states in micropathological fibrotic niche
mean abundance
FB5
PC3_str
EC3_cap
EC6_ven
SMC2_art
MoMP
EC2_cap
NK_CD16hi
EC4_immune
LYVE1+IGF1+MP
FB3
G
Cytokine interactions
Ligand | Receptor
LR mean
BMP5 | ACVR1_BMPR2
BMP5 | ACVR_1A2A receptor
BMP5 | BMPR1A_BMPR2
BMP5 | BMR1A_ACR2A
TGFB1 | EGFR
TNFSF10 | RIPK1
TGFB1 | TGFBR3
TGFB3 | TGFBR3
TGFB2 | TGFBR3
TGFB1 | TGFbeta receptor1
TGFB1 | TGFbeta receptor2
vCM3_stressed
FB4_activated
FB5
EC2_cap
EC3_cap
EC4_immune
EC6_ven
PC3_str
SMC2_art
LYVE1+IGF1+MP
MoMP
NK_CD16hi
Ligand-expressing cells
FB4_activated
H
Cytokine interactions
Ligand | Receptor
BMP5 | ACVR1_BMPR2
BMP5 | ACVR_1A2A receptor
BMP5 | BMPR1A_BMPR2
BMP6 | BMPR1A_BMPR2
BMP4 | BMPR1A_BMPR2
BMP5 | BMR1A_ACR2A
IL6 | IL6 receptor
TNFSF10 | RIPK1
TNFSF13B | TFRC
TGFB1 | TGFBR3
TGFB2 | TGFBR3
TGFB1 | TGFbeta receptor1
TGFB1 | TGFbeta receptor2
TNFSF12 | TNFRSF12A
Ligand-expressing cells
vCM3_stressed
I
vCM1
vCM2
vCM3_stressed
vCM4
vCM5
IL6R
TNFRSF12A
IL1R1
IL1RAP
Fraction of cells in group (%)
1020304050
Mean expression in group
J
Mo/MP
NK
Endothelial cells
Pericytes
TGFB1
TGFB1
IL6
TBFSF10
TNFSF12
TGFB1
TGFB2
FB4_activated
vCM3_stressed
